# The cleaved C-terminal domain of GmERS1-1 modulates cell cycle via GmACR1

**DOI:** 10.64898/2026.10.07.757225

**Authors:** Kai Chen, Yongbin Zhou, Wensi Tang, Daoping Wang, Yanming Bai, Chunxiao Wang, Weiwei Li, Xin Li, Dongdong Guo, Meishan Zhang, Jiji Yan, Jia Guo, Yan Zhao, Baoxue Shan, Jun Chen, Zhaoshi Xu, Yubing He, Junkui Ma, Qingxin Song, Haiyan Li, Lijuan Qiu, Haiyang Wang, Ming Chen, Youzhi Ma

## Abstract

Ethylene signaling orchestrates plant development and stress adaptation, yet how post-translational modifications of its receptors coordinate growth regulation remains unresolved. Here, we uncover a proteolytic processing mechanism of the soybean ethylene receptor GmERS1-1 that directly gates cell cycle progression. The *GmERS1-1* mutants exhibit elevated yield under field conditions, whereas overexpression suppresses growth and hypersensitizes plants to ethylene. Ethylene triggers nuclear translocation of C-terminal domain (GmERS1-1-CEND), which is indispensable for receptor function. Mechanistically, GmERS1-1-CEND scaffolds a trimeric complex with GmACR1 and GmCDK7, enforcing G2/M phase arrest, thereby decoupling cell division from canonical ethylene transcriptional responses. Strikingly, ethylene-induced cell cycle blockade during apple fruit maturation mirrors this pathway, suggesting deep conservation across angiosperms. Our work defines a receptor processing-dependent signaling axis that directly interlinks phytohormone perception with cell cycle control, proposing tunable targets (e.g., GmERS1-1 cleavage efficiency) for engineering crop architecture and productivity.

**In brief:** The cleaved C-terminal fragment of ethylene receptor GmERS1-1 enters into nucleus to inhibit cell cycle via interaction with GmACR1 in soybean.

**Highlights:**

1. Mutating ethylene receptor *GmERS1-1* significantly increases soybean yield in field.
2. The cleaved C-terminal fragment of GmERS1-1 enters into the nucleus induced by ethylene and independent of CTR1.
3. The C-terminal fragment has similar function to the full-length ethylene receptor GmERS1-1 to induce ethylene reaction in soybean and *Arabidopsis*.
4. The C-terminal fragment of GmERS1-1 interacts with soybean GmACR1 to inhibit cell division.

## INTRODUCTION

Soybean is an important bulk cash crop, which provides a lot of plant protein and oil for human beings. The better the root system development of soybeans, the less they are affected by water stress and the stronger their resistance to stress. Therefore, it is of great practical significance to deeply explore the regulatory mechanisms of soybean root development under adverse conditions, in order to alleviate the threat of drought or waterlogging and ensure the safe production of soybeans. The growth and development of plant roots are inseparable from cell division and differentiation. Whether it is the initiation and division of root primordia, or the formation and activation of meristem tissue, both involve two alternating cycles of cell division and cell growth in the cell cycle (*1*). Therefore, studying the mechanism of cell cycle regulation not only elucidates the molecular mechanisms of plant growth and development, but also helps to elucidate the mechanism of stress on crop growth.

Plant growth, which depends on cell division as one of the critical processes, is regulated by several hormones. Ethylene has long been recognized as a growth inhibitor (*2*). For example, the excessive production of ethylene leads to the short fruit phenotype of cucumbers (*3*). Ethylene represses the growth promoting influence of cytokinin, whether endogenous or applied (*4*). During osmotic stress, a cell cycle arrest was observed in *Arabidopsis thaliana* leaves in parallel with an increase in 1-aminocyclopropane-1-carboxylate (ACC) levels and activation of ethylene signaling (*5*, *6*). However, the molecular mechanism by which ethylene inhibits cell division is still unclear.

The classic ethylene signaling pathway starts from the interaction between ethylene receptors and downstream Raf-like protein CTR1, activating CTR1, which in turn phosphorylates the downstream ethylene signaling component EIN2 and inhibits EIN2 from transmitting ethylene signals. When receiving ethylene, the receptor is unable to activate CTR1, causing EIN2 to undergo functionally protein cleavage. The carboxyl end of EIN2 enters the nucleus to transmit ethylene signals, activating downstream transcription factor EIN3 to generate ethylene responsive gene expression and inducing ethylene response (*7–10*). However, the study of ethylene signal pathway in soybean has not been carried out in depth, so it is particularly important to analyze the ethylene signal transduction mechanism in soybean for further understanding the growth and development process of soybean.

There are five ethylene receptor proteins in CTR1-mediated traditional ethylene signal transduction pathway: ETR1, ERS1, ETR2, ERS2 and EIN4 (*11*). They have some functional redundancy and are negative regulators of ethylene signaling. In this study, we found that the carboxyl terminal (CEND) of ethylene receptor GmERS1-1 can output receptor signals in the absence of CTR1, inhibiting the growth of plants, which is dependent on the entry of CEND into the nucleus. We obtained an arsenate reductase GmACR1 that interacts with GmERS1-1 through screening from a yeast library. Plant arsenate reductase belongs to the ACR family. There is a CDC25 catalytic domain (RHOD) homologous protein AtASR (also arsenate reductase) in *Arabidopsis thaliana*, and studies have shown that this protein has the activity of reducing arsenic acid. AtASR also exhibits phosphatase activity, indicating its CDC25-like function (*12*). All of these suggest that GmACR1 may have functions similar to CDC25. And GmERS1-1 interacts with GmACR1 to cascade the ethylene signaling pathway and cell cycle regulation.

The eukaryotic cell cycle is composed of four well-defined phases, G1, S, G2 and mitosis (M). Progression through the cell cycle is strictly regulated, especially during transitions from the G1 phase to S phase, from G2 phase to M and when leaving M to return to G1. Cell cycle checkpoints are control mechanisms that ensure the accurate replication of the genome to pass genomic integrity onto the daughter cell. In actuality, one study found that ethylene perception is involved in the arrest of stamen development in female cucumber flowers through the induction of DNA damage (*13*). 17K protein, encoded by barley yellow dwarf viruses and related poleroviruses, delays G_2_/M transition and disrupts mitosis in both host (barley) and nonhost (fission yeast, Arabidopsis thaliana, and tobacco) cells through interrupting the function of Wee1-Cdc25-CDKA/Cdc2 via direct protein-protein interactions and alteration of CDKA/Cdc2 phosphorylation (*14*). Therefore, ethylene may inhibit plant growth through cell cycle regulation by GmERS1-1-GmACR1 module.

The presence of these pathways increases the complexity of ethylene signal transduction pathways, suggesting that the transition from receiving ethylene signals to generating specific ethylene responses in plants may be a complex and sophisticated process in which different pathways act synergistically.

## Results

### The mutation of *GmERS1-1* boosts soybean yield in field

Overexpression of *AtERS1* could enhance sensitivity to ethylene, manifested as an enhanced the triple response phenotype in *Arabidopsis* (*15*, *16*). We hypothesized that ERS like ethylene receptors may have similar functions in soybean because both of soybean and *Arabidopsis* belong to dicotyledonous plants. Sequence alignment revealed that soybean contains 11 ethylene receptor genes. Cluster analysis on all ethylene receptor families in soybean and *Arabidopsis* showed that both of *GmERS1-1* (*Glyma.03G216700.1*) and *AtERS1* (*AT2G40940.1*) belong to the same branch in the second subfamily **(Fig. 1A)**. Gene expression profiling analysis showed that *GmERS1-1* had the highest expression level in almost all tissues and organs of soybean among all ethylene receptor genes **(Fig. 1B)**. Gene expression profiling of *GmERS1-1* were also confirmed by promoter activity analysis using GUS as reporter gene in *Arabidopsis* **(Fig. 1C)**. In addition, expression analysis using the soybean root transient expression system showed that *GmERS1-1* was induced in soybean roots by ethylene treatment **(Fig. 1, D and E)**. In summary, *GmERS1-1* were induced by ethylene and expressed throughout almost tissues of the plant, indicating that GmERS1-1 as important member of ethylene receptor gene family, plays a key role in the soybean ethylene signaling pathway.410.

**Figure 1.**
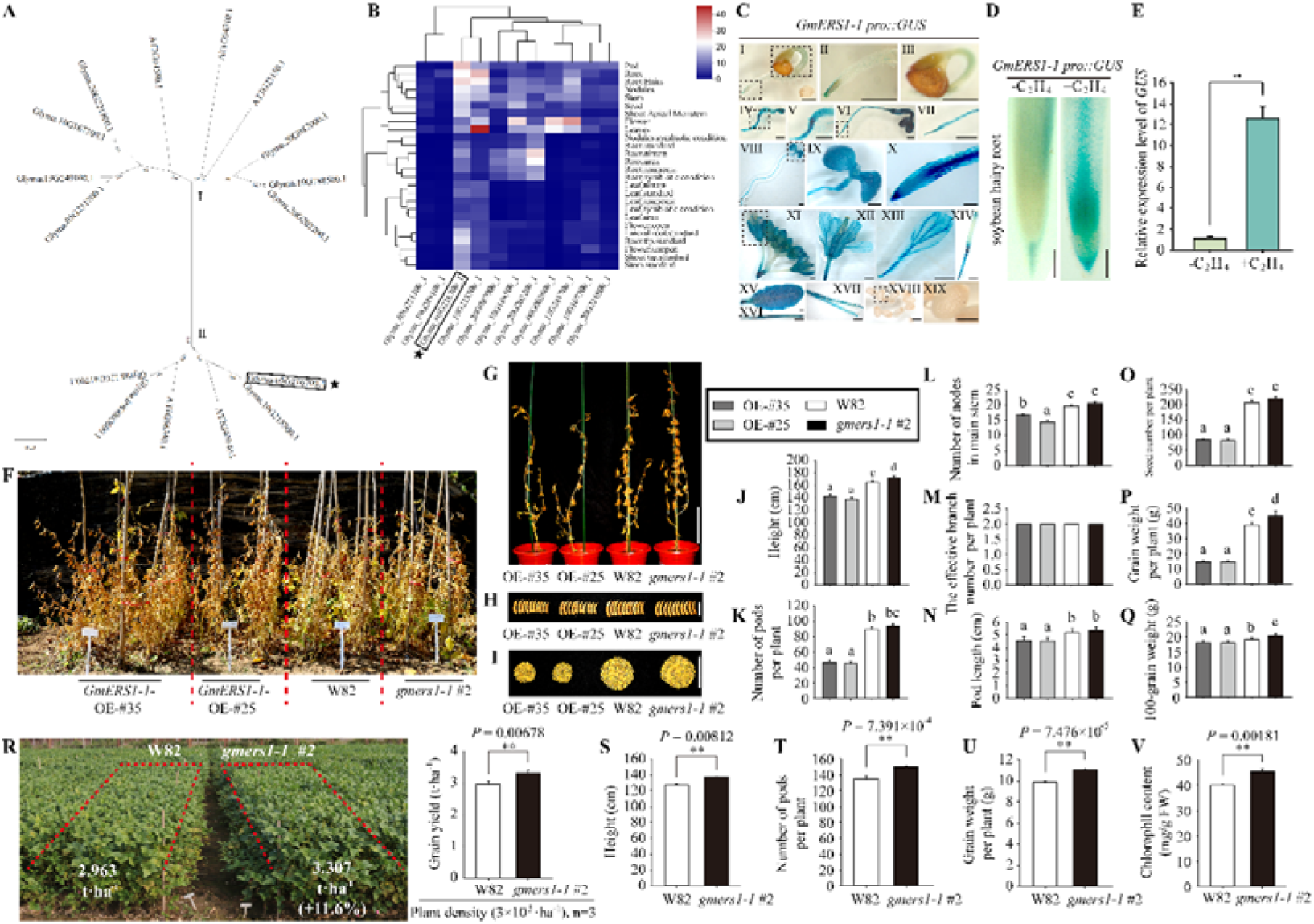
The overexpression of *GmERS1-1* inhibits the growth of soybeans. **(A-E)** Analysis of the expression pattern of soybean ethylene receptor *GmERS1-1*. **(A)** Construct evolutionary trees for the ethylene receptor families of soybeans (11) and *Arabidopsis* (5) by using DNAMAN software. The results showed that GmERS1-1 (Glyma.03G216700.1) belongs to the second subfamily. GmERS1-1 was marked with a black box and an asterisk. **(B)** Expression profiles of 11 soybean ethylene receptor genes. *GmERS1-1* (Glyma.03G216700) was circled by red box. **(C) Ⅰ-XIX** are tissue specific expression of *GmERS1-1* (GUS staining). We obtained stable transformed Arabidopsis lines of *GmERS1-1 pro::GUS* using floral dip method, and then observed the expression of *GUS* reporter gene throughout the entire plant life cycle. **(Ⅰ)** The seeds germinate for three days. **(Ⅱ)** The root tip in **Ⅰ** was shown. **(Ⅲ)** The cotyledons that are about to break open the seed coat. Like **Ⅱ**, **Ⅲ** is a zoomed in view of a specific area in **Ⅰ**. **(Ⅳ)** Seedlings that have germinated for 5 days. **(Ⅴ)** The transition zone at the junction of the hypocotyl and root. **(Ⅵ)** Seedlings that have germinated for 6 days. **(Ⅶ)** The root tip in **Ⅵ** was shown. **(Ⅷ)** Seedlings that have germinated for 7 days. **(Ⅸ)** Enlarged image of fully unfolded cotyledons in **Ⅷ**. **(Ⅹ)** The root tip in **Ⅷ** was shown. **(Ⅺ)** Inflorescence. **(Ⅻ)** Bud. **(XIII)** Petal. **(XIV)** Pistil. **(XV)** Leaf. **(XVI)** Silique. **(XVII)** Cracked siliqua. **(XVIII)** Young and tender seeds. **(XIX)** Localized enlargement of **XVIII**. Scale bar=1mm. **(D)** After obtaining the root of soybean with *GUS* reporter gene, it was subjected to ethylene treatment. The results showed that *GmERS1-1* can respond to ethylene signals. **(E)** *GUS* transcription with C_2_H_4_ treatment for 30 minutes. Detect at least 30 replicates per organism. The error line represents SE. Statistical analysis was conducted using Student’s *t*-test. Asterisks indicate comparison with the control group (**, *P* < 0.01). **(F)** Field planting experiment of soybean lines. From left to right are: overexpression line #35 (OE-#35) of *GmERS1-1*, overexpression line #25 (OE-#25) of *GmERS1-1*, wild-type Williams 82 (W82), and mutant of *GmERS1-1* (*gmers1-1* #2). They are separated by a red dashed line. Repeat planting two rows for each line. **(G)** G is the image of each single line in F. Bar=20cm. **(H)** This figure shows the top 10 pods of a single line in F from top to bottom. Bar=5cm. **(I)** This picture shows all the seeds of a single soybean plant in F. Bar=10cm. **(J-Q)** Statistical analysis of field data. **(J)** Plant height of different materials. **(K)** Number of pods per plant for different materials. **(L)** The number of main stem nodes of different materials. **(M)** The effective branching number of different materials. **(N)** The pod length of different materials. **(O)** The number of grains per plant for different materials. **(P)** The individual grain weight of different materials. **(Q)** The one hundred-grain weight of different materials. The columns of different colors represent different lines, and the legend is shown in the black box. Values are means ± SE (n = 30). Letters indicate significant differences by two-way ANOVA analysis. **(R-V)** Field phenotype comparison experiment between wild-type W82 and mutant *gmers1-1* #2. **(R)** The on-site images of the field phenotype comparison experiment were taken during the soybean grain filling stage in Shunyi District, Beijing. The lower part of the figure indicates the grain yield data, and the right side is a bar chart of the grain yield data. **(S)** shows the plant height. **(T)** shows the number of pods per plant. **(U)** shows the yield per plant. **(V)** displays the chlorophyll content (\*\**P* < 0.01; two-tailed Student’s *t*-test; data are mean ± SE; n=3).

Next, we demonstrate the function of GmERS1-1 in soybeans through a series of experiments, particularly its impact on soybean field yield, which had not been analyzed in model plants. Firstly, we verified the function of *GmERS1-1* in soybean using the method of the transient transformation of soybean hairy roots **(fig. S1)**. The result showed that overexpression of *GmERS1-1* not only reduced soybean plant height **(fig. S1, A and D)**, but also inhibited growth of root and pods **(fig. S1, B, C, E and F)**. Secondly, we obtained the stably transgenic soybean *GmERS1-1 OE* (overexpressing) lines by the agrobacterium infecting cotyledons method **(fig. S2)**. After the molecular detection of the transformed soybeans **(fig. S2, A to C)** and the phenotypic investigation of T_1_ generation plants **(fig. S2, D to H)**, we selected T_4_ generation homozygous OE lines #25 and #35 of *GmERS1-1* for the subsequent experimental. We found that the growth of *GmERS1-1* OE lines was weaker than that of wild type in the absence of ethylene **(fig. S2, C, F to H)**. In addition, we obtained M5 generation homozygous mutants *gmers1-1* #2 **(fig. S3, A and B)** with consistent phenotypes of the EMS mutation described by iSoybean (*17*). Finally, we conducted field experiments of transgenic soybean and mutants **(Fig. 1, F to Q)**. Certainly, what exceeded our expectations the most was that *gmers1-1* #2 would result in better soybean growth than the wild type, as evidenced by increased plant height, number of main stem nodes, increased number of pods per plant, increased pod length, increased number of grains per plant, increased grain weight per plant, and ultimately increased yield **(Fig. 1, F to Q)**. In standard soybean plot experiment (plots area 20 m^2^) using medium planting densities (3×10^5^·ha), the yield of *gmers1-1* (3.307 t·ha^-1^) increased 11.6% compared with wild type W82 (2.963 t·ha^-1^) **(Fig. 1R)**, which suggesting that mutant *gmers1-1* had great potential to enhance grain yield of soybean. Notably, in addition to excellent performance in indicators such as plant height (**Fig. 1S**), number of pods per plant **(Fig. 1T)**, and grain weight per plant **(Fig. 1U)**, *gmers1-1 #2* has higher chlorophyll content **(Fig. 1V)** and better green retention during the grain filling stage **(Fig. 1R)**, suggesting that *gmers1-1* mutant have higher yield comparted to W82 under normal condition. To further verify the reliability of field trial results, we conducted field cultivation experiments in Fenyang, Shanxi Province in 2025 **(fig. S4)**. The field trial data from Fenyang, Shanxi (2025) revealed that under a planting density of 2×10⁵ plants per hectare, the actual grain yield of the *gmers1-1* #2 mutant plot reached 2,747 kg/ha, representing a 13.97% yield increase compared with the wild-type W82 (2,410 kg/ha), with a significant difference in grain yield per hectare between the two genotypes (*P*=0.014663, n=3) **(fig. S4, A and B)**. Measurements of individual plant agronomic traits demonstrated that the *gmers1-1* #2 mutant exhibited significantly greater plant height (*P*=0.047352) and higher leaf chlorophyll content (*P*=0.033921) than W82, while no statistically significant differences were detected between the two genotypes for main stem node number, pod number per plant, lowest pod height, and number of effective branches **(fig. S4, C to H)**. Collectively, these results further confirm that mutation of *GmERS1-1* markedly improves soybean grain yield per unit area by enhancing plant height and leaf photosynthetic pigment levels, without altering agronomic traits including main stem node count, branching architecture and pod-setting characteristics.

**Figure 2.**
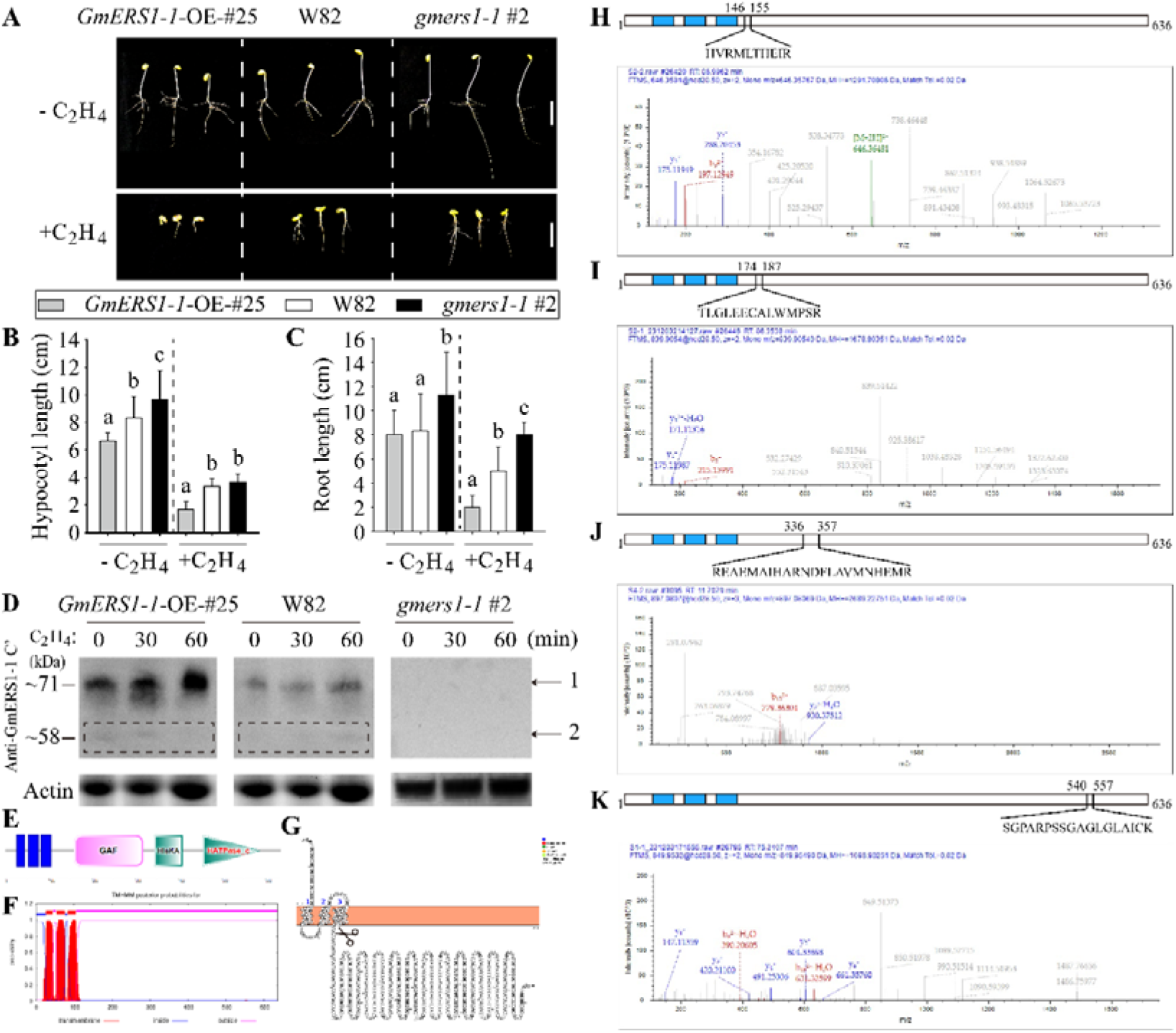
The overexpression of *GmERS1-1* enhances soybean sensitivity to ethylene. **(A)** The phenotype of “triple reaction” of the overexpression line #25 (OE-#25) of *GmERS1-1*, wild-type Williams 82 (W82), and mutant of *GmERS1-1* (*gmers1-1* #2), with no ethylene treatment on the top line and ethylene treatment on the row below. These white dashed lines separate these three lines. Each group contains three biological replicates. Bar=5cm. **(B-C)** Statistical analysis of hypocotyl length (B) and root length (C) in A. The columns of different colors represent different lines, and the legend is shown in the black box. Data statistics conducted ANOVA-DUNCAN’s multiple range tests at P<0.05, error bars report SE of the mean, n=3. **(D)** Using Western-blot to detect the expression level of GmERS1-1 protein with Anti-GmERS1-1 C’ mouse monoclonal antibody during the ethylene treatment process. 1 represents the full-length protein of GmERS1-1, and 2 represents the unknown fragment of GmERS1-1. The dashed box marks the additional protein bands that appear. Use Actin protein as a control to ensure consistent sample loading. **(E-G)** Analyze the protein structure of GmERS1-1 using multiple prediction methods. **(E)** SMART (Simple modular architecture research tool) (http://smart.embl-heidelberg.de/) analysis of GmERS1-1. The blue square represents the transmembrane domain. **(F)** DeepTMHMM - 1.0 (https://services.healthtech.dtu.dk/services/DeepTMHMM-1.0/) analysis of GmERS1-1. The red area represents the transmembrane domain. **(G)** Protter (interactive protein feature visualization) (http://wlab.ethz.ch/protter/#) analysis of GmERS1-1. Scissors indicate possible cutting points. **(H-K)** Mass spectrometry analysis was performed on the unexpected protein in A (the part within the dashed box), and a total of four peptides were detected. The secondary structure diagram of GmERS1-1 is marked above each mass spectrum, and the blue rectangle in the secondary structure diagram represents the transmembrane domain. The identified peptide segment positions are marked with black solid lines. The red line in the mass spectrum represents the target peptide segment. **(H)** The peptide segment detected in H is HVRMLTHEIR, located from position 146 to position 155 of the amino acid sequence. **(I)** The peptide segment detected in I is TLGLEECALWMPSR, located from position 174 to position 187 of the amino acid sequence. **(J)** The peptide segment detected in J is REAEMAIHARNDFLAVMNHEMR, located from position 336 to position 357 of the amino acid sequence. **(K)** The peptide segment detected in K is SGPARPSSGAGLGLAICK, located from position 540 to position 557 of the amino acid sequence.

Structural prediction showed that soybean GmERS1-1 and *Arabidopsis* AtERS1 have similar structures, suggesting that they may have similar functions and the Anti-GmERS1-1 C’ antibody (antibody for GmERS1-1 C-terminal) can be used to detect AtERS1-CEND (free C-terminal of AtERS1 in *Arabidopsis*) **(fig. S5A)**. Therefore, in order to analyze the conservation of GmERS1-1 functions in *Arabidopsis*, we obtained some genetic materials including *Arabidopsis AtERS1* mutants *aters1* **(fig. S5, B and C)** and overexpressing *GmERS1-1* in *Arabidopsis* **(Figure. S6)**. Phenotypic analysis results showed that the growth of *Arabidopsis* with overexpressing *GmERS1-1* was weaker than that of wild type *Arabidopsis*, and *Arabidopsis* mutants *aters1* were better than wild type *Arabidopsis* including the leaf number, fresh weight of aboveground parts, number of pods, effective number of branches and seed yield **(fig. S6, A to O)**. Therefore, the function analysis results of *GmERS1-1* in soybean and *Arabidopsis* indicated that mutations of *GmERS1-1* promoted plant growth and increased yield, while overexpression of the *GmERS1-1* gene inhibited plant growth and reduced yield.

### The overexpression of *GmERS1-1* enhances soybean sensitivity to ethylene

Normally, high concentration ethylene acts as a growth inhibitor, mainly by promoting leaf senescence, limiting root growth, and so on (*18*, *19*). In this research, the leaves of the *GmERS1-1* overexpression lines were prematurely aged and shed **(Fig. 1F)**, whereas delaying aging characteristic of the *gmers1-1* mutants are more pronounced compared to wild type soybean **(Fig. 1R and fig. S4A)**. We speculate that overexpression of *GmERS1-1* enhances soybean sensitivity to ethylene. Generally, the response of plants to ethylene can be analyzed through “the triple reactions” includes inhibiting stem elongation growth, promoting lateral thickening of the stem or hypocotyl, and causing lateral growth of the stem (*20*). We injected 10 ppm ethylene gas into 20 L sealed box to treat etiolation seedlings (*21*) for testing the triple responses in soybean **(Fig. 2A)**. The results showed that *GmERS1-1* OE #25 had shorter hypocotyls and roots, whereas mutant *gmers1-1* had longer hypocotyls and roots compared to W82 after ethylene treatment **(Fig. 2, B to C)**, which indicated that overexpression of *GmERS1-1* enhanced ethylene sensitivity in soybean.

The results of the western blot using the Anti-GmERS1-1 C’ antibody showed that the level of protein GmERS1-1 in the overexpression lines was higher than that in the wild type. Surprisingly, we found small molecular weight band (∼58 kDa) below the full-length protein band (∼71 kDa) in transgenic and wild type soybean, whereas nothing in mutant *gmers1-1* **(Fig. 2D)**. The structure prediction shows that there are three transmembrane domains in the N-terminus of GmERS1-1, while the C-terminus is the free terminus **(Fig. 2, E to G)**. Because the Anti-GmERS1-1 C’ antibody is a monoclonal antibody specific for the C-terminus of GmERS1-1, we hypothesized that the C-terminus of GmERS1-1 could somehow dissociate from membrane, causing us to detect small molecular weight C-terminus of GmERS1-1 **(Fig. 2D)**. We further used the mass spectrometry to perform amino acid sequence for small molecular weights bands (∼58 kDa), and the results showed that the detected small peptides were concentrated at the C-terminus of GmERS1-1 **(Fig. 2, H to K)**. Therefore, we believe that the C-terminus of GmERS1-1 can dissociate from the intact GmERS1-1 located on the membrane under certain circumstances.

### Ethylene can induce the C-terminus of GmERS1-1 to enter the nucleus in soybean and *Arabidopsis*

As shown in **Fig. 1, C and D**, *GmERS1-1* expressed in many tissues such as root, hypocotyl, stem, cotyledons, leaves, flowers, cracked fruit pods in *Arabidopsis*. Because *GmERS1-1* highly expressed in roots, we detected the subcellular localization of GmERS1-1 protein in soybean W82 roots using the immunofluorescence method with Anti-GmERS1-1 C’ antibody **(Fig. 3A)**. We found that *GmERS1-1* highly expressed in the meristematic and elongation zones of soybean root tips, and GmERS1-1 was observed to be localized not only on the ER but also in the nucleus of the meristematic and elongation zones **(Fig. 3B)**. Due to the presence of three transmembrane domains at the N-terminus of GmERS1-1, we speculate that the whole GmERS1-1 protein is difficult to directly enter the nucleus and the free C-terminus GmERS1-1 in the cytoplasm might dissociate and enter into nucleus. To prove the speculation, we fused GFP with the N-terminus (*GmERS1-1 pro:GFP-GmERS1-1*) and C-terminus (*GmERS1-1 pro:GmERS1-1-GFP*) of GmERS1-1 to construct protein localization vectors and transformed soybean protoplasts, respectively **(Fig. 3, C to F)**. The results showed that when the GFP tag was fused with the N-terminus of GmERS1-1, the green fluorescence signal was not present in the nucleus with or without the presence of ethylene, only was located in the ER **(Fig. 3, C and D)**. When the GFP tag was fused with the C-terminus of GmERS1-1, we can observe GFP signals in the nucleus regardless of ethylene treatment, and ethylene application enhanced the fluorescence signal in the nucleus and weakened the signal in the ER **(Fig. 3, E and F)**.

**Figure 3.**
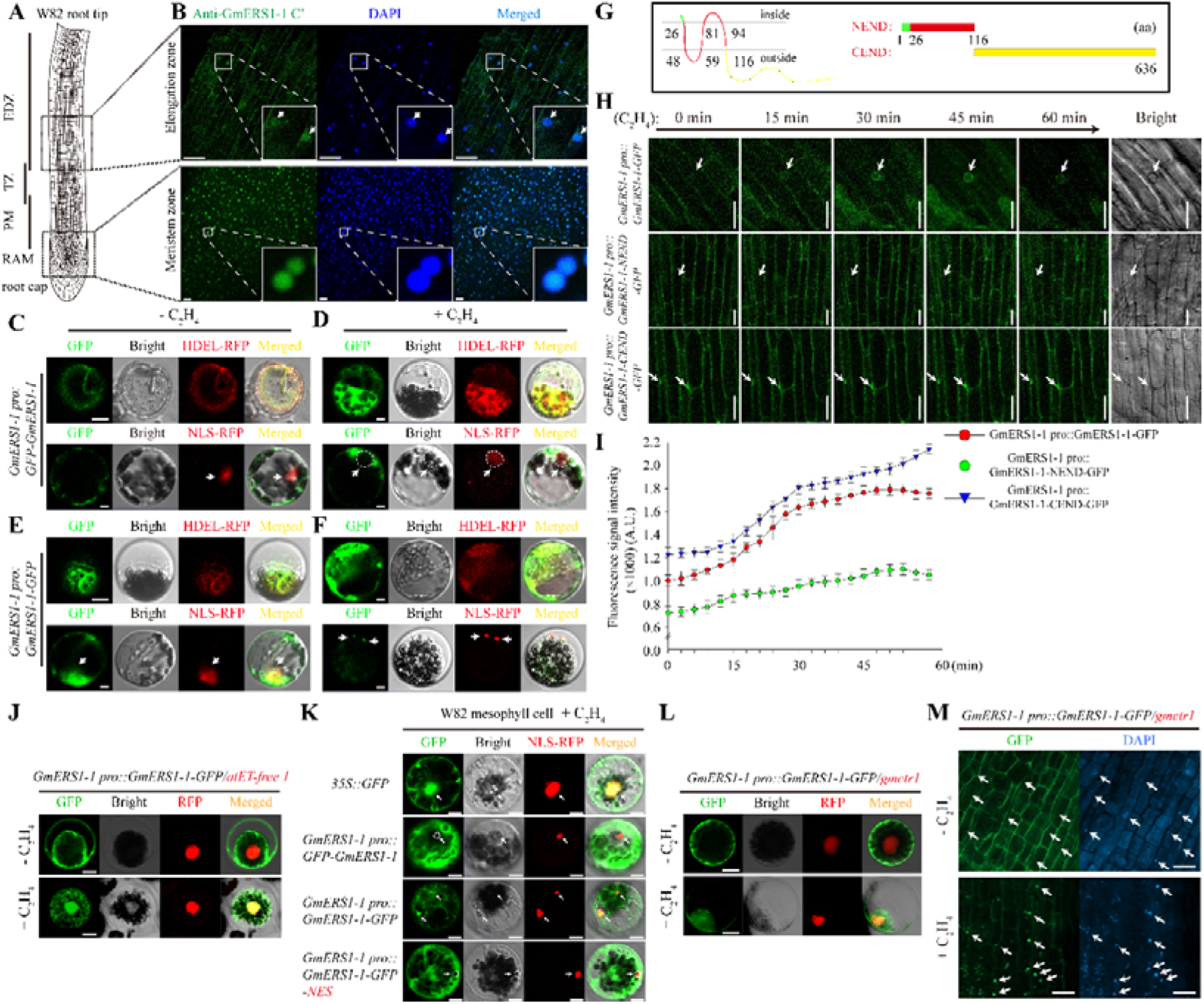
Under the treatment of ethylene, the C’ terminal of GmERS1-1 is transferred to the nucleus. **(A-B)** Localization of GmERS1-1 in soybean root tip cells. **(A)** The left side is a schematic diagram of root sampling. The figure on the left shows a complete root tip. In the figure on the right is an enlarged view of thse two specified positions, meristematic zone and elongation zone. EDZ, elongation and differentiation zone; TZ, transition zone; PM, proximal meristem. RAM, apical meristem tissue. Scale bar, 0.1 mm. **(B)** Immunofluorescence localization of GmERS1-1 protein in two different parts of the root tip. The enlarged area is located in the lower right corner of each field of view. Those arrows pointed to the nucleus. Green represents GmERS1-1 localization (Anti GmERS1-1 C’) and blue represents DAPI. The material used is the root of wild-type Williams 82 grown under normal conditions. Scale bar, 20 μm. **(C-F)** Expression of *GmERS1-1-GFP* in soybean protoplast cells using polyethylene glycol mediated method. The (**C** and **D**) GFP protein is located before the N’-terminus of GmERS1-1. The (**E** and **F**) GFP protein is located behind the C’-terminus of GmERS1-1. After 16 hours of conversion, treat with C_2_H_4_ (right half). After 30 minutes of treatment with C_2_H_4_ (100ppm), the expression of the introduced gene was observed. NLS-DsRed (nuclear localization signal Discosoma sp. red fluorescent protein) co transformed into nuclear markers. Co transform HDEL-DsRed (endoplasmic reticulum localization signal *Discosoma* sp. red fluorescent protein) into an endoplasmic reticulum marker. Using a laser confocal scanning microscope (Zeiss Microsystem LSM 700, http://www.zeiss.com.cn/microscopy). Collect images and encode them in green and red as GFP and DsRed/HDEL, respectively. Merge two confocal cross-sectional images into a two-dimensional image. Arrows mark the nucleus. Scale bar, 10 μm. **(G)** GmERS1-1 segmented schematic diagram. On the left is a transmembrane schematic diagram of the amino acid sequence of GmERS1-1. The corresponding amino acid positions are marked in the diagram. On the right is a schematic diagram of our segmentation of GmERS1-1. Among them, NEND contains the amino acid sequence of positions 1-116, and CEND contains the amino acid sequence of positions 117-636. **(H)** Time delayed confocal images of root cells in 3-day-old soybean hairy roots treated with 100 ppm ethylene gas (GmERS1-1-GFP, GmERS1-1 NEND-GFP, and GmERS1-1 CEND-GFP). From 0 minutes to 60 minutes, we continue to observe the same position of the same material. The expression of these materials is regulated by the self-promoter of *GmERS1-1*. The arrow points to the nucleus, indicating the accumulation of GmERS1-1 and its fragment (green fluorescence) responsive to ethylene. The bright field of view is displayed on the right. Scale bar, 20 μm. **(I)** Trend chart of fluorescence signal intensity changes for three channels in B. The red dot represents GmERS1-1-GFP, the green dot represents GmERS1-1 NEND-GFP, and the blue triangle represents GmERS1-1 CEND-GFP. Use ZEN blue software to analyze the images. Values are means ± SE (n = 3). **(J)** Localization of GmERS1-1 in the *Arabidopsis thaliana* none ethylene producing *ACS* five mutant *atET-free 1*. The upper part is untreated with ethylene, and the lower part is treated with ethylene. **(K)** GmERS1-1 C’ end nuclear pump verification under ethylene treatment. From top to bottom, there are *35S::GFP*, *GmERS1-1 pro::GFP-GmERS1-1* (GFP at N’ end), *GmERS1-1 pro::GmERS1-1-GFP* (GFP at C’ end), and *GmERS1-1 pro::GmERS1-1-GFP-NES* (nuclear pump output signal at C’ end of GFP). NLS-RFP as a nuclear localization marker. The white arrows pointed towards the nucleus. **(L)** Localization of GmERS1-1 in the protoplasts of soybean *CTR1* mutant *gmctr1*. The upper part is untreated with ethylene, and the lower part is treated with ethylene. Scale bar, 5 μm. **(M)** Localization of GmERS1-1 in the root tips of soybean *CTR1* mutant *gmctr1*. The upper part is untreated with ethylene, and the lower part is treated with ethylene. The red arrows pointed towards the nucleus. Scale bar, 10 μm.

To observe the process of GmERS1-1 protein entering the nucleus after ethylene treatment, we completed transformation of three vectors including *GmERS1-1 pro:GmERS1-1-GFP* (GFP fused with full-length GmERS1-1), *GmERS1-1 pro:GmERS1-1-NEND-GFP* (GFP fused with free N-terminus of GmERS1-1, GmERS1-1-NEND, 1-116 aa) and *GmERS1-1 pro:GmERS1-1-CEND-GFP* (GFP fused with free C-terminus of GmERS1-1, GmERS1-1-CEND, 117-636 aa), respectively by the soybean hairy root transformation system **(Fig. 3, G and H)**. Time-lapsed imaging was used to monitor subcellular positions dynamic changes of GmERS1-1 fusion protein upon ethylene treatment. The results showed that at time point 0 min (in the absence of ethylene), no GFP signal was observed in the nucleus of *GmERS1-1 pro::GmERS1-1-GFP* **(Movie. S1)** and *GmERS1-1 pro::GmERS1-1-NEND-GFP* **(Movie. S2)**, but GFP signal appeared in part of the nucleus of *GmERS1-1 pro::GmERS1-1-CEND-GFP* transformations **(Movie. S3)** (**Fig. 3, H and I**). With the following of the treatment time, GFP signals began to accumulate in the nucleus of lines *GmERS1-1 pro:GmERS1-1-GFP* and *GmERS1-1 pro:GmERS1-1-CEND-GFP* at time point 15 min, and the signal intensity reached the strongest at time point 45 min, whereas no GFP signals still appeared in the nucleus of *GmERS1-1 pro:GmERS1-1-NEND-GFP* line (**Fig. 3, H and I**).

In order to demonstrate that nuclear localization of GmERS1-1 specificity is induced by ethylene, we obtained *Arabidopsis* mutant *atET-free-1* of free ethylene production, which reduce the interference of ethylene synthesis by cells itself (*22*) and completed transient protoplasts transformation of *atET-free-1* mutant cells using the vector *GmERS1-1 pro::GmERS1-1-GFP*. The result showed that in mutant *atET-free-1*, there wasn’t signal of GFP in the cell nucleus before ethylene treatment, and only external application of ethylene can lead to GFP signal in the nucleus **(Fig. 3J)**, which indicated that the entry of GmERS1-1 into the nucleus only occurs in the presence of an ethylene signal. To further determine whether the entering of GmERS1-1 into the nucleus were related to the C-terminus of GmERS1-1, we connected the nuclear pumping signal (NES) with the C-terminus of GmERS1-1 to put the C-terminal out of nucleus, and observe changes in the nucleus localization of the C-terminal of GmERS1-1. The results showed that there was no fluorescence signal in the nucleus when the NES was connected with C-terminus of GmERS1-1 **(Fig. 3K)**. These results demonstrated that the C-terminus of GmERS1-1 was crucial for the nuclear localization of GmERS1-1.

As a direct downstream modulator of ethylene receptor, CTR1 (Constitutive Triple Response 1) has been firmly established as a negative regulator in the ethylene signaling pathway (*23*). To verify whether the CTR1 affects the entry of GmERS1-1-CEND into the nucleus, we completed transient transformation with the vector *GmERS1-1 pro:GmERS1-1-GFP* on *gmctr1* soybean mutant background through the transient transformation method, and found that the nuclear localization of GmERS1-1 was not affected in *gmctr1* soybean mutant **(Fig. 3, L and M)**. The results showed that in the absence of CTR1, the ethylene still can induce nuclear localization of GmERS1-1 in both soybean protoplasts **(Fig. 3L)** and root tip cells **(Fig. 3M)**.

To verify the conservation of ethylene receptors entering the nucleus, we analyzed the subcellular localization of all the members in soybean and *Arabidopsis* **(fig. S7)**, and the results showed that two individual members of subfamily Ⅰ had no nuclear localization **(fig. S7A)**, whereas all the members of the subfamily Ⅱ had nuclear distribution **(fig. S7B)**, which suggesting that members of subfamily Ⅱ have conserved characteristics to enter the nucleus in *Arabidopsis* and soybean. We calculated the proportion of cells with GmERS1-1 located in the nucleus, and found that the cells number with the nucleus localization transformed using *GmERS1-1 pro: GmERS1-1-GFP* vector were higher after ethylene treatment than that before treatment, and the GFP signal in the nucleus were also stronger after ethylene treatment than that before treatment **(fig. S8, A and B)**. In addition, we observed nuclear localization of overexpressing *GmERS1-1* (*GmERS1-1-OE*, GFP was fused in C-terminus of GmERS1-1) in transgenic *Arabidopsis* (**fig. S9, A to D**), which indicated that the nuclear localization of ethylene receptor GmERS1-1 is conserved in soybean and *Arabidopsis*.

### Free GmERS1-1-CEND can transmit ethylene signals by entering the nucleus in soybean and *Arabidopsis*

We have demonstrated that overexpression of full-length *GmERS-1-1* can enhance ethylene response **(Fig. 2, A to C)**. Next, free *GmERS1-1-CEND* was expressed under background of *Arabidopsis aters1* mutants and wild type soybean W82, respectively to identify the functions of the GmERS1-1-CEND in ethylene signaling pathway by the triple reactions assay. The results showed that overexpression of soybean *GmERS1-1-CEND* could complement the ethylene sensitivity of the *Arabidopsis aters1* mutant, which were similarly with functions of full-length GmERS1-1, whereas free *GmERS1-1-NEND* can’t complement the ethylene sensitivity of *Arabidopsis aters1* mutant **(Fig. 4, A to C)**. Especially, overexpression of *GmERS1-1-CEND* could further enhance ethylene sensitivity under wild type *Arabidopsis*, specifically manifested as the shortening of hypocotyl and root length, which was similarly with plant overexpressing full-length *GmERS1-1* and was different with plants overexpressing *GmERS1-1-NEND* **(Fig. 4, A to C)**. Considering the protein structural similarity between GmERS1-1 and AtERS1 **(fig. S5A)**, we speculated that they have similar functions for transmitting ethylene signals. Finally, we found that the free C-terminus of ethylene receptor *AtERS1* (*AtERS1-CEND*) could also complement the *aters1* mutant phenotype, but *AtERS1-NEND* could not **(fig. S10, A to C)**, which indicated that the functions of free AtERS1-CEND and GmERS1-1-CEND are conservative and can transmit ethylene signals in plants. At the same time, we found that there are full-length GmERS1-1 and GmERS1-1-CEND in total protein isolated from plants, whereas only the GmERS1-1-CEND was detected in proteins isolated from the nucleus **(Fig. 4, D and E)**. The GmERS1-1-CEND were present in both wild-type *Arabidopsis* and *Arabidopsis* overexpressing full-length *GmERS1-1* or *GmERS1-1-CEND*, but not in the *aters1* mutant and *GmERS1-1-NEND/ates1* **(Fig. 4, D and E)**. The abundance of GmERS1-1-CEND in *Arabidopsis* overexpressing full-length *GmERS1-1* or *GmERS1-1-CEND* were greater than that in wild type *Arabidopsis* (**Fig. 4, D and E**). Meanwhile, we overexpressed both *GmERS1-1-CEND* and *AtERS1-CEND* in soybean using the soybean root transient expression system, and found that overexpressed both of free *GmERS1-1-CEND* and *AtERS1-CEND* could significantly inhibit the elongation of soybean roots (**fig. S11, A and B**), and we can detect nuclear distribution of free GmERS1-1-CEND and free AtERS1-CEND in soybean root tip cells **(fig. S11C)**. Soybean root tip cells overexpressing *GmERS1-1-NEND* and *AtERS1-NEND* had no nuclear distribution of GmERS1-1-CEND and AtERS1-CEND, and *GmERS1-1-NEND* and *AtERS1-NEND* could not inhibit roots growth **(fig. S11, A to C)**. These results demonstrated that GmERS1-1-CEND and AtERS1-CEND have similar functions of entering the nucleus to induce ethylene signals in soybean and *Arabidopsis*, which resulted in inhibition of root elongation and *Arabidopsis* hypocotyl elongation.

**Figure 4.**
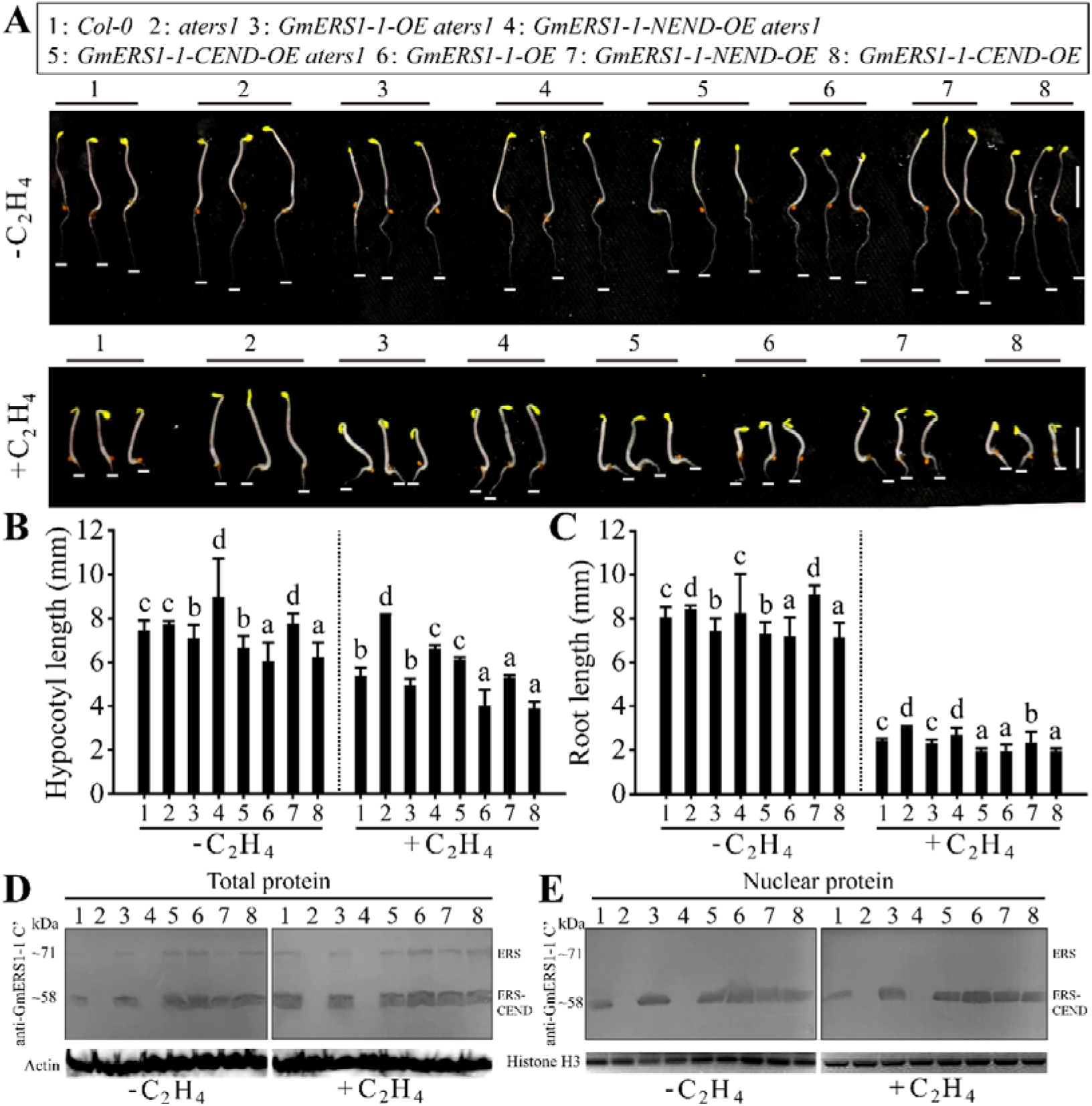
Overexpression of the C’-terminal of *GmERS1-1* can enhance the triple reaction in *Arabidopsis*. After obtaining the mutant *aters1* of Arabidopsis *AtERS1*, the following materials were constructed: **(A)** *GmERS1-1* full length complementary *aters1*, *GmERS1-1 NEND* complementary *aters1*, *GmERS1-1 CEND* complementary *aters1*, overexpressing *GmERS1-1* full length, overexpressing *GmERS1-1* NEND, and overexpressing *GmERS1-1* CEND. The triple response phenotype validation was conducted, and the results showed that the soybean *GmERS1-1* had an enhanced effect on the triple response, manifested as a shortened hypocotyl **(B)** and a shortened root length **(C)**. Data statistics conducted ANOVA-DUNCAN’s multiple range tests at P<0.05, error bars report SE of the mean, n=3. Among them, overexpression of *GmERS1-1 CEND* alone was the most significant, while overexpression of *GmERS1-1 NEND* alone had the least significant effect. This indicates that *GmERS1-1 CEND* plays an important role in the triple reaction process. Changes in protein expression levels before and after ethylene treatment **(D-E)**. (D) Extract the total protein of (A) and perform Western blot (anti GmERS1-1 CEND). The results show that GmERS1-1 CEND does indeed exhibit truncation, and after ethylene treatment, this truncation has a strengthening effect. (E) Extract the nucleus of (A) and perform Western blot (anti GmERS1-1 CEND). The results showed that the GmERS1-1 CEND content in the soybean nucleus increased after ethylene treatment.

### GmERS1-1 regulates the cell cycle by interacting with GmACR1

In order to determine mechanisms of GmERS1-1 after entering the nucleus, we first utilized the String online tool to conduct predictive analysis on the protein-protein interaction (PPI) relationships of GmERS1-1 **(Fig. 5, A to B)**. In addition to the existing members of the ethylene signaling pathway, the results also indicate that some of the interacting proteins of GmERS1-1 are related to stem cell division **(Fig. 5A)**. This piqued our interest, as we have always been eager to unravel the intrinsic mechanism of ethylene-induced growth inhibition. Afterwards, we used GmERS1-1 as bait to screen for interacting proteins through yeast two hybrid system (Y2H). We found that GmERS1-1 can interact with an arsenate reductase GmACR1(Glyma.19G185200) directly by screening of soybean yeast library induced by ethylene **(Fig. 5B)**. Based on this, we analyzed the protein domains and three-dimensional configurations of GmACR1, human (*Humo sapiens*) CDC25B and AtACR1 (AT5G03455), and the results showed that GmACR1 does indeed have a RHOD domain similar to human CDC25B and AtACR1 **(Fig. 5C)**. Amino acid sequence alignment revealed similarities among the three proteins, with a similarity of 16% **(fig. S12)**. GmACR1 and AtACR1 are like truncated human CDC25B. These phenomena suggest that in addition to enzyme activity, GmACR1 may indeed have functions similar to human CDC25B in regulating the cell cycle. In addition, we compared the genomes of common model plants and leguminous plants and found that only one *ACR* gene exists in soybean, which further illustrates the importance of GmACR1 **(Fig. 5D)**.

**Figure 5.**
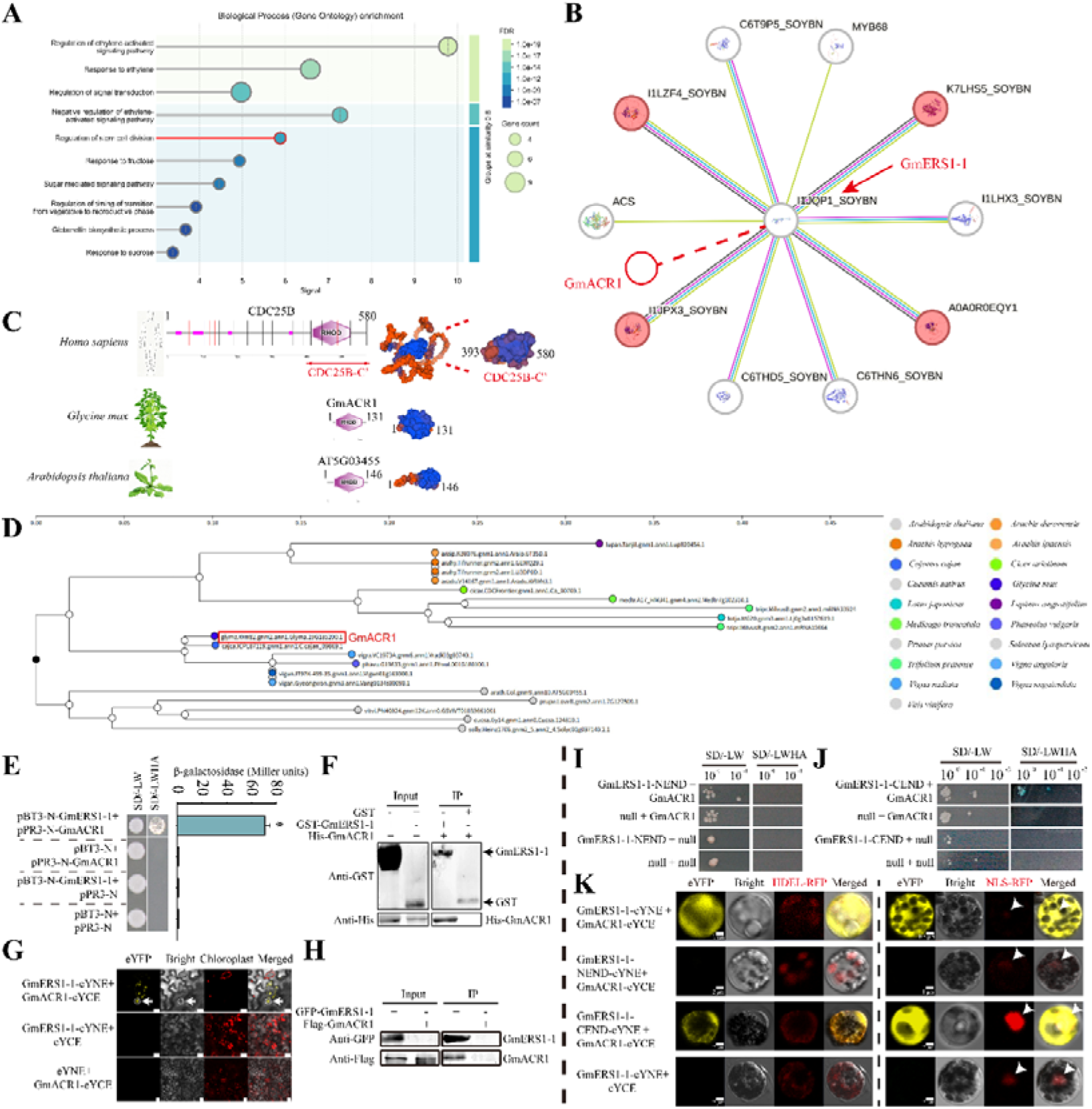
GmERS1-1 can interact with GmACR1 in nucleus. **(A)** Bubble plot for Gene Ontology Biological Process enrichment of GmERS1-1 related genes. The term “Regulation of stem cell division” is highlighted red. Bubble size represents the number of enriched genes, color gradient corresponds to FDR significance. **(B)** Protein-protein interaction network of GmERS1-1. Four proteins annotated to stem cell division regulation are marked red; candidate interactor GmACR1 is connected to GmERS1-1 via a red dashed line. **(C)** Conserved domain prediction (SMART) and SWISS-MODEL homology structures for GmACR1, human CDC25B and AtACR1. Purple domains denote RHOD domains; separate model was built for human CDC25B C-terminal fragment (393–580 aa). **(D)** Gene family classification of GmACR1 from Legume Information System database (SoyBase). GmACR1 is boxed red; colored dots in legend represent different plant species. **(E)** Yeast two-hybrid assay detecting GmERS1-1-GmACR1 interaction. β-galactosidase activity quantifies binding strength. Empty pBT3-N/pPR3-N vectors act as negative controls for the experimental co-expression group. **(F)** In vitro GST pull-down assay validating direct GmERS1-1-GmACR1 binding. Input lanes confirm equal protein loading; anti-GST and anti-His antibodies immunoblot captured protein complexes. **(G)** Tobacco epidermal cell BiFC assay. GmERS1-1 and GmACR1 were fused to eYFP-N and eYFP-C halves, respectively. eYFP, chloroplast and merged fluorescence channels are shown; empty split-YFP vectors serve as negative controls. **(H)** In vivo co-IP assay verifying intracellular GmERS1-1-GmACR1 complex. GFP-GmERS1-1 and Flag-GmACR1 were co-expressed. Anti-GFP antibody pulled down complexes, which were immunoblotted with anti-Flag antibody; input lanes confirm total protein abundance. **(I)** Truncation BiFC mapping of the GmERS1-1 interaction domain with GmACR1 in soybean protoplasts. Serial N/C-terminal GmERS1-1 deletions were tested. HDEL-RFP marks ER, NLS-RFP labels nucleus; white arrowheads point to nuclear eYFP signals. Scale bar = 2 μm.

In this research, we found that the overexpression of *GmERS1-1* can enhance the ethylene sensitivity of plants, which caused inhibition (**Fig. 2 and Fig. 4**), suggesting that GmERS1-1 could regulate plants growth through the interaction with GmACR1. To investigate the function of GmACR1, we first validated the protein interaction between GmERS1-1 and GmACR1. We verified the interaction between GmERS1-1 and GmACR1 through several methods, including the Y2H, the Pull-down assay, the bimolecular fluorescence complementation (BiFC) and the immunoprecipitation (Co-IP) **(Fig. 5, E to K)**. The results of the BiFC showed that GmERS1-1 and GmACR1 can interact in the nucleus and the endoplasmic reticulum (ER) **(Fig. 5G)**. In addition, we also verified the interaction between free GmERS1-1-NEND and GmERS1-1-CEND, and GmACR1, respectively, and the results showed that free GmERS1-1-CEND can interact with GmACR1 in the ER and nucleus, which was similarly with the subcellular localization of interacting protein pairs full-length GmERS1-1 and GmACR1, whereas free GmERS1-1-NEND cannot interact with GmACR1 **(Fig. 5K)**.

Firstly, we verified whether the expression of *GmACR1* was induced by ethylene. Promoter activity analysis showed that the expression of *GmACR1* was indeed induced by ethylene **(Fig. 6, A and B and fig. S13)**. We also tested the triple responses of the *GmACR1* nonsynonymous mutant *gmacr1* **(Fig. 6C)**. The results showed that *gmacr1* had longer hypocotyls and roots, whereas wild type W82 had shorter hypocotyls and roots after ethylene treatment **(Fig. 6, D to F)**, which indicated that mutation of *GmACR1* inhibited ethylene sensitivity in soybean. Then, using hairy root transformation technique, we confirmed that overexpression of *GmACR1* also inhibited the growth of soybean under normal conditions in pot culture and hydroponic culture (**Fig. 6, G to L**). The main manifestation is a reduction in plant height (**Fig. 6, G and H**), root fresh weight (**Fig. 6, I and J**), and dry weight (**Fig. 6K**) of *GmACR1* overexpression lines (**Fig. 6L**). Consistent with this observation, 4′,6-diamidino-2-phenylindole staining and flow cytometry analyses showed that some cells in the overexpressed *GmACR1* lines roots had 8C DNA content compared with WT (4C), suggesting a delay in mitotic exit in some overexpressed *GmACR1* lines roots cells **(Fig. 6, M and N)**. In order to further verify the distribution of GmACR1 protein in soybean cells, we confirmed that GmACR1 is distributed in the cytoplasm and nucleus through immunofluorescence techniques **(Fig. 6O)** and subcellular localization of soybean protoplasts **(Fig. 6P)**. In addition, ethylene treatment would promote GmACR1 to further accumulate into the nucleus (**Fig. 6P**). In order to verify this hypothesis, we separately verified the effect of *GmERS1-1* and *GmACR1* on cell cycle. Overexpression of *GmERS1-1* and *GmACR1* in *Schizosaccharomyces pombe* inhibited cell division and caused cell enlargement **(Fig. 6, Q and R)**. The number of cells in the visual field showed that 41.5% and 35.7% of the cells were in the G_2_/M phase with overexpressed *GmERS1-1* and *GmACR1*, respectively **(Fig. 6S)**. In order to more clearly reflect the functions of *GmERS1-1* and *GmACR1* in influencing soybean growth, we overexpressed *GmERS1-1* and *GmACR1* in soybean hairy roots **(Fig. 6T)**. The results also showed that overexpression of *GmERS1-1* and *GmACR1* inhibited the growth of soybean roots, with *GmACR1* having a stronger effect **(Fig. 6U)**.

**Figure 6.**
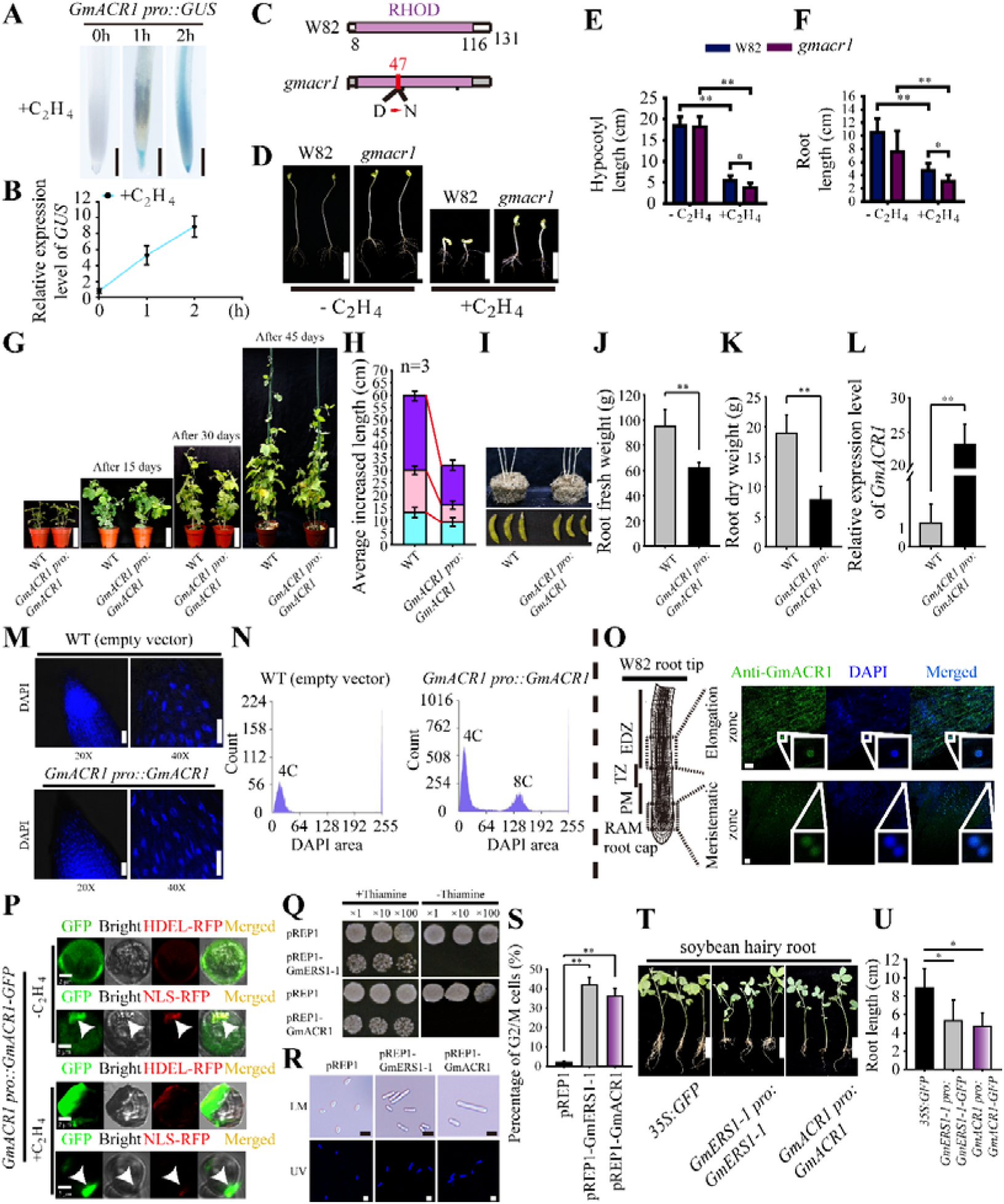
Validation of the function of GmACR1 on plant growth. **(A)** Continuous observation of root tips of *GmACR1 pro::GUS* line after ethylene treatment. Scale bar=1mm. **(B)** Changes in *GUS* gene expression during ethylene treatment. The error line represents SE of the mean. Statistical analysis was conducted using Student’s *t*-test. n=3. **(C-F)** The phenotype of “triple reaction” of wild-type Williams 82 (W82) and the mutant line *gmacr1* of *GmACR1*. **(C)** Display of mutation sites in the mutant line *gmacr1*. The number represents the number of amino acids. The purple domain (8–116) indicates Rhodanese Homology Domain (RHOD). 47/131: D->N (Non-synonymous). **(D)** The group without ethylene treatment is on the left, and the group with ethylene treatment is on the right. Each group contains two biological replicates. Bar=5cm. **(E-F)** Statistical analysis of hypocotyl length (**E**) and root length (**F**) in **D**. The error line represents SE of the mean. Statistical analysis was conducted using Student’s *t*-test. n=6. ** indicates that *P* is less than 0.01. * indicates that *P* is less than 0.05. **(G)** Using *Agrobacterium tumefaciens* to transfer recombinant *35S::GFP* and *GmACR1 pro::GmACR1-GFP* into soybean hairy roots. The figure shows the continuous observation results of soybean growth after transformation in soil. The wild type is on the left, and the overexpressed lines of *GmACR1* is on the right. Scale bar=12cm. **(H)** The vertical axis represents the plant height of soybean growth during each time period. n=3. These results showed that overexpression of *GmACR1* significantly inhibited the growth of soybeans. **(I)** This demonstrates the phenotype of roots washing away soil and pods. **(J-K)** Statistical analysis of fresh root weight **(J)** and dry root weight **(K)**. **(L)** The expression levels of *GmACR1* gene in each line were analyzed. The error line represents SE. Statistical analysis was conducted using Student’s *t*-test. Asterisks indicate comparison with the control group (**, p<0.01). **(M)** DAPI staining of the root tip meristem cell nucleus of soybean (W82 and GmACR1 overexpressing lines). **(N)** Flow cytometry sorting of the nuclei in the meristematic zone of the root tip. **(O)** Immunofluorescence localization of GmACR1 in the root tip of W82 (Anti-Acr1), green GFP signal, blue DAPI signal. The left side is a sampling diagram. The upper right part is the elongation zone, and the lower part is the meristematic zone. Scale bar=10 μm. **(P)** Subcellular localization of GmACR1 in soybean protoplasts (divided into ethylene untreated and ethylene treated), with endoplasmic reticulum markers above and nuclear markers below each treatment group. **(Q)** Growth experiment of *Schizosaccharomyces pombe*. The control group was empty vector pREP1, while the experimental group was pREP1 GmERS1-1 and pREP1 GmACR1. Divided into two treatments, one group added thiamine (gene inhibition), and the other group did not add thiamine (gene normal expression). **(R)** Microscopic observation of the morphology of *Schizosaccharomyces pombe* cells, LM: white light, UV: DAPI staining. **(S)** Count different forms of cells in *S. pombe*, observe at least 50 fields of view per group, and the error line represents SE. Statistical analysis was conducted using Student’s *t*-test. Asterisks indicate comparison with the control group (**, p<0.01). **(T)** Using *Agrobacterium tumefaciens* to transfer recombinant *35S::GFP*, *GmERS1-1 pro::GmERS1-1-GFP*, *GmACR1 pro::GmACR1-GFP* into soybean hairy roots. The picture shows the situation under hydroponic conditions. Scale bar=5 cm. **(U)** Root length statistics. The error line represents SE. Statistical analysis was conducted using Student’s *t*-test. \**P*<0.05.

### Module GmERS1-1-GmACR1 may not require CTR1 to function effectively

Based on subcellular localization of GmERS1-1 in leaf protoplasts and root tip cells of soybean *ctr1* mutant **(Fig. 3, L and M)**, we have tentatively inferred that the entry of GmERS1-1-CEND into the nucleus does not require the involvement of CTR1. To verify whether the function of module GmERS1-1-GmACR1 is affected by the classical ethylene signaling pathway, we analyzed downstream genes (1.5>logFold Change>1.5; FDR < 0.01) using RNA sequencing **(Fig. 7, A to C)**. Among those downstream upregulated genes, we found that the expression of some cell cycle related genes was not affected by *ctr1* mutation **(Fig. 7, A to C)**, such as *GmACR1*, *GmCDK7* (Glyma.08G050900), *GmCyclin-A3-1* (Glyma.14G087100), and so on.

**Figure 7.**
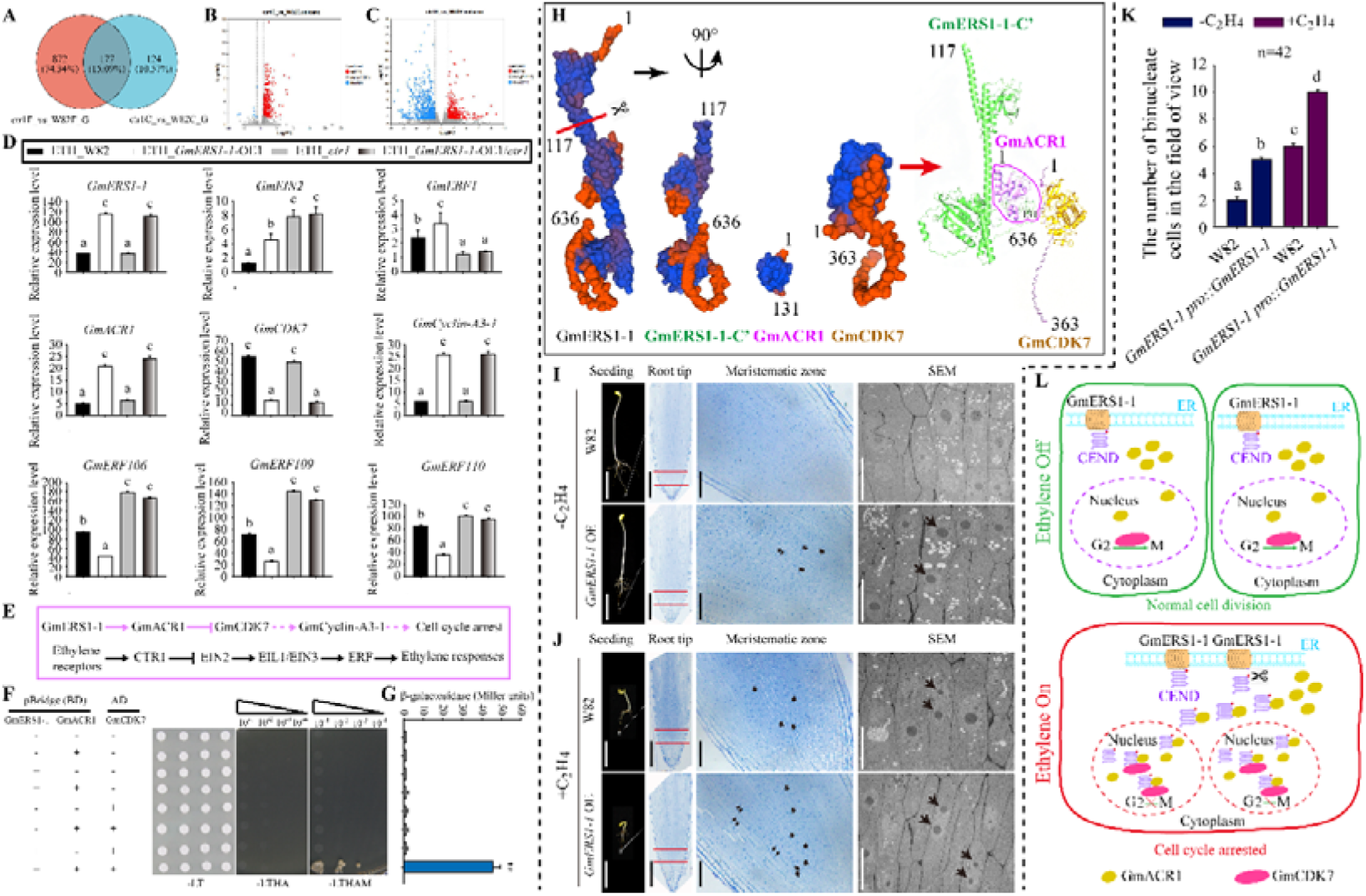
The GmERS1-1-GmACR1-GmCDK7 module is not affected by CTR1. **(A-C)** Transcriptome sequencing of soybean wild-type W82 and *ctr1* mutant after ethylene treatment. **(A)** Venn diagram to show the overlap of compromised ethylene up-regulated genes in W82 and in the *ctr1* mutants. **(B-C)** Volcano plots show the DEGs of different comparative groups under normal **(B)** and ethylene **(C)** conditions. **(D)** Verification of differential gene expression in W82, *GmERS1-1*-OE1, *ctr1* and *GmERS1-1*-OE1/*ctr1* lines under ethylene treatment by using soybean hairy root transient transfection assay. *ACTIN* was used as endogenous control for normalization. Data statistics conducted ANOVA-DUNCAN’s multiple range tests at P<0.05, error bars report SE of the mean, n=3. **(E)** Based on the above results, we propose a hypothetical signaling pathway diagram. GmERS1-1, GmACR1, and GmCDK7 work together to regulate the cell cycle. This pathway is independent of the classical ethylene signaling pathway and does not rely on CTR1. **(F)** Verify the interaction relationship between GmERS1-1, GmACR1, and GmCDK7 using Y3H assay. The colony pictures were taken after 3 days at 30°C in –Leu–Trp (-LT) plates and after 9 days at 30°C in –Leu–Trp–His-Ade (-LTHA) and –Leu–Trp–His-Ade–Met (-LTHAM) plates. **(G)** A quantitative assay for β-galactosidase activity. The presented values are the averages of four independent assays with SE. Statistical analysis was conducted using Student’s t-test. Asterisks indicate comparison with the control group (**, p<0.01). **(H)** On the left is the 3D structure prediction of four proteins, GmERS1-1, GmERS1-1-C’, GmACR1, and GmCDK7, using SWISS-Model. On the right is the protein docking performed on GmERS1-1-C’, GmACR1, and GmCDK7. The protein’s 3D structure was constructed using AlphaFold3. The Hdock docking process involves docking protein GmERS1-1-C’ with protein GmACR1 first, and then docking the protein GmERS1-1-C’-GmACR1 complex with protein GmCDK7. Based on the 3D structure predicted by AlphaFold 3, pymol was used for drawing. The number indicates the number of amino acids. **(I-J)** The plant material in Figure 2A was observed. Longitudinal section and electron microscopy observation of root tip meristematic zone. There is no ethylene treatment group on the top half (H), while the ethylene treatment group is on the bottom half (I). **(K)** Statistics of binuclear cells in the A-B field of view (n=42), error line represents SE. Data statistics conducted ANOVA-DUNCAN’s multiple range tests at P<0.05. **(L)** Working model of ethylene-triggered cell cycle arrest mediated by soybean GmERS1-1. In the absence of ethylene (green box), intact ER-localized GmERS1-1 retains its CEND domain. Limited nuclear GmACR1 allows functional GmCDK7 to drive G2/M transition and sustain normal cell division. When ethylene accumulates (red box), GmERS1-1 undergoes proteolytic cleavage to release soluble CEND fragments. These CEND peptides recruit bulk cytoplasmic GmACR1 into the nucleus, where excess GmACR1 inhibits GmCDK7 activity, blocks G2-to-M progression, and causes cell cycle arrest. Yellow dots, GmACR1; magenta ovals, GmCDK7; scissors, proteolytic cleavage site; ER, endoplasmic reticulum.

To verify the reliability of the results, we used qRT-PCR to detect the expression of these genes in W82, *GmERS1-1*-OE1, *ctr1* and *GmERS1-1*-OE1/*ctr1* soybean hairy root lines under ethylene treatment. The results showed that the expression of *GmERS1-1* was not affected by the mutation of *CTR1* **(Fig. 7D)** and those components of the classical ethylene signaling pathway were affected by the mutation of *CTR1*, such as *EIN2* (Glyma.10G058300), *EBF1* (Glyma.14G116800) **(Fig. 7D)**. While those components related to the cell cycle were not affected, such as *GmACR1*, *GmCDK7*, *GmCyclin-A3-1* **(Fig. 7D)**. In addition, the expression of some ethylene response factors downstream of the ethylene signal is also affected by the mutation of *CTR1*, such as *Gm ERF106* (Glyma.20G070100), *GmERF109* (Glyma.07G212400), *GmERF110* (Glyma.16G164800) **(Fig. 7D)**.

### GmERS1-1, GmACR1 and GmCDK7 can form a trimer to regulate the cell cycle

Our research has demonstrated the relationship between GmERS1-1 and GmACR1 **(Fig. 5 and Fig. 6)**. Studies suggest that CDK activity affected by CDC25 phosphatase at G₂/M (*24*, *25*). Gene expression data shows that *GmERS1-1* and *GmACR1* have different expression pattern changes with *GmCDK7* **(Fig. 7D)**. Based on the spatiotemporal specificity distribution and gene expression of module GmERS1-1-GmACR1-GmCDK7, we speculate that module GmERS1-1-GmACR1-GmCDK7 may be independent of the classical ethylene signaling pathway **(Fig. 7E)**. Based on these results, we speculate that there may be some association between GmERS1-1, GmACR1, and GmCDK7. Yeast three-hybrid (Y3H) analysis verified that GmERS1-1 can form trimer with GmACR1 and GmCDK7 **(Fig. 7, F and G)**. In order to analyze the relationship between GmERS1-1-GmACR1-GmCDK7 ternary complexes, we further predicted those three-dimensional structures of GmERS1-1, GmERS1-1 C’, GmACR1 and GmCDK7 by using the SMART online tool. The results show that there is indeed an interaction space between GmERS1-1 C’, GmACR1 and GmCDK7 in terms of structure **(Fig. 7H)**. To further verify whether GmERS1-1 C’, GmACR1 and GmCDK7 can truly form complexes, we used AlphaFold3 for AF3 modeling and analysis of ternary complexes, with Hdock docking. The results further confirm that the GmERS1-1 C’-GmACR1-GmCDK7 ternary complex may indeed exist **(Fig. 7H)**.

The above experimental results demonstrate that ethylene inhibits growth by inhibiting cell division, which is the most fundamental pathway. In order to visually display the binuclear cells, we conducted a more direct electron microscopy observation. In the triple response experiment, we found the presence of binuclear cells in soybean root tip cells of *GmERS1-1* over-expressed lines, and ethylene treatment will increase its proportion **(Fig. 7, I to K)**. Although we demonstrated that GmERS1-1-GmACR1-GmCDK7 complex controls cell cycle in soybean, we wanted to know whether this represents a general mechanism underlying post-harvest biology across plant species. The ripening stage of apples is a very important stage in the growth and development of fruit trees, which has a significant impact on yield and fruit quality. The two important stages of apple ripening are the swelling stage and the coloring stage. We chose apples that are in the swelling stage as the processing object, as these apples are in a rapid growth phase. We treated the harvested apples (*Malus pumila* ‘Red Fuji’) with ethylene in the swelling stage and analyzed their nuclear ploidy **(fig. S14, A to F)**. The results showed that ethylene does indeed block the cell cycle during the ripening process of apples.

Based on these above results, we can draw a conclusion: GmERS1-1-CEND enters into the nucleus and interacts with GmACR1 under the action of ethylene signal. By affecting the GmERS1-1-GmACR1-GmCDK7 complex, the cell cycle stops at the G_2_/M stage, thereby inhibiting plant growth (**Fig. 7L**).

## DISCUSSION

### Nuclear translocation of ethylene signaling components: comparative analysis and future directions

The nuclear translocation of C-terminal domains in ethylene signaling components represents a conserved yet functionally diversified mechanism across plant species. For instance, in *Arabidopsis*, the C-terminal end of EIN2 (EIN2-CEND) is cleaved and translocated to the nucleus under ethylene signaling, where it stabilizes the transcription factor EIN3/EIL1 to activate downstream genes (*8*). In addition, Park et al. reveals a novel regulatory mechanism in ethylene signaling (*23*). Contrary to previous assumptions, the nuclear translocation of CTR1—a negative regulator of ethylene signaling—does not require the canonical components EIN2 or EIN3, as demonstrated by genetic analyses. Instead, the N-terminal domain of CTR1 is essential for its nuclear localization, as truncation of this region blocks ethylene-induced nuclear movement. This nuclear retention of CTR1 delays hypocotyl elongation recovery after ethylene removal, suggesting that its subcellular dynamics fine-tune the balance between ethylene sensitivity and growth adaptation. These findings redefine CTR1 as a dual-function regulator, transitioning from a cytoplasmic repressor to a nuclear co-activator, and highlight the importance of spatial control in hormone signaling. Similarly, our data showed that the soybean ethylene receptor GmERS1-1 requires its C-terminal domain (GmERS1-1-CEND) for nuclear entry, where it interacts with GmACR1 and GmCDK7 to regulate the cell cycle.

All these cases highlight the critical role of C-terminal domains in nuclear signaling, often involving proteolytic cleavage for functional activation. However, divergence exists in downstream pathways: while EIN2-CEND operates through the canonical EIN3/EIL1 transcriptional cascade; Under normal conditions, CTR1 is recruited to the endoplasmic reticulum (ER) via interaction with ethylene receptors such as ETR1, forming a complex that suppresses ethylene responses. However, when CTR1 dissociates from ETR1, it accumulates in the nucleus, where it stabilizes the transcription factor EIN3 by directly interacting with EBF proteins (EIN3-BINDING F-BOX), thereby enhancing ethylene signaling independently of its kinase activity; GmERS1-1-CEND directly modulates cell cycle regulators, suggesting a non-canonical ethylene pathway. Notably, rice OsEIN2 also exhibits nuclear translocation but integrates ethylene signals with ABA biosynthesis, underscoring species-specific adaptations (*26*). Key differences include structural domain specialization. Unlike EIN2, which relies on N-terminal interactions (e.g., with MHZ3 in rice) for stability (*27*), GmERS1-1’s N-terminal domain primarily mediates ethylene perception, while its C-terminal domain independently governs nuclear functions.

Future research should prioritize identifying nuclear localization signals (NLS) in GmERS1-1-CEND and comparing them with EIN2-CEND’s conserved motifs. Additionally, elucidating the proteases responsible for GmERS1-1 cleavage and validating cross-species conservation (e.g., in monocots like maize) are essential. Potential agricultural applications, such as editing GmERS1-1-CEND to optimize soybean architecture, must balance yield gains with unintended impacts on stress responses. Together, the nuclear role of GmERS1-1-CEND unveils a different mechanism by which ethylene receptors regulate cell cycle progression through domain-specific partitioning, offering dual targets for enhancing crop productivity and resilience.

### Comparative analysis of ethylene receptor roles in plant growth and development

Ethylene receptors play a conserved role in hormone signaling across plant species. In soybeans, GmERS1-1 regulates growth and cell cycle progression, akin to ETR1 in *Arabidopsis*, which mediates ethylene responses in organ senescence and stress adaptation (*28*). Similarly, rice ethylene receptors integrate ethylene signals into developmental programs, albeit with distinct regulatory networks (*26*). Our data showed that ethylene’s ability to induce G_2_/M phase arrest, observed in both soybean roots and apple fruits, aligns with conserved cell cycle checkpoints governed by CDK-cyclin complexes. These findings underscore ethylene’s universal role in linking hormone signaling to cell cycle dynamics.

Divergent mechanisms emerge in receptor functionality and pathway independence. Unlike *Arabidopsis* ETR1, which relies on its N-terminal domain for signal transduction, GmERS1-1 requires its C-terminal domain (GmERS1-1-CEND) for nuclear localization and cell cycle regulation, highlighting structural diversification. Furthermore, the GmERS1-1/GmACR1/GmCDK7 trimer operates independently of the canonical ethylene pathway (e.g., EIN3/EIL1), contrasting with *Arabidopsis* where ethylene-triggered apical hook development depends on EIN3-EIL1/PIF4-mediated transcription (Zhao et al., 2020). This suggests species-specific adaptations in ethylene signaling cascades.

While ethylene receptors retain core signaling functions, their molecular mechanisms exhibit species-specific innovations. Future studies should resolve structural determinants (e.g., GmERS1-1-CEND’s nuclear role) and validate cross-species conservation of ethylene-induced cell cycle arrest. Targeting ethylene-responsive modules (e.g., GmERS1-1-CEND) via gene editing could optimize crop architecture, offering novel strategies for agricultural improvement.

### Limitations of the study

In the current study, the specific molecular mechanisms underlying GmERS1-1-CEND’s nuclear functions—including phosphorylation modifications and protein interaction sites—remain incompletely characterized, and its independence from the classical ethylene signaling pathway requires further validation. Functional analyses have primarily relied on phenotypic observations from gene mutation and overexpression experiments, leaving gene redundancy/compensatory effects unexplored and the conservation of this mechanism in monocot plants unverified at the molecular level. Future research should focus on identifying GmERS1-1-CEND’s direct target genes using ChIP-seq or CUT&Tag technologies to clarify its regulatory role on G_2_/M-phase-related genes, while concurrently investigating the phosphatase activity of GmACR1 and its interaction with GmCDK7. Additionally, the relationship between this module and classical ethylene signaling pathways should be rigorously validated, alongside exploring potential cross-talk with other hormonal pathways in cell cycle regulation. Conservation studies should extend to functional validation of *GmERS1-1* homologs in rice or wheat, supplemented by field trials and multi-omics analyses to evaluate environmental adaptability. Translational applications leveraging gene-editing tools and small-molecule modulators could advance agricultural innovation, while constructing dynamic network models integrating ethylene-regulated cell cycle mechanisms with multi-omics datasets would provide a holistic understanding of their roles in organ development and photosynthetic efficiency.

## ACKNOWLEDGMENTS

We thank Daoping Wang for proteome analysis and the help with HPLC-MS analyses, Qingxin Song for providing *gmers1-1* and *gmctr1* mutant lines, Lijia Qu for providing *atET-free-1* mutant lines, Lijuan Qiu for providing assistance in soybean genetic transformation, Haiyang Wang for providing guidance on the conceptual layout of the article and LetPub (www.letpub.com) for editing assistance on the manuscript. **Funding:** This research was supported by the Major Science and Technology Projects in Biological Breeding (NK2023080205), Innovation Program of Chinese Academy of Agricultural Sciences (CAAS-CSCB-202401 and CAAS-ZDRW202403), and the National Natural Science Foundation of China (32272099). **Author contributions:** Y.M., M.C. directed the research. K.C. performed most of the experiments. Y.Z. and W.T. contributed to the transformation construction and phenotype analysis of genetic materials. D.W., Y.B. and C.W. assisted in doing the protein mass spectrometry analysis. W.L., X.L., D.G., M.Z., J.Y., J.G., Y.Z. and B.S. provided assistance in field experiments. J.C., Z.X., Y.H., J.M., Q.S., H.L. and L.Q. provided assistance in experimental instruments. Y.M. M.C. oversaw the entire study. K.C. wrote the manuscript. All authors discussed the results and commented on the manuscript. **Data, code, and materials availability:** All data are available in the manuscript or the supplementary materials. The raw RNA-seq data have been deposited in the NCBI BioProject under accession number PRJNA1503193. Materials are available upon request from the corresponding author.

## REFERENCES AND NOTES

1. J. Bramsiepe, et al., PLoS Genet. 6, e1000996 (2010).

2. R. Pierik, D. Tholen, H. Poorter, E. J. Visser, L. A. Voesenek, Trends Plant Sci. 11, 176–183 (2006).

3. T. Xin et al., Plant Cell 31, 1063–1076 (2019).

4. E. Stoynova-Bakalova, D. V. Bakalov, T. I. Baskin, PeerJ 10, e14315 (2022).

5. A. Swiatek, M. Lenjou, D. Van Bockstaele, D. Inzé, H. Van Onckelen, Plant Physiol. 128, 201–211 (2002).

6. G. V. Novikova et al., Life 10, 15 (2020).

7. C. Ju, et al., Proc. Natl. Acad. Sci. U.S.A. 109, 19486–19491 (2012).

8. H. Qiao et al., Science 338, 390–393 (2012).

9. X. Wen et al., Cell Res. 22, 1613–1616 (2012).

10. W. Li et al., Cell 163, 670–683 (2015).

11. A. E. Hall, J. L. Findell, G. E. Schaller, E. C. Sisler, A. B. Bleecker, Plant Physiol. 123, 1449–1458 (2000).

12. P. M. Bleeker, H. W. J. Hakvoort, M. Bliek, E. Souer, H. Schat, Plant J. 45, 917–929 (2006).

13. D. H. Wang, F. Li, Q. H. Duan, T. Han, Z. H. Xu, S. N. Bai, Plant J. 61, 862–872 (2010).

14. H. Jin, et al., Sci. Adv. 6, eaba3418 (2020).

15. Q. Liu, C. Xu, C. K. Wen, BMC Plant Biol. 10, 60 (2010).

16. Q. Liu, C. K. Wen, Plant Physiol. 158, 1193–1207 (2012).

17. M. Zhang, et al., Plant Biotechnol. J. 20, 1435–1437 (2022).

18. A. F. Bent, T. K. Hoffman, J. S. Schmidt, G. L. Hartman, D. D. Hoffman, P. Xue, M. L. Tucker, Crop Sci. 46, 893–901 (2006).

19. B. K. Pandey et al., Science 371, 276–280 (2021).

20. Z. Shao et al., New Phytol. 236, 1762–1778 (2022).

21. Y. Zhou et al., Plant Cell, koac250 (2022).

22. W. Li, et al., Mol. Plant 15, 354–362 (2022).

23. H. L. Park, et al., Nat. Commun. 14, 365 (2023).

24. P. Suchomelová-Mašková, O. Novák, H. Lipavská, Plant Physiol. Biochem. 46, 673–684 (2008).

25. H. Lipavská, P. Masková, P. Vojvodová, Ann. Bot. 107, 1071–1086 (2011).

26. X. K. Li, et al., Nat. Commun. 15, 5987 (2024).

27. H. Zhao et al., Plant Cell 32, 1626–1643 (2020).

28. A. B. Bleecker, H. Kende, Annu. Rev. Cell Dev. Biol. 16, 1–18 (2000).

