## Supplemental Figure 1-14 for "The cleaved C-terminal domain of GmERS1-1 modulates cell cycle via GmACR1"

Kai Chen et al.

**The PDF file includes:**

Materials and Methods

Supplementary Text

Figs. S1 to S14

Movie S1 to S3

Tables S1 to S5

References

Other Supplementary Material for this manuscript includes the following:

The gene and protein sequence information related to this study.

**Materials and Methods**

**Phylogenetic Tree Construction**

The coding sequence of *GmERS1-1* (Glyma.03G216700) was retrieved from Phytozome (https://phytozome-next.jgi.doe.gov/). BLAST searches were performed against the *Glycine max* Wm82.a6.v1 reference genome to identify homologs, yielding 11 ethylene receptor protein sequences in soybean. These sequences, together with 5 ethylene receptor proteins from *Arabidopsis thaliana*, were compiled into a single FASTA file. The FASTA file was imported into MEGA software via File → Open A File/Session. Sequence integrity was verified by checking length and naming consistency; font settings were adjusted via File → Font. Multiple sequence alignment was conducted using Alignment → Align by ClustalW or Align by MUSCLE. The aligned dataset was saved as a .meg file via Data → Save Session. Phylogenetic trees were constructed using the neighbor-joining (NJ) method under Phylogeny → Construct/Test Neighbor-Joining Tree. Parameters were set as follows: Test of Phylogeny = Bootstrap method with 1000 replicates; Substitution Model = p-distance; all other parameters were kept as default (*1*).

**Heatmap Preparation**

Expression profiles of 11 soybean ethylene receptor genes were downloaded from SoyBase (https://soybase.org). Heatmaps were generated using HemI 1.0 (Heatmap Illustrator; http://www.hemi.biocuckoo.org/). Briefly, data were loaded via File → Load (.xls or .csv format); for Excel files, the corresponding worksheet was selected and confirmed. Data ranges were defined manually or via Auto select, followed by Finish. Color schemes (e.g., red–white–blue) and custom gradient thresholds were set under the Set tab. Clustering of rows (genes) and/or columns (samples) was applied to enhance pattern recognition. Axis labels, font sizes, and image rotation were adjusted; the plot was refreshed via REFRESH after each parameter change. Final images were exported at 300 dpi via File → Export Image (*2*).

**GUS Staining**

Vector Construction: A 2000-bp genomic fragment upstream of the *GmERS1-1* ATG start codon was amplified and inserted into the *Eco*RI site of pCAMBIA1305 to generate the *GmERS1-1*pro::GUS reporter construct.

Agrobacterium-Mediated Transformation: The construct was introduced into *Agrobacterium tumefaciens* strains GV3101(pSoup-p19) (ZOMANBIO, ZK3718, Beijing) and K599 (ZOMANBIO, ZK3720, Beijing) by chemical competence.

Arabidopsis Transformation (GV3101): Arabidopsis plants were well-watered 1 day before infiltration. A 1-mL overnight bacterial culture (YEP + Kan⁺/Rif⁺, 28 °C, 200–250 rpm, OD₆₀₀ > 2.0) was harvested by centrifugation at 4000–6000 rpm for 10 min. The pellet was resuspended in 3 volumes of infiltration medium (½ MS, 50 g/L sucrose, 0.5 g/L MES, pH 5.7–5.8, 2.7 g/L MgCl₂·6H₂O) to OD₆₀₀ ≈ 0.8. Silwet L-77 was added to 0.02% (15 μL per 100 mL) immediately before use. Floral dip infiltration was performed for 50 s to 5 min. Plants were covered with a black plastic bag for 5–10 min, infiltrated a second time, then placed horizontally in the dark for 24 h. A second infiltration was performed 1 week later if needed.

Soybean Hairy Root Transformation (K599): Bacteria were streaked onto YEP + Kan⁺/Str⁺ solid medium for propagation. Uniform soybean seeds were germinated; seedlings with unfolded cotyledons were inoculated by puncturing the hypocotyl below the cotyledonary node with a needle loaded with bacteria, avoiding damage to the shoot apical meristem and true leaves. Seedlings were covered with a 15–20-cm transparent cup and kept in high humidity and darkness for 24 h. After 2–3 days of growth, the cup bottom was removed, and moist soil was added to cover the wound. Plants were watered from the bottom, and additional soil was supplied as hairy roots emerged.

GUS Staining Protocol (Kit RTU4032, Real-Times, Beijing): **Tissue preparation**: Tissues (leaves, petals, roots, stems) were sliced into small pieces (1–3 mm for thick organs). Vacuum infiltration was used for large tissues to facilitate substrate penetration. **Staining**: Samples were submerged in GUS staining solution, wrapped in foil, and incubated on a shaker until blue color appeared. Negative and positive controls were included. **Destaining**: Tissues were transferred to absolute ethanol or 80% acetone and decolorized 2–3 times until control tissues were white. **Observation**: Stained tissues were examined under a stereomicroscope or light microscope (*3*).

**Generation of Transgenic Soybean Plants**

Overexpression Lines: Specific primers containing *Nco*I and *Bs*TEII sites were designed to amplify *GmERS1-1* from soybean cDNA. The PCR product was purified and double-digested with *Nco*I and *Bs*TEII, then ligated into the similarly digested p3301 vector. The ligation product was transformed into *Escherichia coli* Trans10 competent cells. Positive clones were confirmed by sequencing to generate p3301-*GmERS1-1*. The recombinant plasmid was introduced into *A. tumefaciens* EHA105 by the freeze–thaw method. Cotyledonary nodes of wild-type soybean W82 were used as explants for *Agrobacterium*-mediated transformation to generate T₀ transgenic plants. Positive T₀ lines were screened by PCR and Bar strip test, then selfed to homozygosity. Stable T₃ homozygous overexpression lines were used for experiments.

EMS Mutant Lines: Wild-type W82 seeds were soaked in deionized water, then mutagenized with 0.04 mol/L EMS (pH 7.0, phosphate buffer) for 6 h. Seeds were thoroughly rinsed and air-dried. The M₁ generation was planted at high density and harvested individually. M₂ families were grown and screened for morphological variants (leaf shape, stem, flower, pod). M₃–M₄ lines were analyzed for seed nutritional quality by near-infrared spectroscopy. Targeted point mutations were identified by TILLING combined with CelⅠ digestion. A stable homozygous M₅ mutant (*gmers1-1#2*) carrying a G→A substitution (Q→Stop, premature termination) was confirmed by DNA sequencing. Loss of C-terminal protein expression was verified by Western blot. Agronomic traits (yield, plant height, pods per plant) were scored at maturity (*4*).

**Mass Spectrometry Analysis**

In-Gel Digestion: Target protein bands were excised, diced, and washed twice with distilled water. Gel pieces were destained with 25 mM NH₄HCO₃ / 50% acetonitrile (ACN) at 37 °C until clear, then dehydrated with ACN and dried by vacuum centrifugation. Gel pieces were reduced with 25 mM dithiothreitol at 55 °C for 45 min, alkylated with 55 mM iodoacetamide in the dark for 30 min, then washed and dried again. Trypsin solution (10 μg in 790 μL buffer: 200 mM NH₄HCO₃, ACN, water) was added to cover gel pieces, followed by overnight digestion at 37 °C. Digestion was stopped with 10% trifluoroacetic acid (TFA). Peptides were extracted twice with 50% ACN/0.1% formic acid (FA) at 37 °C for 30 min. Extracts were combined, concentrated to ~10 μL, and reconstituted in 0.1% FA twice.

LC-MS/MS Analysis: Peptides were separated on an Easy nLC 1000 system using mobile phases A (0.1% FA in water) and B (0.1% FA in ACN). Flow rate = 0.4 μL/min; gradient: 3–6% B (3 min), 6–22% B (78 min), 22–100% B (1 min), 100% B (8 min). Mass spectrometry was performed on a Q Exactive Plus Orbitrap mass spectrometer in data-dependent acquisition (DDA) mode. MS1 parameters: resolution = 70,000; mass range = 300–1800 m/z; AGC = 3×10⁶; max IT = 50 ms. MS2 parameters: resolution = 17,500; AGC = 1×10⁵; max IT = 45 ms; NCE = 27; top 20 precursors per cycle. Three technical replicates were acquired per sample.

Protein Identification: Raw files were processed with Proteome Discoverer 2.0 using the SEQUEST HT engine against the SoyBase soybean protein database (https://www.soybase.org/). Search parameters: high peptide confidence; max 2 missed cleavages; peptide length = 6–144; precursor mass tolerance = ±10 ppm; fragment mass tolerance = 0.02 Da; fixed modification = carbamidomethylation (+57.021 Da); variable modifications = methionine oxidation (+15.995 Da) and protein N-terminal acetylation (+42.011 Da); FDR = 1% (*5*).

**Soybean Protoplast Isolation and Transfection**

A standardized protocol for high-efficiency soybean mesophyll protoplast isolation and transfection was established. Fully expanded trifoliate leaves from FT14-stage plants were sliced into strips and digested in enzyme solution. Combinations V (1.0% cellulase + 0.2% pectolase) and VI (1.0% cellulase + 0.4% pectolase) yielded the highest viability and yield.

In a 220-μL reaction, 10⁴–10⁵ protoplasts were mixed with 10 μg plasmid DNA and 20% PEG4000, then incubated at 25 °C in the dark for 15 min. Transfection efficiency routinely exceeded 80%. Transfected protoplasts were cultured in the dark and used for subcellular localization, protein colocalization, and other functional assays (*6*).

**Nuclear and Membrane Protein Fractionation**

Membrane and nuclear fractions were isolated from 3-day-old dark-grown *Arabidopsis* Col-0 seedlings treated with or without ethylene. For membrane extraction, 1 g tissue was homogenized in cold buffer (30 mM Tris pH 7.4, 150 mM NaCl, 10 mM EDTA, 20% glycerol, protease inhibitor cocktail). Homogenates were filtered through miracloth and centrifuged at 10,000 g for 5 min. The supernatant was ultracentrifuged at 100,000 g for 30 min to pellet membranes.

For nuclei isolation, 1 g tissue was homogenized in extraction buffer (20 mM PIPES-KOH pH 7.0, 4 M hexylene glycol, 10 mM MgCl₂, 0.25% Triton X-100, 4 mM 2-mercaptoethanol, protease inhibitor cocktail). The lysate was filtered and loaded onto a 30%/80% Percoll gradient, then centrifuged at 2000 g for 30 min. Nuclei were collected from the 30–80% interface.

Protein samples were separated by SDS-PAGE and analyzed by Western blotting. EIN2 was detected with anti-EIN2 antibodies. ACA2 (plasma membrane Ca²⁺ pump) and histone H3 were used as fraction-specific markers (*7*).

**Transmission Electron Microscopy**

Fixed tissues were washed 5 times with 0.1 M PBS (pH 7.4) for 5–10 min each, then post-fixed with 1% osmium tetroxide at 4 °C for 2 h and washed again. Samples were dehydrated in an acetone series (50%, 70%, 90%, 100%, 100%) at 4 °C for 15 min each, then infiltrated with gradient resin–acetone mixtures (50%, 70%) for 2 h each, followed by two changes of pure resin for 8 h each. Samples were embedded in pure resin in molds and polymerized stepwise: 37 °C for 12 h, 45 °C for 12 h, 60 °C for 24 h. Semithin sections (300 nm) were stained with 1% toluidine blue for light microscopy screening. Ultrathin sections (70 nm) were cut at 3–5 mm/s using a diamond knife on an EM UC7 ultramicrotome. Sections were stained with uranyl acetate (30 min, dark) and lead citrate (5–10 min, CO₂ free), then observed under an HT7700 transmission electron microscope (*8*).

**Flow Cytometric Analysis of Root Tip Ploidy**

All steps were performed at 4 °C to prevent DNA degradation. Fresh root tips (0.5–1 cm) were digested in enzyme solution (1.5% cellulase, 0.3% pectinase, 0.4 M mannitol, pH 5.7–6.0) at 28 °C in the dark for 1–2 h. The cell suspension was filtered through a 40-μm mesh and centrifuged at 200×g for 5 min. For staining: DAPI (1 μg/mL) was used directly without permeabilization. PI staining required RNase A (50 μg/mL, 37 °C, 30 min) and 0.1% Triton X-100 permeabilization. The flow cytometer was calibrated with standard beads to CV < 5%. DAPI was excited at 355 nm; PI at 488 nm. Doublets were excluded by pulse gating. Ploidy was assigned using soybean 4C peaks as reference; endoreduplicated peaks (8C, 16C) were validated by chromosome counting (*9*).

**Soybean Root Tip Immunofluorescence**

Fixation and Sample Preparation: Tissues were fixed in solution under vacuum for 60 min at room temperature, then washed twice with 1× PBS. For hard tissues (hypocotyl, cotyledons), dewaxing and chlorophyll removal were performed with methanol and ethanol–xylene series, followed by gradient rehydration. Samples were mounted on slides: Root tips: air-dried in water droplets. Ovules/embryos: gently squashed to release embryos, then air-dried. Hypocotyl/leaves: frozen in liquid nitrogen 5× to permeabilize, then air-dried. Slides were stored at −20 °C. Permeabilization: Slides were rehydrated in PBS, then treated with 2% driselase at 37 °C for 30–60 min, washed 5× with PBS, incubated with 3% Triton X-100 + 10% DMSO for 60 min, then washed 5–10× with PBS. Blocking and Antibody Incubation: Samples were blocked with 3% BSA for 60 min, then incubated with primary antibody (100–150 μL) at 37 °C for ≥4 h or 4 °C overnight. After 5× PBS washes, secondary antibody was applied and incubated at 37 °C in the dark for ≥3 h. Staining and Imaging: Nuclei were stained with DAPI (1:1000) for 30 min if required. Slides were washed, mounted in anti-fade solution, and observed under a fluorescence microscope (*10*).

**RNA Purification and qRT-PCR**

Total RNA was extracted using TRIzol reagent (TaKaRa), treated with RNase-free DNase I (TaKaRa), quantified spectrophotometrically, and verified by 1.5% denaturing agarose gel electrophoresis. cDNA was synthesized from 3 μg RNA using oligo-dT(18), dNTPs, and M-MLV reverse transcriptase (TaKaRa). qRT-PCR was performed as described (*11*) with four technical replicates per sample. Primer sequences are listed in Table S1 (*11*).

**Subcellular Localization**

Full-length *GmERS1-1* cDNA was amplified and cloned into p16318hGFP to generate N- or C-terminal GFP fusions driven by the double CaMV 35S promoter. Nuclear markers were generated by fusing NLS sequences to GFP/RFP or by DAPI staining. HDEL-RFP was used as an ER marker. Fusion constructs were transiently expressed in soybean protoplasts as described (*6*). Fluorescence was observed under a Zeiss LSM780 confocal microscope.

**Yeast Two-Hybrid (Y2H) Assay**

The *GmERS1-1* coding sequence and potential interactor *GmACR1* were cloned into pBT3-N (bait) and pPR3-N (prey), respectively. Constructs were co-transformed into Y2H Gold yeast cells and selected on SD/−Leu/−Trp medium at 30 °C for 3 days. Interaction was tested on SD/−Leu/−Trp/−Ade/−His medium supplemented with 10 mM 3-AT or 4 mg/mL X-gal. β-galactosidase activity was assayed by filter lift and liquid culture using X-gal or o-nitrophenylgalactoside as substrate. (*11*) Primer sequences are listed in Table S1.

**Bimolecular Fluorescence Complementation (BiFC) Assay**

Full-length *GmERS1-1*, *GmERTS1-1* N-terminal, and *GmERTS1-1* C-terminal fragments were cloned into pSAT1-nEYFP. *GmACR1* was cloned into pSAT1-cEYFP. Constructs were co-transfected into soybean protoplasts with nuclear (NLS-GFP/RFP or DAPI) and ER (HDEL-RFP) markers. Protoplasts were incubated at 22 °C in darkness for 16 h. YFP fluorescence (Ex 514 nm, Em 527 nm) was observed using a Zeiss LSM780 Meta confocal microscope (*11*).

**Co-Immunoprecipitation (Co-IP)**

GFP-*GmERTS1-1* and FLAG-*GmACR1* constructs were transformed into *A. tumefaciens* GV3101 and infiltrated into 4–6-week-old *Nicotiana benthamiana* leaves. After 48 h, tissues were ground in liquid nitrogen and extracted in cold buffer (200 mM MOPS pH 7.0, 5% PVPP, 1% Triton X-100, 10% glycerol, 2 mM DTT, protease/phosphatase inhibitors). Extracts were centrifuged at 20,000 g for 20 min at 4 °C. Supernatants were incubated with anti-GFP, anti-FLAG, or control IgG antibodies for 1 h, then with protein A/G beads for 2 h at 4 °C. Beads were washed three times, boiled in loading buffer, and analyzed by SDS-PAGE and Western blotting (*11*).

**Overexpression in *Schizosaccharomyces pombe***

The *GmERS1-1*，*GmACR1* and *GmCDK7* coding sequence was amplified using primers in **Table S1**, cloned into pREP1, and verified by sequencing. Overexpression was induced in *S. pombe* under thiamine-depleted conditions as described (*12*).

**Gene Ontology Biological Process enrichment analysis and network visualization**

Perform Gene Ontology (GO) Biological Process enrichment analysis on the gene GmERS1-1, and visualize the enrichment results. Highlight the term Regulation of stem cell division in red in the enrichment plot. Construct a protein interaction network for GmERS1-1 to display functional enrichment information of associated proteins; mark the four proteins annotated to Regulation of stem cell division with red color, and draw a red dashed line to show the interaction connection between target protein GmACR1 and GmERS1-1.

**Gene family classification of GmACR1**

Query the gene family information of GmACR1 via the LegumeInfo Funnotate database (https://funnotate.legumeinfo.org/?family=legfed_v1_0.L_M6P16H). Mark the gene locus of GmACR1 with a red box in the gene family classification diagram, and set a legend with dots of distinct colors to distinguish different species sources.

**Conserved domain prediction and protein homology modeling**

Conserved domain prediction: Submit the protein sequences of GmACR1, Homo sapiens CDC25B, and Arabidopsis thaliana AtACR1 (AT5G03455) to the SMART online tool (https://smart.embl.de/) for structural domain prediction. Label the amino acid position scale on the schematic, and use purple blocks to annotate Rhodanese Homology Domain (RHOD). Homology modeling: Use SWISS-MODEL to construct three-dimensional homology models for the above three full-length proteins. For human CDC25B, an independent structural model is additionally constructed for its C-terminal fragment spanning amino acids 393–580; amino acid position numbers are labeled on all protein models.

**Prediction of Protein–Protein Interaction Complex Structures**

Three-dimensional structures of bait proteins were constructed using AlphaFold3 (*13*, *14*). Protein–protein docking was performed via HDOCK: protein GmERS1-1 C’ was first docked with protein GmACR1, followed by docking of the obtained GmERS1-1 C’–GmACR1 complex with protein GmCDK7. Output files were sorted, and the 3D structure with the second-highest confidence score predicted by AlphaFold3 was selected for visualization using PyMOL.

**Figure S1**

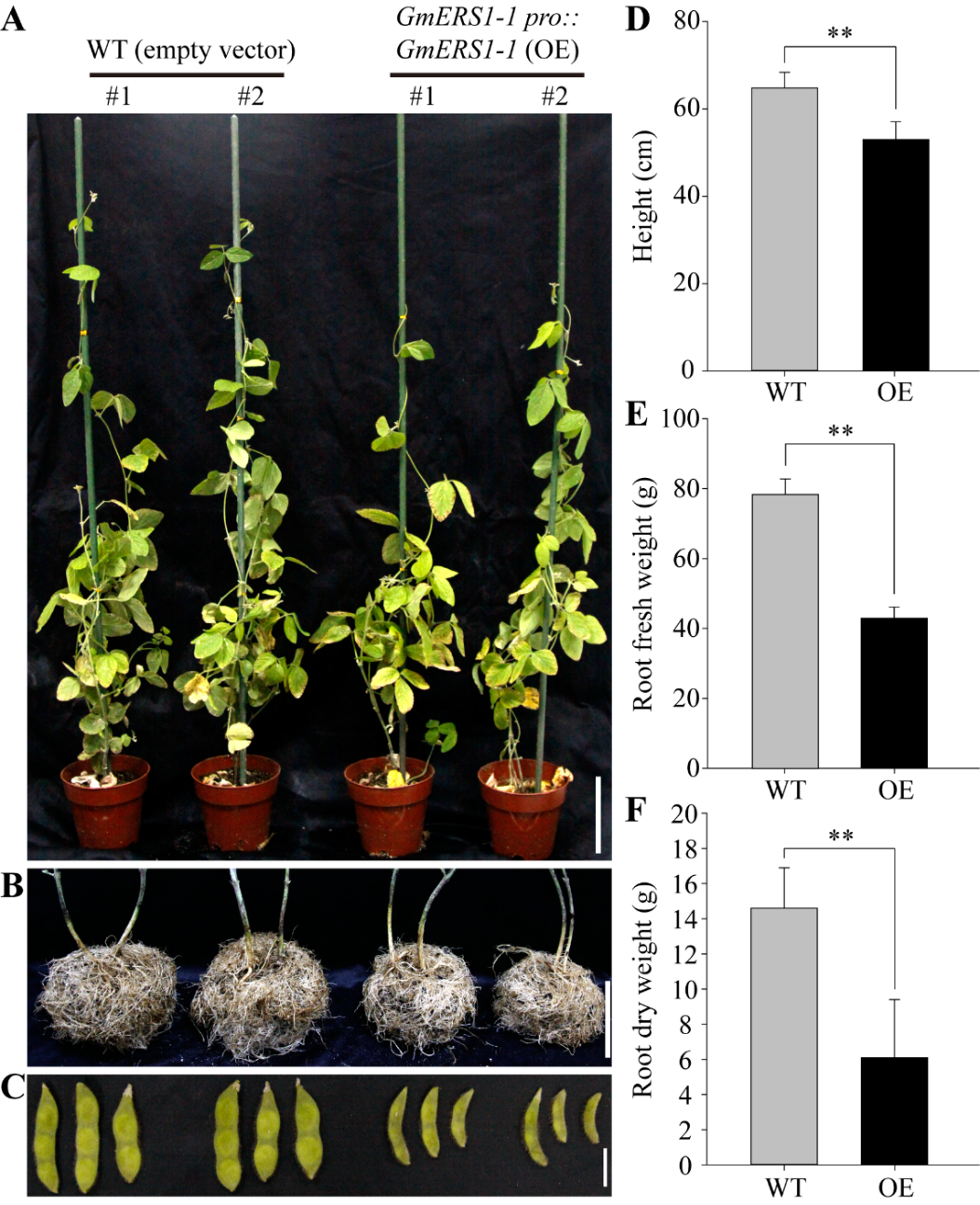
**Figure S1. Overexpression of *GmERS1-1* inhibits the growth of soybean.** The soybean hairy root transformation system was used to verify the functions of *GmERS1-1* in soybean. **(A)** Potted phenotype. **(B)** Root morphology. **(C)** Pods phenotype. **(D)** Plant height, **(E)** fresh weight of roots, **(F)** dry weight of roots were analyzed, and the error line represents SE. Statistical analysis was conducted using Student’s *t*-test. Asterisks indicate comparison with the control group (**, *P*<0.01). The results showed that overexpression of *GmERS1-1* significantly inhibited the growth of soybean.

**Figure S2**

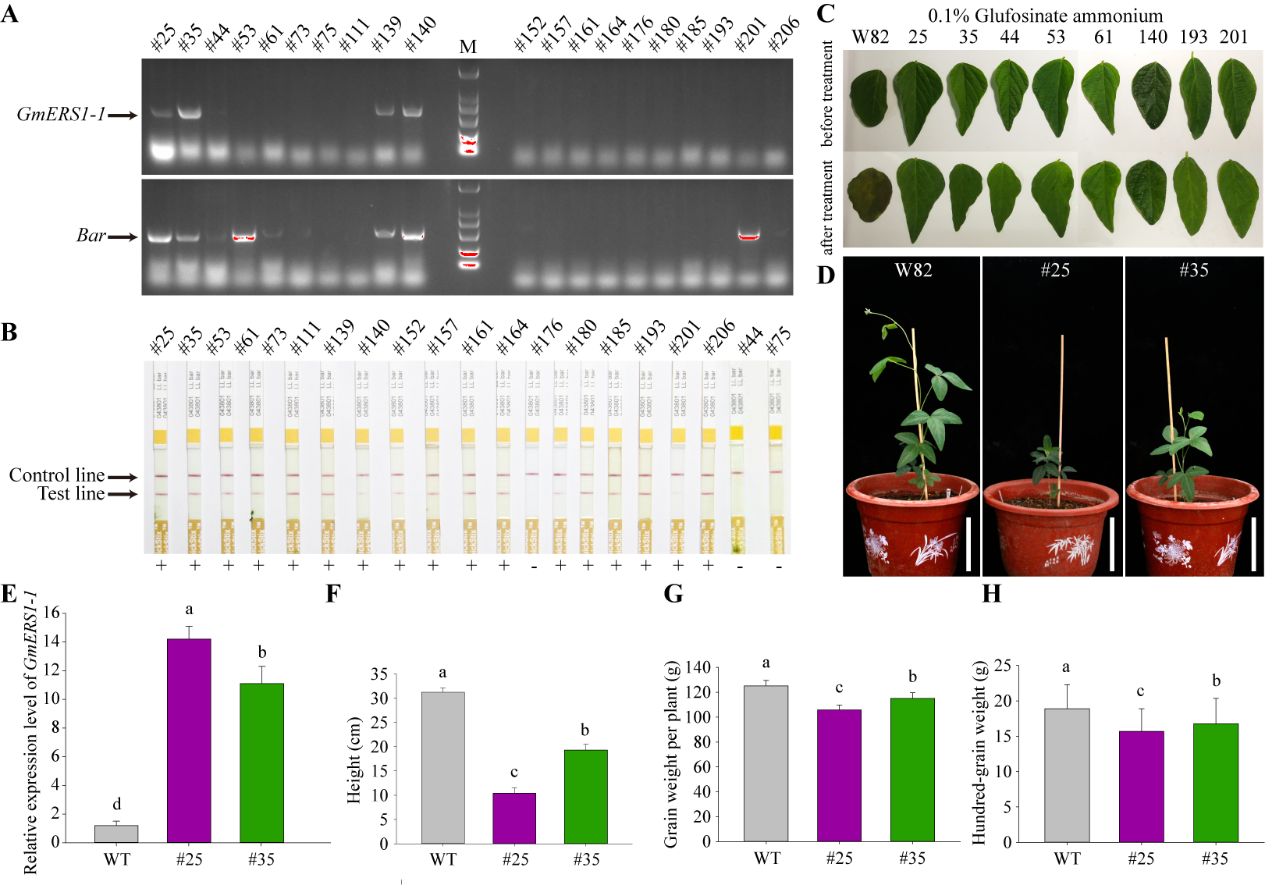

**Figure S2. Molecular identification and seedling phenotype of *GmERS1-1* transgenic soybean. (A)** PCR detection of specific fragments of *GmERS1-1* gene and *Bar* gene in transgenic plants. **(B)** Test strip detection for *Bar* gene: the upper band is the control band, the lower band is the experimental band, and plants with two bands are positive transgenic plants, while only one band is no transgenic plants. Spray 0.1% glyphosate on leaves **(C)** of transgenic lines to detect *Bar* gene activity during seedling stage, and wild type soybean W82 showed sensitivity to herbicides, while transgenic soybeans showed resistance to herbicides. Finally, transgenic lines # 25 and # 35 with *GmERS1-1* and *Bar* genes, and exhibiting herbicide resistance were selected for further experiments. **(D)** The phenotype of W82 and transgenic lines # 25 and # 35 during the seedling stage was showed. **(E-H)** The expression level of *GmERS1-1* gene in transgenic lines #25 and #35, and plant height, single plant grain weight, and hundred grain weight in both transgenic lines. Data statistics conducted ANOVA-DUNCAN's multiple range tests at *P*<0.05, error bars report SE of the mean, n=3.

**Figure S3**
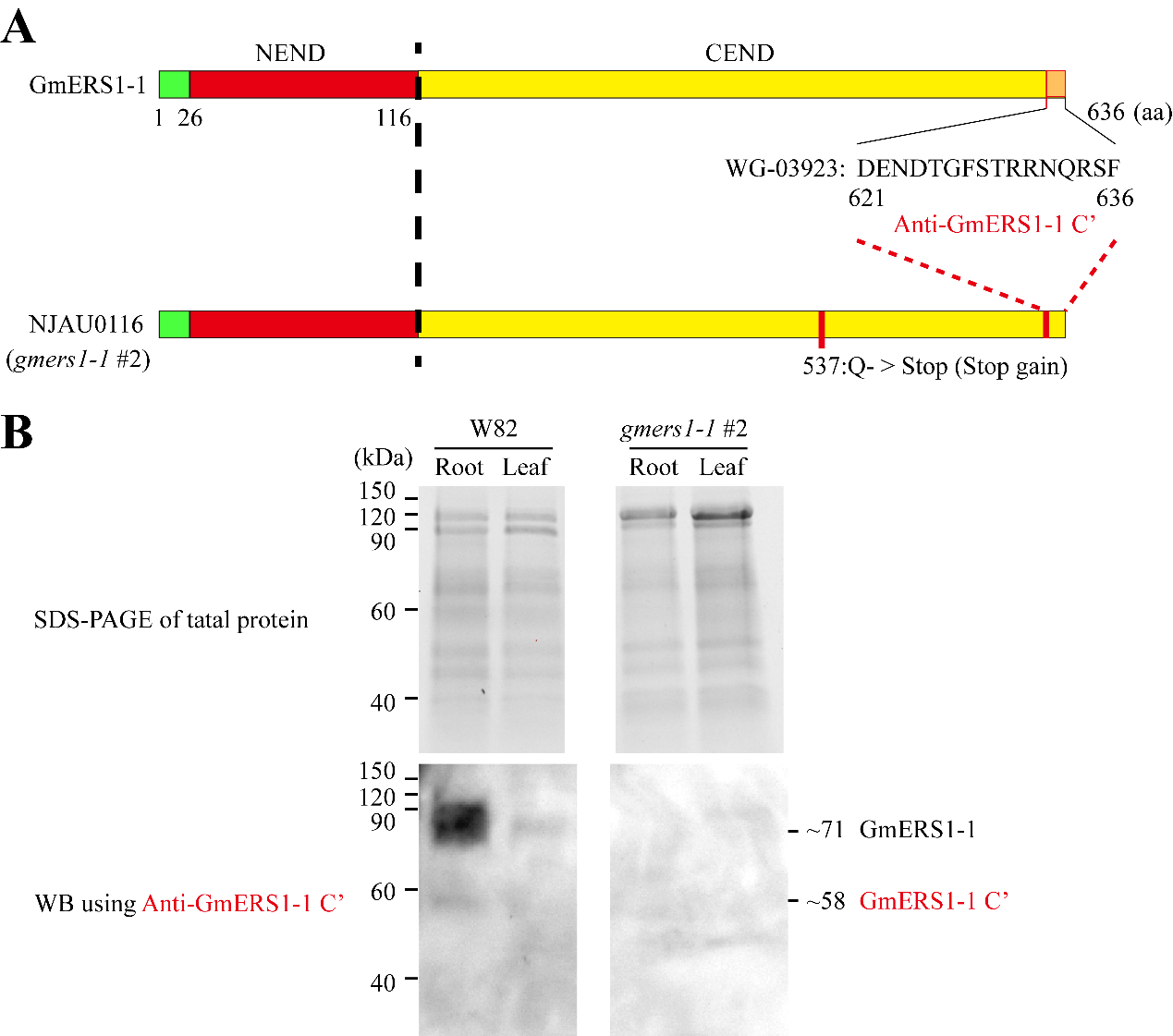

**Figure S3. Detecting mutants of GmERS1-1 by western blot using monoclonal antibody Anti-GmERS1-1 C’.** **(A)** Mouse monoclonal antibody Anti-GmERS1-1 C' (WG-03923) binding sites (621-636 aa.) and mutation sites (537:Q->Stop for mutant2) of *GmERS1-1* mutants. **(B)** The immunoblotting results of GmERS1-1 mutant and wild-type soybean total protein were performed using mouse monoclonal antibody Anti-GmERS1-1 C'.

**Figure S4**
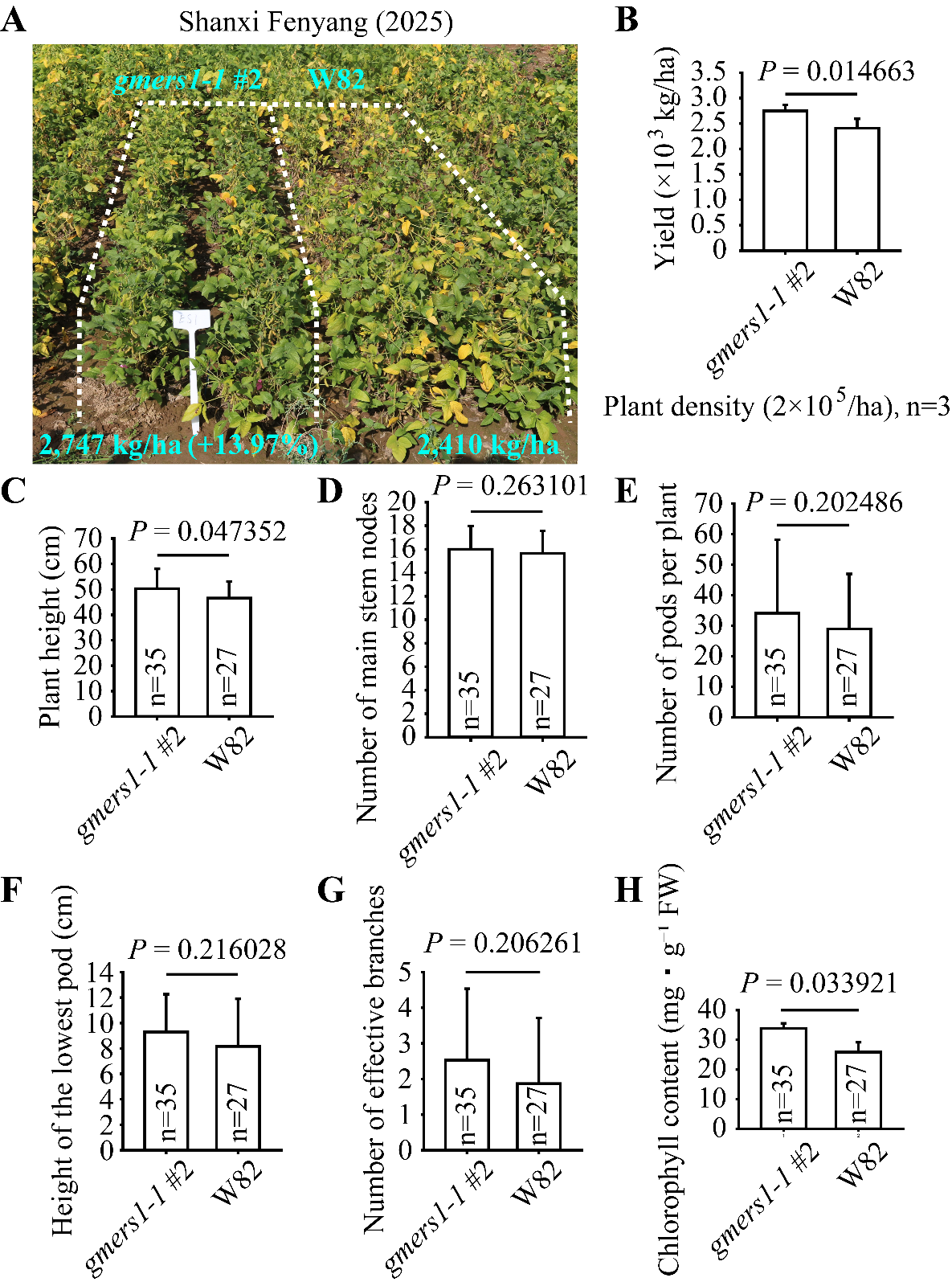

**Figure S4. Field agronomic performance of the *gmers1-1* mutant line #2 and wild-type W82 grown in Fenyang, Shanxi Province (2025 growing season).** **(A)** Representative field phenotype of *gmers1-1* #2 and W82 soybean plants under identical cultivation conditions. Actual grain yields are labeled below each plot; the *gmers1-1* line exhibits a 13.97% yield increase relative to W82. **(B)** Comparison of grain yield per hectare between *gmers1-1* #2 and W82 at a planting density of 2×10⁵ plants ha⁻¹ (n = 3 biological replicates). **(C-H)** Quantitative agronomic and physiological trait measurements for *gmers1-1* #2 (n = 35 individual plants) and W82 (n = 27 individual plants): **(C)** Plant height; **(D)** Number of main stem nodes; **(E)** Number of pods per plant; **(F)** Height of the lowest pod; **(G)** Number of effective branches; **(H)** Leaf chlorophyll concentration (mg g⁻¹ fresh weight, FW). Student’s *t*-test, all *P* values were derived from statistical comparison between the two genotypes; error bars represent standard error of the mean.

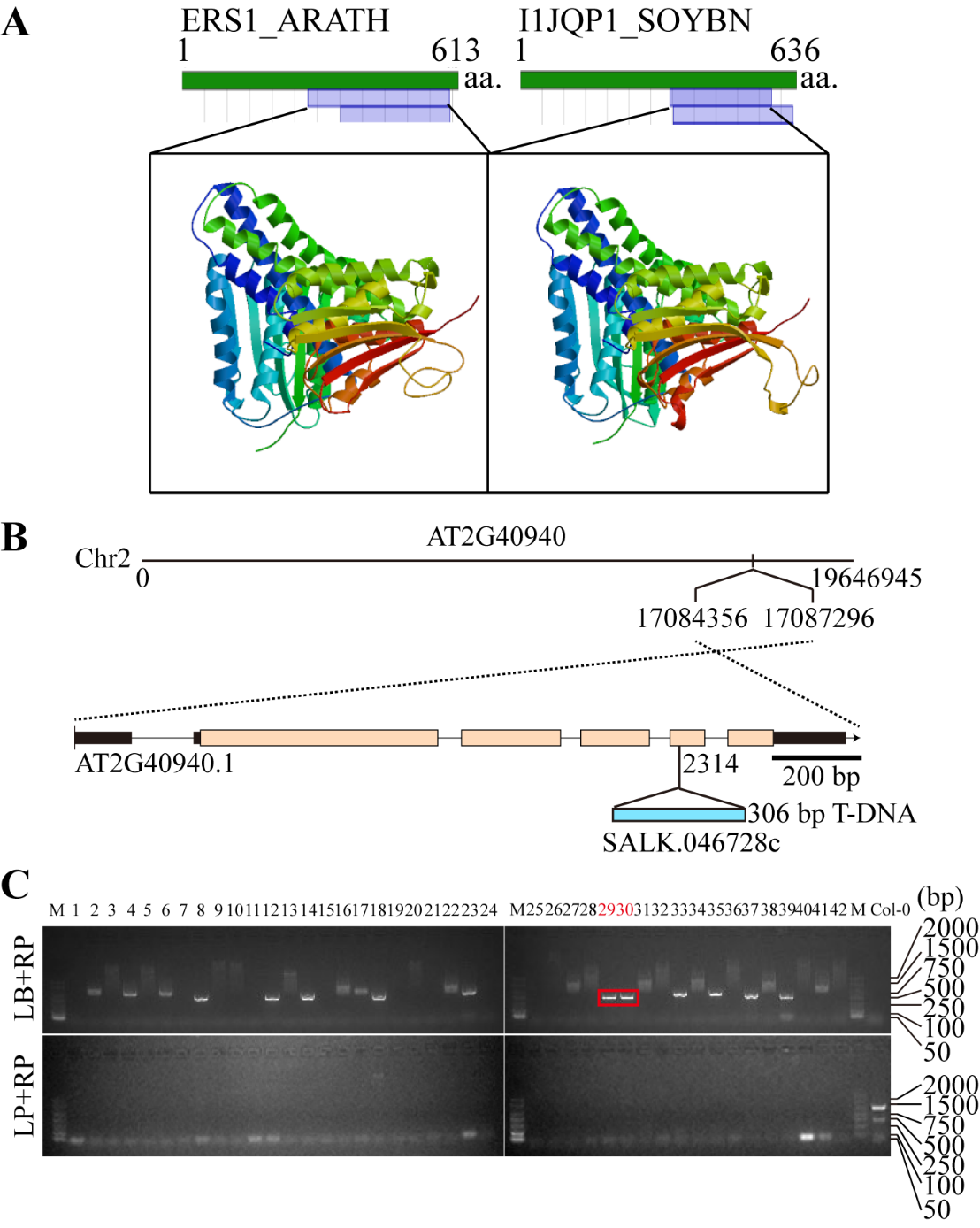
**Figure S5**

**Figure S5. Identification of ethylene receptor mutant *aters1* in *Arabidopsis*. (A)** The C-terminal structures of *Arabidopsis* AtERS1 and soybean GmERS1-1 are highly similar, which indicated that the monocloning antibody Anti-GmERS1-1 C’ (antibody for GmERS1-1 C-terminal) can also be used to detect AtERS1-CEND (C-terminal of AtERS1 in *Arabidopsis*). **(B)** Schematic diagram of T-DNA insertion sites in *Arabidopsis* ethylene receptor *AtERS1* (AT2G40940) in mutant *aters1*. **(C)** The homozygous validation of mutant *aters1* was performed using a dual primer method, and two homozygous lines # 29 and # 30, were selected for further experimental.

**Figure S**
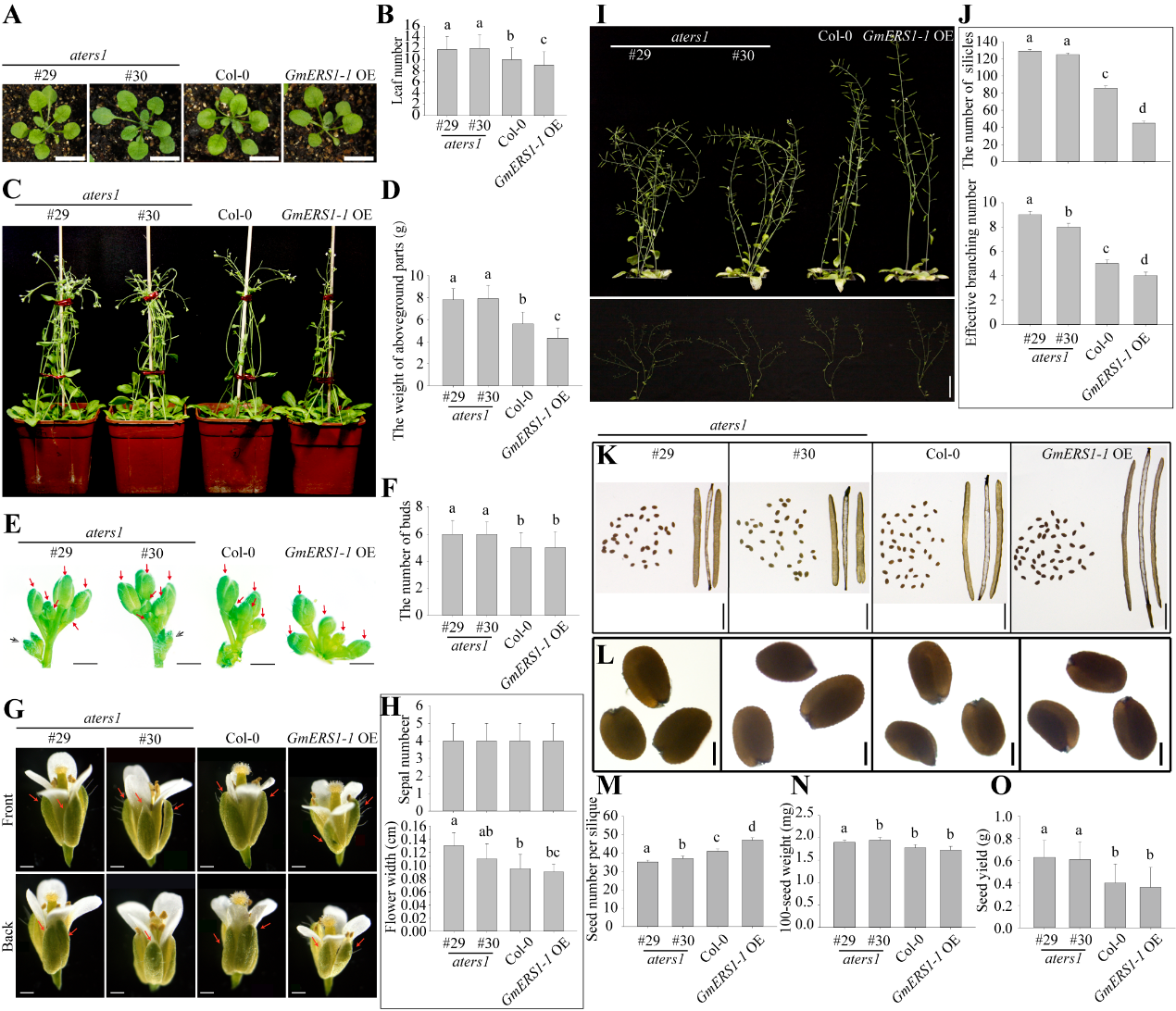
**6**

**Figure S6. Phenotype of *Arabidopsis* *AtERS1* mutants (*****aters1*) and overexpressing *GmERS1-1* (*****GmERS1-1*-OE) in *Arabidopsis* background.** Continuous observation of the entire growth period of the *Arabidopsis* wild-type Col-0, mutant *aters1* (# 29 and # 30), and *GmERS1-1*-OE lines. **(A)** Lotus leaf during seedling stage. **(B)** Leaf number. **(C)** Phenotype of aboveground parts during seedling stage. **(D)** Fresh weight of aboveground parts during seedling stage. **(E)** Apex inflorescence bud. **(F)** Number of apical inflorescence buds. **(G)** Flower morphology during flowering period. **(H)** Sepal number and the width of flower. **(I)** Mature aboveground parts. **(J)** Number of pods, effective number of branches. **(K)** Pod. **(L)** Seed. **(M-O)** Number of seeds per pod, weight of 100 seeds, and seed yield. Data statistics conducted ANOVA-DUNCAN's multiple range tests at *P*<0.05, error bars report SE of the mean, n=3. These results showed that overexpression of *GmERS1-1* inhibited growth and development throughout the entire growth period under *Arabidopsis* background, which is opposite to the phenotype of *aters1* mutant in *Arabidopsis*.

**Figure S7**

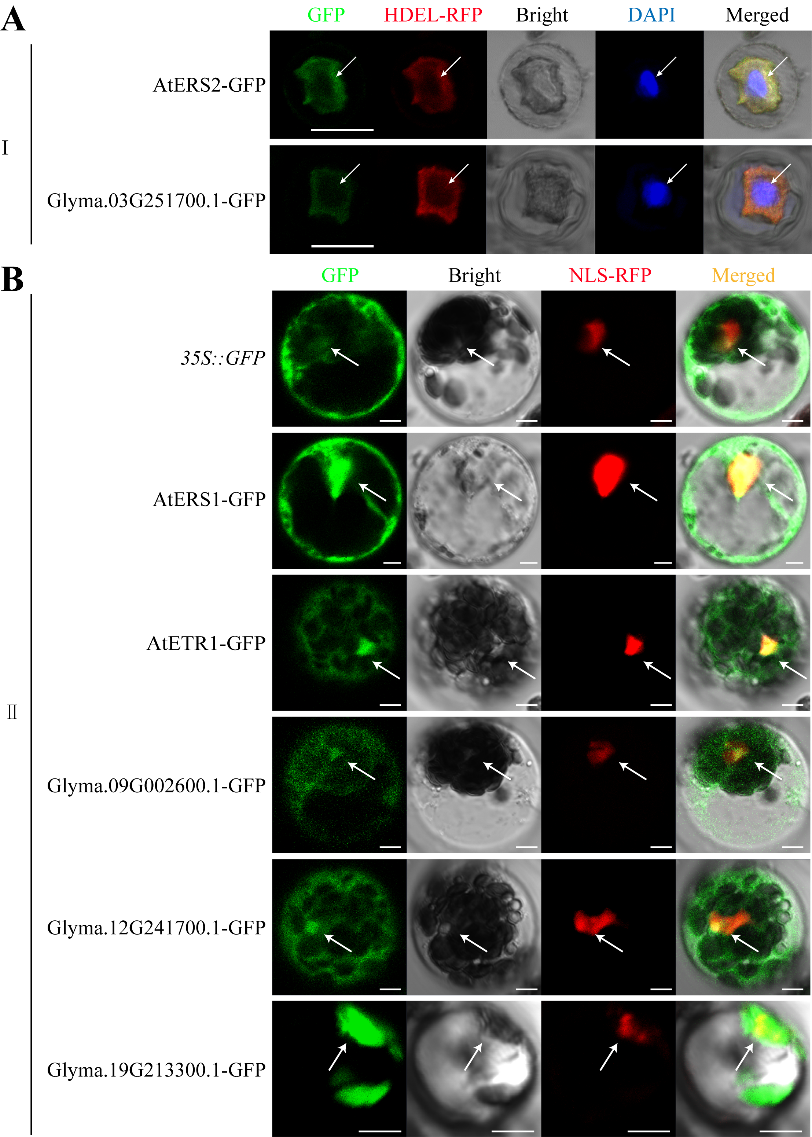

**Figure S7**. **Subcellular localization of the two subfamilies members in ethylene receptor gene family.** The results showed that **(A)** members of the subfamily I containing ERS2 cannot enter the nucleus, **(B)** members of the subfamily Ⅱ containing ERS1 and ETR1 entered the nucleus in *Arabidopsis* and soybean.

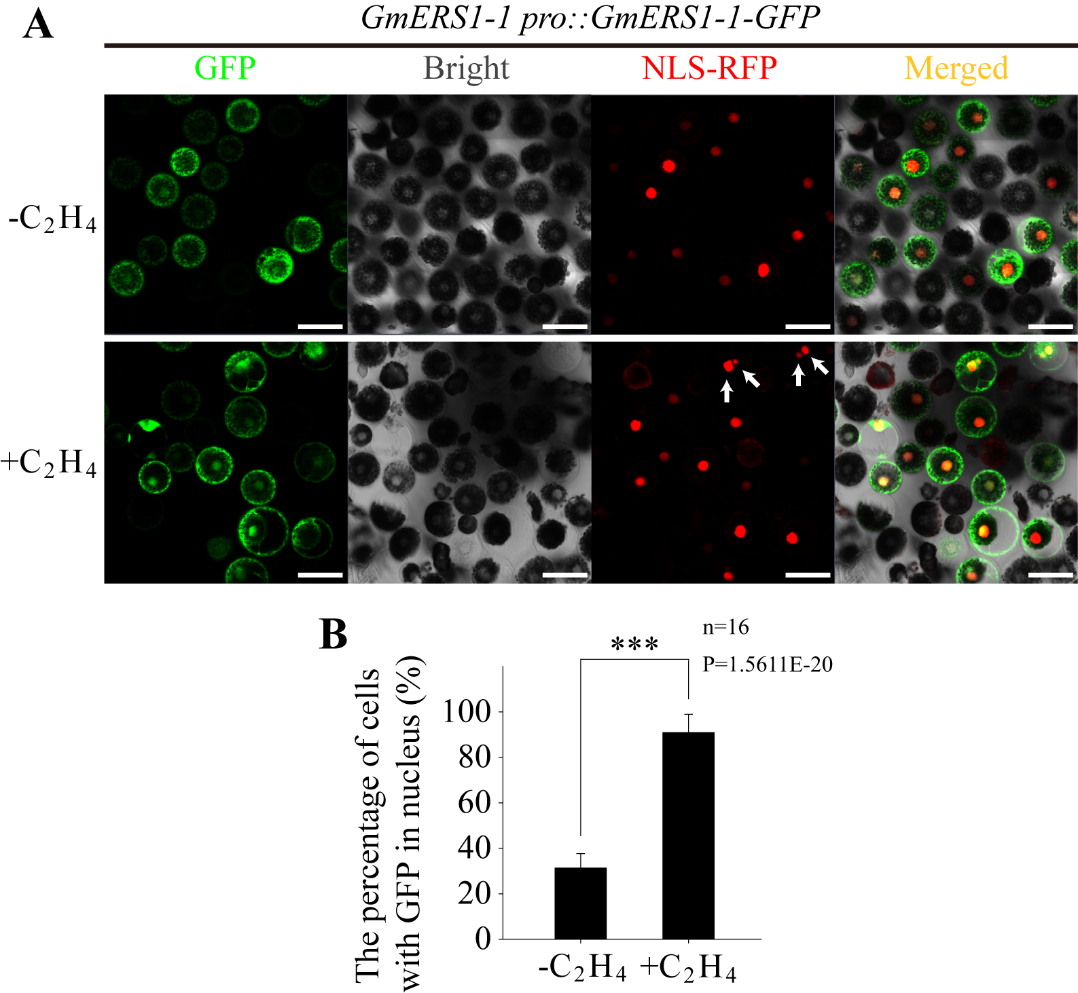
**Figure S8**

**Figure S8. The proportion of cells entering the nucleus of GmERS1-1 C-terminus fused with GFP before and after ethylene treatment. (A)** Transient transformation of soybean protoplasts using *GmERS1-1 pro: GmERS1-1-GFP* vector (n=16). The upper part was untreated and the lower part was transient transformation of soybean protoplasts treated with ethylene. The white arrow points to the binucleated cells that appear after ethylene treatment. **(B)** The proportion of cells with visible fusion protein GmERS1-1-GFP localization in the nucleus. The error line represents SE. Statistical analysis was conducted using Student’s *t*-test. Asterisks indicate comparison with the control group (**, *P*<0.01).

**Figure S9**

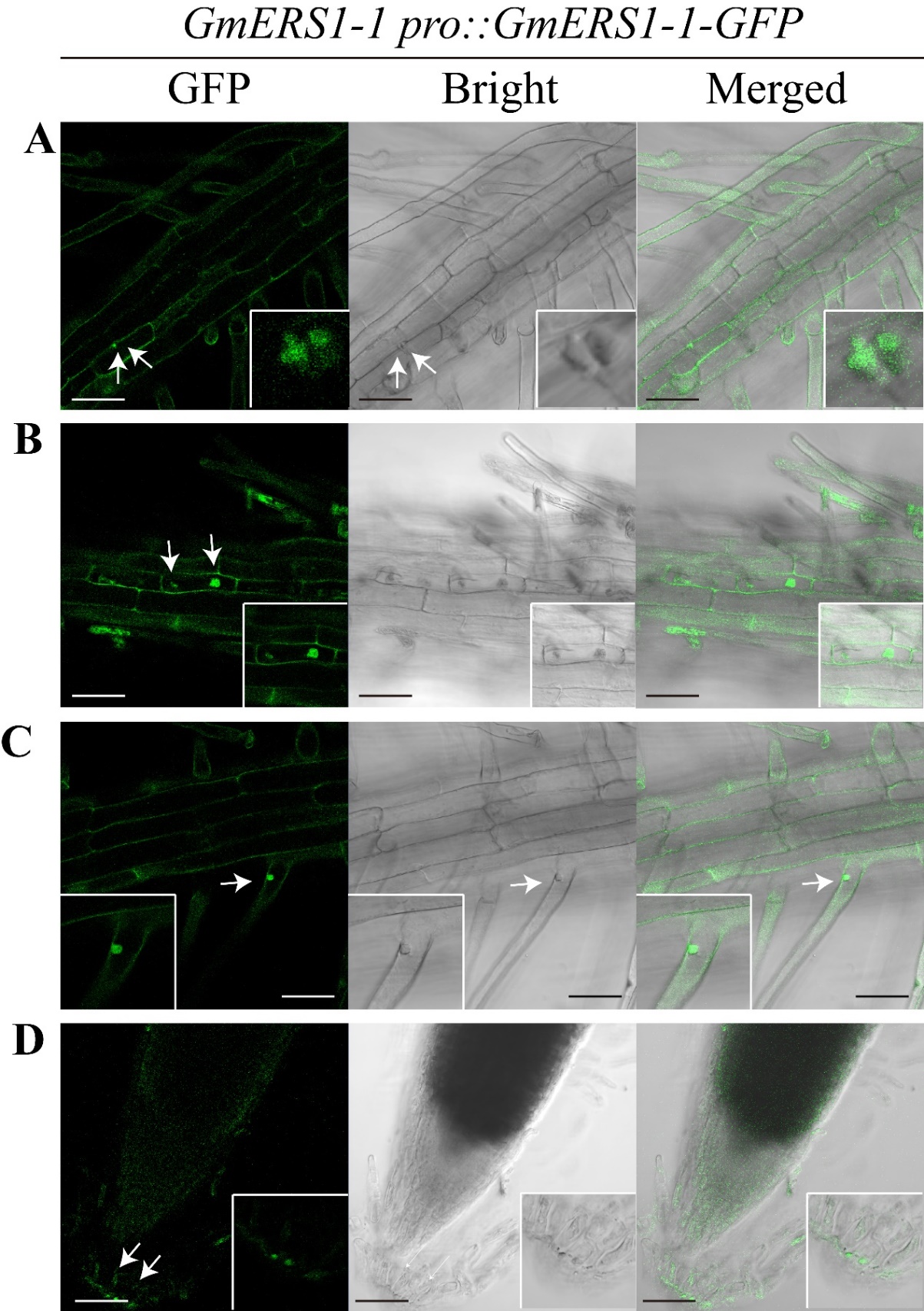

**Figure S9. Subcellular localization of C-terminally GFP-tagged GmERS1-1 in roots of GmERS1-1-overexpressing transgenic Arabidopsis thaliana.**

Fluorescence imaging was performed on transgenic Arabidopsis roots expressing the *GmERS1-1 pro::GmERS1-1-GFP* reporter construct. Each row presents GFP fluorescence (left), brightfield (middle), and merged (right) views for distinct root regions **(A–D)**; insets show magnified views of the marked regions. White arrows indicate nuclear GFP accumulation. Scale bars, 1 μm.

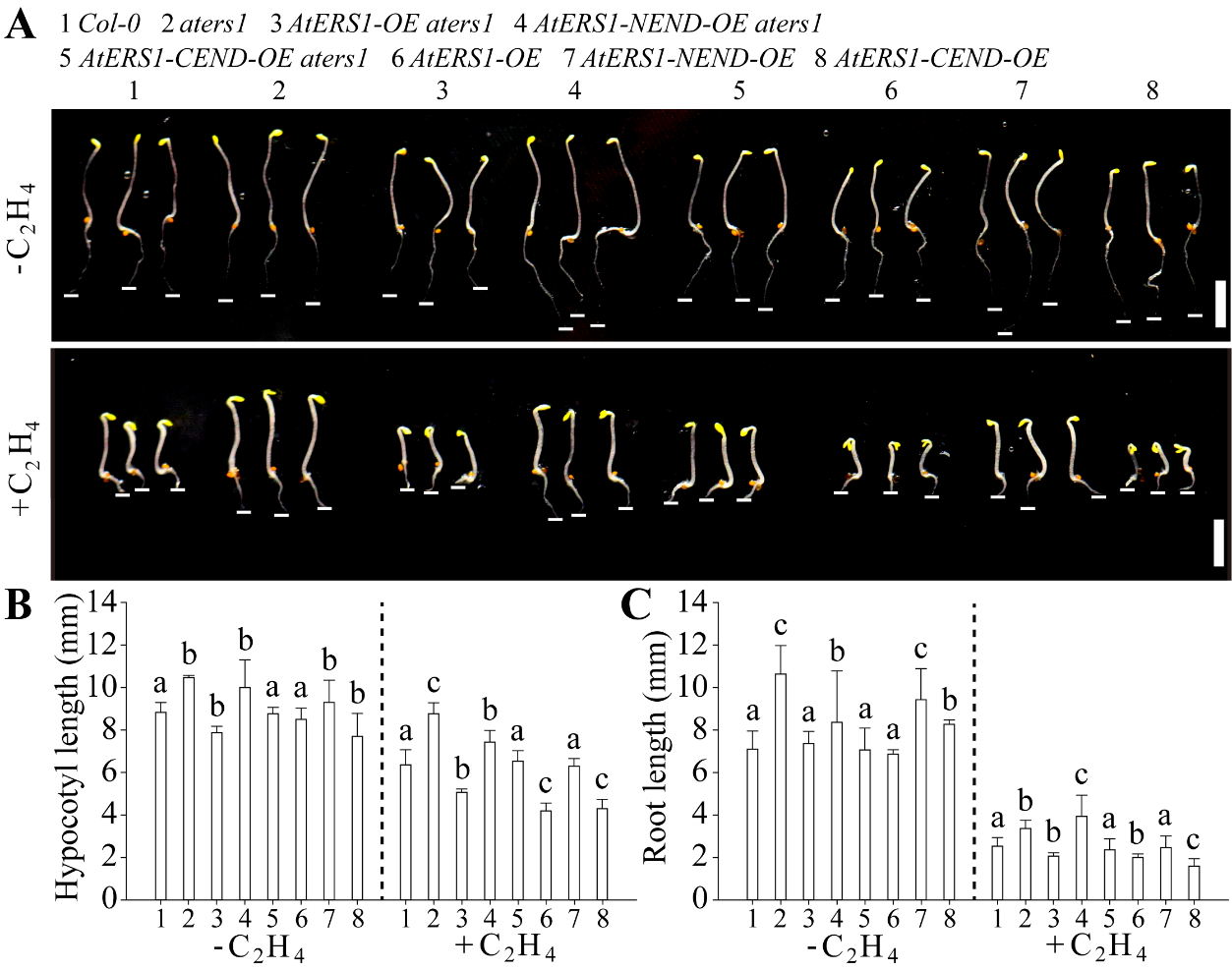
**Figure S10**

**Figure S10. Overexpression of the C-terminus of the ethylene receptor *AtERS1* (*AtERS1-CEND*) can enhance the triple reactions in *Arabidopsis*.** **(A)** full-length *AtERS1*, *AtERS1-NEND* (*the free N-terminus of AtERS1*), *AtERS1-CEND* were overexpressed under *aters1* mutants and wild type *Arabidopsis* background, respectively. The triple responses phenotype of all plants was investigated and the results showed that the full-length *AtERS1* and *AtERS1-CEND* had similar functions and enhanced the triple response, manifested as shortening hypocotyl **(B)** and shortening root length **(C)**. Data statistics conducted ANOVA-DUNCAN's multiple range tests at *P*<0.05, error bars report SE of the mean, n=3.

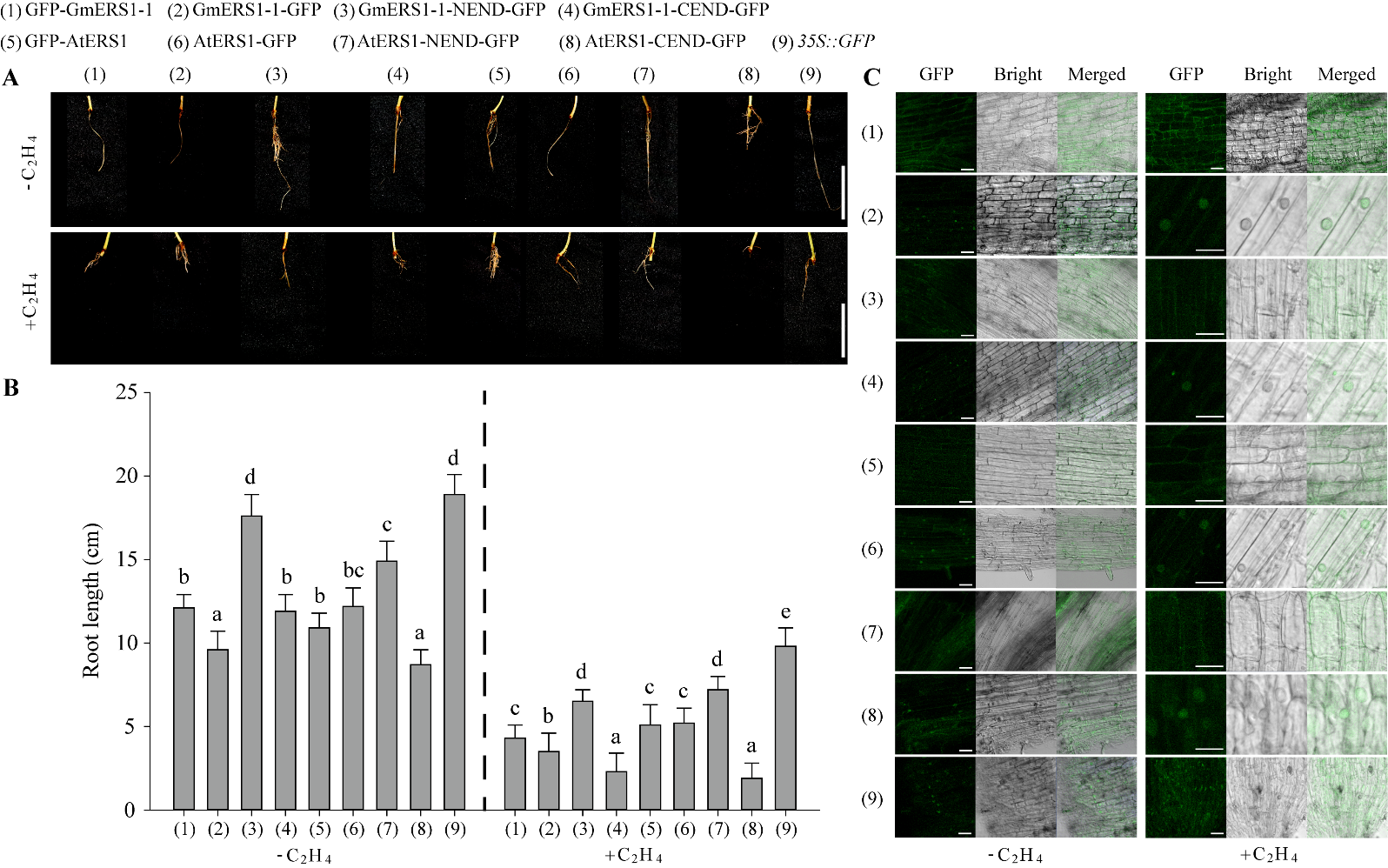
**Figure S11**

**Figure S11. Overexpression of both *GmERS1-1-CEND* and *AtERS1-CEND* can significantly inhibit the growth of soybean hair roots, and** **subcellular localization of all transformations. (A)** Both *GmERS1-1-CEND* and *AtERS1-CEND* were overexpressed in soybean using the soybean root transient expression system, and the free GmERS1-1-CEND and AtERS1-CEND could significantly inhibit the elongation of soybean roots. (1) overexpressing *GmERS1-1* with *GFP* tags located at the N-terminus; (2) overexpression of *GmERS1-1* with *GFP* tag located at the C-terminus; (3) overexpression of the free *GmERS1-1* N-terminus (*GmERS1-1-NEND*) labeled with GFP; (4) overexpression of free *GmERS1-1* C-terminus (*GmERS1-1-CEND*) with *GFP*; (5) overexpression of *AtERS1* with *GFP* tag located at the N-terminus; (6) overexpression of *AtERS1* with *GFP* tag located at the C-terminus; (7) overexpression of *AtERS1* N-terminus (*AtERS1-NEND*) with *GFP*; (8) overexpression of *AtERS1* C-terminus (*AtERS1-CEND*) with *GFP*; (9) control group transformed with empty vectors. The upper row was no treated with ethylene, while the lower row was treated with ethylene. **(B)** Root length statistics results. Data statistics conducted ANOVA-DUNCAN's multiple range tests at *P*<0.05, error bars report SE of the mean, n=3. **(C)** The subcellular localization results of all transformations in soybean hairy roots untreated with ethylene on the left and treated with ethylene on the right, bar=5 μ M.

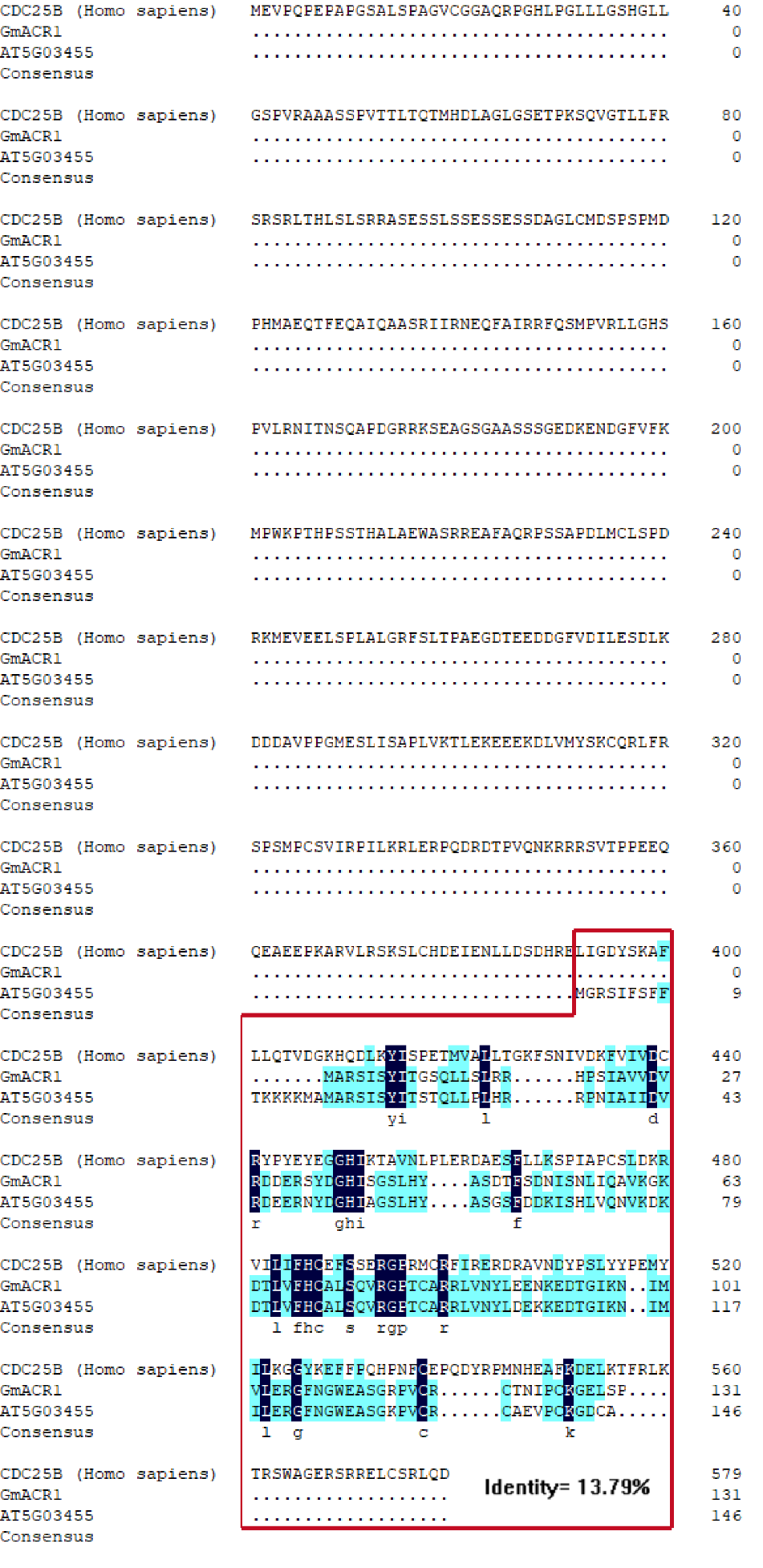
**Figure S12**

**Figure S12. Multiple sequence alignment of human CDC25B, soybean GmACR1, and Arabidopsis AtACR1 (AT5G03455).** Full-length protein sequences of human CDC25B (*Homo sapiens*), soybean GmACR1, and Arabidopsis AtACR1 were aligned. Dots represent sequence gaps introduced for optimal alignment; the red box demarcates the only conserved region shared across the three proteins, with identical amino acid residues shaded in dark blue and similar residues highlighted in light cyan. The overall pairwise sequence identity between human CDC25B and soybean GmACR1 is indicated as 13.79% at the bottom of the alignment. The consensus row summarizes conserved amino acid signatures within the highlighted homologous domain.

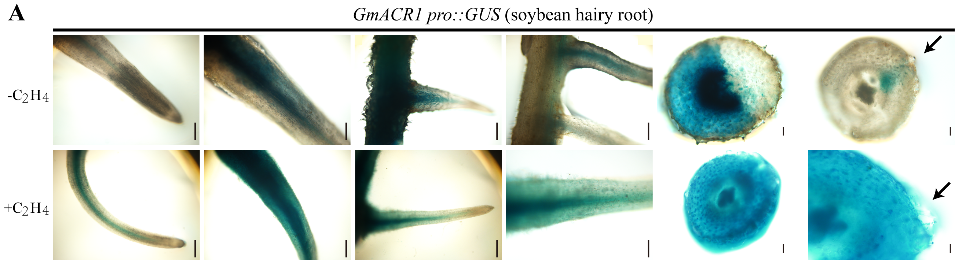
**Figure S13**

**Figure S13. Activity analysis of GmACR1 promoter. (A)** Using the soybean hairy root transformation system to obtain transgenic *GmACR1 pro::GUS* hairy roots, observe them before and after treatment, and observe the treatment conditions as shown in Figure 1.5. The black arrow points to the lateral root primordium.

**Figure S1**
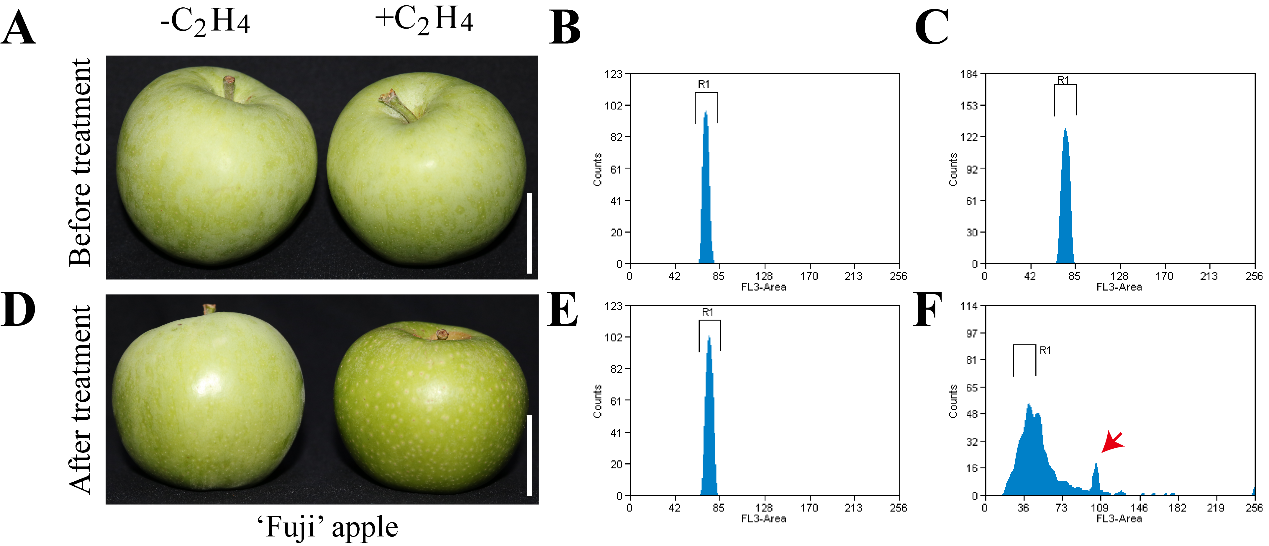
**4**

**Figure S14. Ethylene induces cell cycle arrest in ‘Fuji’ apple fruit cells.** **(A, D)** Representative photographs of ‘Fuji’ apple fruit before **(A)** and after **(D)** mock (−C₂H₄) or ethylene (+C₂H₄) treatment. Bar=5 cm. **(B, C)** Flow cytometry profiles of fruit cells isolated from −C₂H₄ **(B)** and +C₂H₄ **(C)** apples prior to treatment, showing nearly identical cell cycle distribution gated in region R1. **(E, F)** Post-treatment flow cytometry histograms for −C₂H₄ **(E)** and +C₂H₄ **(F)** fruit cells. The red arrow in **(F)** marks an emerging secondary peak indicative of G2/M-phase blockage in ethylene-exposed apple cells, which is absent in the mock control. FL3-Area, fluorescence signal intensity for nuclear DNA content; Counts, number of detected nuclei.

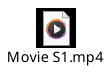

**Movie S1. Temporal observation of fluorescence signals in root tip cell nuclei of *GmERS1-1 pro::GmERS1-1-GFP* overexpression strains.** **It is a supplement to Figure 3H.**

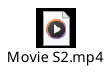

**Movie S2. Temporal observation of fluorescence signals in root tip cell nuclei of *GmERS1-1 pro::GmERS1-1-NEND-GFP* overexpression strains.** **It is a supplement to Figure 3H.**

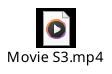

**Movie S3. Temporal observation of fluorescence signals in root tip cell nuclei of *GmERS1-1 pro::GmERS1-1-CEND-GFP* overexpression strains.** **It is a supplement to Figure 3H.**

**Table S1. Oligonucleotide sused in this study.**

| Primers used for ***GmERS1-1*** gene cloning | ATGATGGAGTCCTGTGATTGTATAG | TTAGAAACTTCTTTGATTGCGTCTA |
| --- | --- | --- |
| Primers used for ***GmERS1-1* promoter** cloning | TTGCTTTTTTATTCTTTTGT | CTAACCCCCGTATGA |
| Primers **used for promoter activity analysis of *GmERS1-1* in *Arabidopsis thaliana* and soybean hairy root (1305)** | ccatgattacgaattcTTGCTTTTTTATTCT | ctcagatctaccatggCTAACCCCCGTATGA |
| Primers **used for** construction of *GmERS1-1* overexpressing lines (3301) | GGACTCTTGACCATGATGATGGAGTCCTGT | ATTCGAGCTGGTCACCTTAGAAACTTCTTTG |
| Primers **used for** construction of *GmERS1-1-GFP overexpression in soybean hairy root* (1302) | GGGACTCTTGACCATGATGATGGAGTCCTGT | TCAGATCTACCCATGGGAAACTTCTTTG |
| Primers **used for detection of *gmers1-1 #2* mutant** | ATGATGGAGTCCTGTGATTGTATAG | TTAGAAACTTCTTTGATTGCGTCTA |
| Primers **used for subcellular localization (16318)** | TATCTCTAGAGGATCCATGATGGAGTCCTGTGATTGTATAG | TGCTCACCATGGATCCGAAACTTCTTTGATTGCGTCTA |
| nuclear export signal (NES) | GAACGAGCTTGCTCTTAAGTTGGCTGGACTTGATATTAACAAGACTGGAGGATAGCCCGGGCTGCA | GCCCGGGCTATCCTCCAGTCTTGTTAATATCAAGTCCAGCCAACTTAAGAGCAAGCTCGTTCTGCA |
| SALK-LBb1.3 | ATTTTGCCGATTTCGGAAC | |
| SALK-LP | CGCAAATCATTAAGAGCCAAG | |
| SALK-RP | GACACAGGGTGTGGAATTCAC | |
| 3301-1F | GGACTCTTGACCATGatgatggagtcctgt | |
| 3301-1R | ATTCGAGCTGGTCACCttagaaacttctttg | |
| 3301-1-1-C-F | GGACTCTTGACCATGATGAAGAGGCGCGAATCG | |
| 3301-1-1-C-R | ATTCGAGCTGGTCACCTTAGAAACTTCTTTGATT | |
| 3301-1-1-N-F | GGACTCTTGACCATGATGATGGAGTCCTGT | |
| 3301-1-1-N-R | ATTCGAGCTGGTCACCTTAGACACTCAACAGATC | |
| 1302/1-1F | gggactcttgaccatgatgatggagtcctgt | |
| 1302/1-1R | tcagatctaccatggcttagaaacttctttg | |
| 1302/1-1NF | gggactcttgaccatg**ATGATGGAGTCCTGT** | |
| 1302/1-1NR | tcagatctaccatggGACACTCAACAGATC | |
| 1302/1-1CF | gggactcttgaccatg**atgatggagtcctgt** | |
| 1302/1-1CR | tcagatctaccatgg**GAAACTTCTTTGATT** | |
| AtETR1-F | ATGGAAGTCTGCAATTGTATTGAAC | |
| AtETR1-R | TTACATGCCCTCGTACAGTACCCGG | |
| AtERS1-F | ATGGAGTCATGCGATTGTTTTGAGA | |
| AtERS1-R | TCACCAGTTCCACGGTCTGGTTTGT | |
| AtETR2-F | ATGGTTAAAGAAATAGCTTCTTGGT | |
| AtETR2-R | TTAGAGAAGTTGGTCAGCTTGCAAC | |
| AtERS2-F | ATGTTAAAGACATTGTTAGTCCAAT | |
| AtERS2-R | TCAGTGGCTAGTAGACGGAGGAGTT | |
| AtEIN4-F | ATGTTAAGATCTTTAGGTCTTGGAT | |
| AtEIN4-R | TCACTCGCTCGCGGTCTGCAAAGCC | |
| 16G-AtETR1-F | TATCTCTAGAGGATCCATGGAAGTCTGCAATTG | |
| 16G-AtETR1-R | TGCTCACCATGGATCCCATGCCCTCGTACAGTA | |
| 16G-AtERS1-F | TATCTCTAGAGGATCCATGGAGTCATGCGATTG | |
| 16G-AtERS1-R | TGCTCACCATGGATCCCCAGTTCCACGGTCTGG | |
| 16G-AtETR2-F | TATCTCTAGAGGATCCATGGTTAAAGAAATAGC | |
| 16G-AtETR2-R | TGCTCACCATGGATCCGAGAAGTTGGTCAGCTT | |
| 16G-AtERS2-F | TATCTCTAGAGGATCCATGTTAAAGACATTGTT | |
| 16G-AtERS2-R | TGCTCACCATGGATCCGTGGCTAGTAGACGGAG | |
| 16G-AtEIN4-F | TATCTCTAGAGGATCCATGTTAAGATCTTTAGG | |
| 16G-AtEIN4-R | TGCTCACCATGGATCCCTCGCTCGCGGTCTGCA | |
| 1-1jianceA-F | gaagttcatttcatttggagagaaca | |
| 1-1jianceA-R | ccaacttgcacatgataggtta | |
| 1-1jianceBar-F | gacgggcaggaccggacg | |
| 1-1jianceBar-R | agcccagaacgacgcccg | |
| GmACR1-F | ATGTCAGACATGGATCCGTCCAAAA | |
| Gm ACR1-R | TCAGTTGAAATCACCAAAACTGGCA | |
| pCold-GmACR1-F | TACCCTCGAGGGATCCATGGCTCGCAGTATA | |
| pCold-GmACR1-R | GCTTGAATTCGGATCCTTAAGGAGACAACTC | |
| pGEX-4T-GmACR1-F | TggATCCCCggAATTCATGGCTCGCAGTATA | |
| pGEX-4T-GmACR1-R | GTCgACCCgggAATTCTTAAGGAGACAACTC | |
| pET-28a-GmACR1-F | TCGCGGATCCGAATTCATGGCTCGCAGTATA | |
| pET-28a-GmACR1-R | GACGGAGCTCGAATTCTTAAGGAGACAACTC | |
| Q-ERS1-1F | CGCCACTAGTGGATCCATGATGGAGTCCTGT | |
| Q-ERS1-1R | TACTATCGATGGATCCgaaacttctttgatt | |
| Q-ACR1F | CGCCACTAGTGGATCCATGGCTCGCAGTATA | |
| Q-ACR1R | TACTATCGATGGATCCaggagacaactctcc | |
| 1-1-F no-ter | ATgATggAgTCCTgTgATTg | |
| 1-1-R no-ter | gAAACTTCTTTgATTgCgtct | |
| 1-F no-ter | ATGGCTCGCAGTATATCGTA | |
| 1-R no-ter | aggagacaactctcctttgc | |
| 0800-1pro-F | cagcccgggggatccTAATTCAATGAAAAA | |
| 0800-1pro-R | AGAACTAGTGGATCCTATCTCATACCTCCT | |
| 1pro-f | TAATTCAATGAAAAAATATTTAAAA | |
| 1pro-r | TATCTCATACCTCCTCTCCACTCTT | |
| 0800-1pro-F | cagcccgggggatccTAATTCAATGAAAAA | |
| 0800-1pro-R | AGAACTAGTGGATCCTATCTCATACCTCCT | |
| hz1pro/1305-F | ccatgattacgaattcTAATTCAATGAAAAA | |
| hz1pro/1305-R | ctcagatctaccatggTATCTCATACCTCCT | |
| qGmActin-F | CGGTGGTTCTATCTTGGCATC | |
| qGmActin-R | GTCTTTCGCTTCAATAACCCTA | |
| Q6-ERS1-1-KM-F | CGCCACTAGTGGATCCATGATGGAGTCCTGT | |
| Q6-ERS1-1-KM-R | TACTATCGATGGATCCGACACTCAACAGATC | |
| Q6-ERS1-1-YL-F | CGCCACTAGTGGATCCATGAAGAGGCGCGAATCG | |
| Q6-ERS1-1-YL-R | TACTATCGATGGATCCGAAACTTCTTTGATT | |
| GFP-ERS1-1-KM-F | TATCTCTAGAGGATCCATGATGGAGTCCTGT | |
| GFP-ERS1-1-KM-R | TGCTCACCATGGATCCGACACTCAACAGATC | |
| GFP-ERS1-1-YL-F | TATCTCTAGAGGATCCATGAAGAGGCGCGAATCG | |
| GFP-ERS1-1-YL-R | TGCTCACCATGGATCCGAAACTTCTTTGATT | |
| BT3N-ERS1-1-KM-F | CCTCGGCCCCATGGATGATGGAGTCCTGT | |
| BT3N-ERS1-1-KM-R | AGCTACTTACCATGGTTAGACACTCAACAGATC | |
| BT3N-ERS1-1-YL-F | CCTCGGCCCCATGGATGAAGAGGCGCGAATCG | |
| BT3N-ERS1-1-YL-R | AGCTACTTACCATGGTTAGAAACTTCTTTGATT | |
| ppr3n-hz1-f | GCCATTACGGCCCGGGAAATGGCTCGCAGTATA | |
| ppr3n-hz1-r | CATGTTTTTTCCCGGGTTAAGGAGACAACTC | |
| pBridge-hz1-F | AGAAAGGTGGCGGCCATGGCTCGCAGTATA | |
| pBridge-hz1-R | GGCTAATGCGGCCGCTTAAGGAGACAACTC | |
| 1-1F | ATGATGGAGTCCTGTGATTG | |
| 1-1R | TTAGAAACTT CTTTGATTGC | |
| 1-1P+G-F | TTGCTTTTTTATTCTTTTGT | |

**Table S2. Antibodies and germplasm materials.**

| **REAGENT or RESOURCE SOURCE** | **SOURCE** | **IDENTIFIER** |
| --- | --- | --- |
| **Antibodies** | | |
| Anti-GmERS1-1-C’ | ABclonal | WG-03923 |
| FITC-conjugated Goat anti-Mouse IgG (H+L) | ABclonal | AS001 |
| HRP-conjugated Mouse anti GST-Tag mAb | ABclonal | AE027 |
| HRP-conjugated Mouse anti His-Tag mAb | ABclonal | AE028 |
| ACR1 Rabbit mAb | ABclonal | A97580 |
| FITC-conjugated Goat anti-Rabbit IgG (H+L) | ABclonal | AS011 |
| HRP-conjugated Mouse anti GFP-Tag mAb | ABclonal | AE030 |
| HRP-conjugated Mouse anti DDDDK-Tag mAb | ABclonal | AE024 |
| **Experimental models: Organisms/strains** | **SOURCE** | **IDENTIFIER** |
| Arabidopsis Col-0 | *15* | N/A |
| *Arabidopsis ers1* (*aters1*) | TAIR | SALK.046728c |
| *Arabidopsis ET-free-1* | *16* | N/A |
| Soybean *ers1* (*gmers1-1* #2) | *17* | NJAU0116 |
| Soybean *ctr1* (*gmctr1*) | *17* | NJAU0144 |
| Soybean *acr1* (*gmacr1*) | *17* | NJAU0298 |

**Table S3.The expression data of 11 ethylene receptors were downloaded from the SoyBase database (https://soybase.org).**

|  | Glyma.03G216700.1 | Glyma.03G251700.1 | Glyma.09G002600.1 | Glyma.10G167300.1 | Glyma.10G188500.1 | Glyma.12G241700.1 | Glyma.19G213300.1 | Glyma.19G249100.1 | Glyma.20G087000.1 | Glyma.20G202200.1 | Glyma.20G221800.1 |
| --- | --- | --- | --- | --- | --- | --- | --- | --- | --- | --- | --- |
| Flower | 10.1992 | 1.04157 | 27.0097 | 4.38072 | 26.0381 | 33.0222 | 17.104 | 0.848719 | 2.30151 | 7.29296 | 5.19453 |
| Flower.open | 9.856 | 0.514 | 0.562 | 3.308 | 3.453 | 3.131 | 5.674 | 0.326 | 3.399 | 2.591 | 1.811 |
| Flower.unopen | 14.693 | 0.465 | 0.208 | 6.393 | 1.252 | 0.926 | 6.29 | 0.44 | 2.99 | 1.132 | 2.164 |
| Lateral root.standard | 17.08 | 1.242 | 0.35 | 2.973 | 1.623 | 3.161 | 6.235 | 0.799 | 2.664 | 1.747 | 1.219 |
| Leaf.ammonia | 5.648 | 0.772 | 0.187 | 2.819 | 0.556 | 0.713 | 5.207 | 0.454 | 0.191 | 1.209 | 0.857 |
| Leaf.nitrate | 7.47 | 0.704 | 0.243 | 2.739 | 0.817 | 1.136 | 8.117 | 0.462 | 0.475 | 1.675 | 0.862 |
| Leaf.standard | 9.587 | 0.576 | 0.17 | 3.234 | 0.646 | 0.684 | 7.722 | 0.525 | 0.39 | 1.225 | 0.928 |
| Leaf.symbiotic condition | 4.916 | 1.361 | 0.215 | 3.708 | 0.672 | 0.692 | 6.137 | 0.61 | 0.126 | 1.587 | 0.958 |
| Leaf.urea | 6.205 | 0.818 | 0.215 | 4.345 | 0.65 | 0.8 | 6.401 | 0.483 | 0.107 | 1.212 | 1.193 |
| Leaves | 13.2176 | 3.55609 | 21.5191 | 7.28246 | 13.9888 | 17.224 | 44.5529 | 1.8035 | 0.807505 | 14.5666 | 3.52837 |
| Nodules | 20.9157 | 4.0636 | 9.79736 | 5.75544 | 11.5678 | 14.9859 | 17.3668 | 1.9633 | 3.67514 | 10.7006 | 6.67189 |
| Nodules.symbiotic condition | 4.29 | 0.092 | 0.245 | 0.655 | 4.607 | 0.5 | 2.308 | 0.077 | 2.79 | 10.655 | 0.678 |
| Pod | 28.7022 | 1.15112 | 5.72041 | 8.73108 | 7.34986 | 8.99694 | 14.1967 | 0.72173 | 5.37365 | 2.95827 | 9.37486 |
| Root | 22.5154 | 2.29986 | 6.2563 | 6.4167 | 6.39102 | 12.562 | 29.3782 | 5.86837 | 1.77706 | 5.08688 | 4.04082 |
| Root.ammonia | 9.798 | 0.346 | 0.686 | 1.598 | 11.243 | 1.554 | 3.719 | 0.389 | 8.184 | 13.294 | 0.928 |
| Root Hairs | 26.3653 | 3.31932 | 6.77976 | 4.1116 | 11.3193 | 17.47 | 23.5193 | 1.7796 | 5.54842 | 8.92199 | 3.59583 |
| Root.nitrate | 13.016 | 0.19 | 0.284 | 0.978 | 13.841 | 0.841 | 3.508 | 0.178 | 9.039 | 26.214 | 0.659 |
| Root.standard | 8.862 | 0.079 | 0.136 | 2.19 | 5.495 | 0.642 | 2.917 | 0.158 | 3.004 | 11.574 | 0.864 |
| Root.symbiotic condition | 5.076 | 0.174 | 0.523 | 0.885 | 9.348 | 0.948 | 2.161 | 0.234 | 6.001 | 14.635 | 0.755 |
| Root tip.standard | 19.505 | 1.338 | 0.289 | 2.872 | 1.894 | 3.387 | 6.759 | 0.749 | 2.808 | 2.011 | 1.269 |
| Root.urea | 9.514 | 0.284 | 0.582 | 0.806 | 12.414 | 1.509 | 3.595 | 0.133 | 10.498 | 22.541 | 0.545 |
| Seed | 17.458 | 4.14936 | 4.19122 | 4.32382 | 16.4294 | 7.11407 | 4.83927 | 0.876043 | 7.79843 | 8.10198 | 4.2571 |
| Shoot Apical Meristem | 18.3948 | 2.9113 | 4.61798 | 1.61637 | 5.65156 | 5.48331 | 10.4099 | 1.02068 | 2.30661 | 4.70117 | 1.79088 |
| Shoot tip.standard | 13.334 | 0.533 | 0.175 | 0.988 | 1.533 | 0.694 | 4.164 | 0.338 | 4.167 | 1.539 | 0.68 |
| Stem | 19.2255 | 2.99658 | 12.0626 | 5.25682 | 12.255 | 12.9645 | 15.9114 | 1.59151 | 10.3114 | 5.43315 | 6.30021 |
| Stem.standard | 14.353 | 0.323 | 0.16 | 1.978 | 1.711 | 0.671 | 5.431 | 0.188 | 2.407 | 1.596 | 1.084 |

**Table S4. Member information of soybean ethylene receptor.**

| Chr01 | | | 56831624 | | |
| --- | --- | --- | --- | --- | --- |
| Chr02 | | | 48577505 | | |
| Chr03 | | | 45779781 | | |
| Chr04 | | | 52389146 | | |
| Chr05 | | | 42234498 | | |
| Chr06 | | | 51416486 | | |
| Chr07 | | | 44630646 | | |
| Chr08 | | | 47837940 | | |
| Chr09 | | | 50189764 | | |
| Chr10 | | | 51566898 | | |
| Chr11 | | | 34766867 | | |
| Chr12 | | | 40091314 | | |
| Chr13 | | | 45874162 | | |
| Chr14 | | | 49042192 | | |
| Chr15 | | | 51756343 | | |
| Chr16 | | | 37887014 | | |
| Chr17 | | | 41641366 | | |
| Chr18 | | | 58018742 | | |
| Chr19 | | | 50746916 | | |
| Chr20 | | | 47904181 | | |
| 1 | Glyma.03G216700.1 | Chr03:42073661..42078243 reverse | 42073661 | 42078243 | Chr03 |
| 2 | Glyma.03G251700.1 | Chr03:44736493..44741844 reverse | 44736493 | 44736493 | Chr03 |
| 3 | Glyma.09G002600.1 | Chr09:219108..224886 forward | 219108 | 224886 | Chr09 |
| 4 | Glyma.10G167300.1 | Chr10:40161567..40166587 reverse | 40161567 | 40166587 | Chr10 |
| 5 | Glyma.10G188500.1 | Chr10:42156133..42158963 reverse | 42156133 | 42158963 | Chr10 |
| 6 | Glyma.12G241700.1 | Chr12:39999320..40004986 reverse | 39999320 | 40004986 | Chr12 |
| 7 | Glyma.19G213300.1 | Chr19:46666253..46671097 reverse | 46666253 | 46671097 | Chr19 |
| 8 | Glyma.19G249100.1 | Chr19:49516179..49520236 reverse | 49516179 | 49520236 | Chr19 |
| 9 | Glyma.20G087000.1 | Chr20:32527435..32530216 | 32527435 | 32530216 | Chr20 |
| 10 | Glyma.20G202200.1 | Chr20:43919183..43923660 | 43919183 | 43923660 | Chr20 |
| 11 | Glyma.20G221800.1 | Chr20:45676789..45681839 | 45676789 | 45681839 | Chr20 |

**Table S5. Member information of *Arabidopsis thaliana* ethylene receptor.**

| Chr1 | | | 29105111 | |
| --- | --- | --- | --- | --- |
| Chr2 | | | 19646945 | |
| Chr3 | | | 23172617 | |
| Chr4 | | | 17549867 | |
| Chr5 | | | 25953409 | |
| AtETR1 | AT1G66340.1 | Chr1 | 24734162 | 24737756 |
| AtERS1 | AT2G40940.1 | Chr2 | 17084356 | 17087297 |
| AtETR2 | AT3G23150.1 | Chr3 | 8254663 | 8257735 |
| AtERS2 | AT1G04310.1 | Chr1 | 1154764 | 1157711 |
| AtEIN4 | AT3G04580.1 | Chr3 | 1235283 | 1238475 |

**References**

1. K. Tamura, G. Stecher, S. Kumar, Mol. Biol. Evol. 38, 3022–3027 (2021).

2. W. Ning et al., Nucleic Acids Res. 50, W405–W411 (2022).

3. X. Zhang et al., Plant Physiol. 174, 2487–2500 (2017).

5. M. Zhang et al., Plant Biotechnol. J. 20, 1435–1437 (2022).

6. D. G. Lee et al., Anal. Methods 2, 1577–1583 (2010).

7. X. Lu et al., Sci. China Life Sci. 62, 1070–1077 (2019).

8. H. Qiao et al., Science 338, 390–393 (2012).

9. A. Khojasteh et al., Plant Physiol. Biochem. 141, 133–141 (2019).

10. J. Doležel, J. Bartoš, Ann. Bot. 95, 99–110 (2005).

11. M. Sauer et al., Nat. Protoc. 1, 98–103 (2006).

12. K. Chen et al., Plant J. 107, 740–759 (2021).

13. Q. Lin et al., Nat. Commun. 3, 752 (2012).

14. J. Abramson et al., Nature 630, 493–500 (2024).

15. Y. Yan et al., Nat. Protoc. 15, 1–24 (2020).

16. K. Chen et al., Front. Plant Sci. 11, 555794 (2020).

17. W. Li et al., Mol. Plant 15, 354–362 (2022).

18. M. Zhang et al., Plant Biotechnol. J. 20, 1435–1437 (2022).

**Protein sequences of 5 ethylene receptors in Arabidopsis and 11 ethylene receptors in soybean**

>Glyma.03G216700.1

MMESCDCIDTQYPPDELLVKYQYISDVLIALAYFSIPVELIYFVQKSAFFPYRWVLMQFGAFIVLCGATHFINLWTFSPHSKSVAVVMTIAKVSCAIVSCATALMLVHIIPDLLSVKRRESILKNKAEELDREMGLILTQEETGRHVRMLTHEIRSTLDRHTILKTTLVELGRTLGLEECALWMPSRSGLNLQLSHTLTYHVQVGSTVQTNNPIVNEVFNSPQAMRIPPTCPLARIRPLVGRYVPPEVVAVRVPLLNLSNFQINDWPDISAKSYAIMVLILPTDSVRKWRDHELELVDVVADQVAVALSHAAILEESMRARDQLMEQNVALDLARREAEMAIHARNDFLAVMNHEMRTPMHAIIALSSLLLETELTPEQRVMIETVLKSSNVLATLINDVLDLSRLEDGSLELEMGKFNLHGVLGEIVELIKPIASVKKLPITLILSPDLPTHAIGDEKRLTQTLLNVVGNAVKFTKEGYVSVRVSVAKPESSQDWRPPEFYPASSDGHFYIRVQVKDSGCGILPQDIPHLFTKFAQSRSGPARPSSGAGLGLAICKRFVNLMGGHIWIESEGLDKGSTATFIVKLGICGNPDPSDHQAANRSQAYSGSGGLARFKPFITDENDTGFSTRRNQRSF

>Glyma.03G251700.1

MERGLLLLLFLLLVMVLSVCGNDVEYSQCNCDEEGLWSIHNVLVCQKVSDFFIAIAYFSIPLELLYFVSCSNVPFKLVFLQFIAFIVLCGLTHLLNAYTYYGPHSFQLFLSLTVAKFLTALVSCATAISFPTLIPLLLKIKVRELFLRQNVLELGQEVGMMKKQKEASWHVRMLTCEIRKSLDKHTILYITLVELSKALDLHNCAVWMPDEDRREMHLTHELKPNSTRIFHNSIPISDPDVLDIKKSQGVWILRPDSALGAASSGGGGSGDSGAVAAIRMPILHVSNFKGGTPEFVETSYGVLVLVLPNSDSRAWTSHEMEIVKVVADQVAVALSHASVLEESQLMSQKLAEQNRALQQAQKNAMMARKARSSFEKVMSHGMRRPMHSILGLLSMFQEDNIRPEQKIVIDSILKVSNALSRLINDVMEIAANDNGSFQLEMKPFHLHSMMREASCTAKCLCIYKGFGLEVDVDKSLPDLVIGDEARTFQVILHMIGYLLNIYDKGNLIFQVYLKSDSGDRDDRSFGLWRSSMQNEYVHIKFNFQINGISSQSDESVSTRNYTGRRHYNNEPKEGLSFSMCKTLVQMMQGNIWISTNSLGLAQGMTLLLKFQIGSSHGRFTLAPTDFSNSQFRGLKVVLADDDDVNRTVTKKLLEKLGCQVTAVSSGFECLGAISGSGNSFKIILLDLHMPEMDGFEVARRIRKFQSHNWPLIIAFTASAEEHIKERCLQVGMNGLIRKPILLREIADELGTVLQRAGTNKFRCGQSKW

>Glyma.09G002600.1

MESCNCIDPQVPADDLLMKYQYISDFFIALAYFSIPLELIYFVKKSAVFPYRWVLVQFGAFIVLCGATHLINLWTFRIHSRTVAVVMTTAKVLTAVVSCATALMLVHIIPDLLSVKTRELFLKNKAAELDREMGLIRTQEETGRHVRMLTHEIRSTLDRHTILKTTLVELGRTLALEECALWMPTRTGLELQLSYTLRQQNPVGYTVPIHLPVINQVFSSNRAVKISPNCPVARLRPYAGKYMPGAVVAVRVPLLHLSNFQIYDWPEVSTRSYALMVLMLPSDSARQWHVHELELVEVVADQVAVALSHAAILEESMRARDQLMEQNVALDLARREAETAIRARNDFLAVMNHEMRTPMHAVIALSSLLQETDLTAEQRLMVETILKSSNLLATLINDVLDLSRLEDGSLQLEATTFNLHSLFREVLNLIKPVASVKKLSLTSHIASDLPMYAIGDEKRLMQTILNVVGNAVKFSKEGCISISAFVAKPESFRDARIPDFLPVLSDNHFYLRVQVKDSGSGINPQDIPKIFTKFAQNQSLTTRNPAGSGLGLAICRRFVNLMEGHIWVESEGIGKGCTVTFIVKLGIPDRSNEFKLPFVPKVPGNHGSTNFAGLKVLVTDDNGVSRTVTKGLLMHLGCDVTTASSSEECLRVVSLEHEVVFMDVCAGLDGYELAIRIHEKFTKHQDRPLIVALTGNTKKVTKENCMRVGMDGLILKPVSVDKMRGVLSELLERRVLFETV

>Glyma.10G008500.1

MSQEGCLLIFLICSSQGPLSANDQYTIYIVGTVKVLVSMLPIFFFSPLILCFVCLLCYCQLMQGNIWLVPNAQGFPQVMALFLRFQLRRSIAVSISEPGENSEPSNSNSFFRGLQVLLADNDDVNRAVTQKLLQKLGCVVTSVSSGLECLSVIGPAGSSFQVILLDLHMPELDGFEVATRIRKFRSRNWPVIVALTASTEDLWERCMQIGMNGVIRKPVLLHGIASELRRILLQGNSAM

>Glyma.10G167300.1

MEITPPLLHHLWFLLLLLLLCYLVLCASATDVEFDNCNCDDGEGIWSIHSILVGQKVSDFFIAVAYFSIPIELLYFVSRSNVPFKLLFLQFIAFIVLCGMTHLLNAYSYHGPPSFQLLLSLTVAKFLTALVSCATALTLPPLIPLLLKVKVRELFLRQNVMELGQEVGMMKKQKEASWHVRMLTREIRKSLDKHNILYTTLVELSKALDLHNCAVWMPNEDRREMHLTHELKTNSAKNFQNSIPVNDPDVLEIRKTKGVKILGPESALGAASSGGSVELGAVAAIRMPLLHVSNFKGGTPELVETCYAILVLVLPSSSTRVWTYHEMEIVEVVADQVAVALSHASVLEESQLMRQKLEERNRALQQAKKNAMMASQARKSFQKVMSHGMRRPMHSVLGMLSLFQEDNLRSEQKIIGDTMLKVGHVLSSLINDVMEISENEKGGFRLEMKPFLLHSMVREAASIAKCLCVYEGFGFEIDVQKSLPETVMGDEARTFQVILHMIGYLLNMNDKGTLNFRVFLESDGGDRDDKNIGIWRSSNQNEYVHIKFDFQITESSQSDEAISTIHYTGRRQYYNNEPKGGLSFSMCKKLVQMMQGNIWISPNSLGLVHGMTLLLKFQIGPSLEKSIFAPKDYSSSQFRGLKVVLAEDDGVNRTVTKKLLEKLGCQVIAVSSGFECLSAVSGAGNSFRIILLDLHMPEMDGFELAKRIRKFHSRSWPLIIALITSAEEHVREKCLLAGMNGLIQKPIVLHQIADELRTVLQRAGEKL

>Glyma.10G188500.1

MLKAVASGLVITSILICVSANDNGFPRCNCDDEASFWTIESILECQRVGDFLIAVAYFSIPIELLYFISCSNVPFKWVLIQFIAFIVLCGLTHLLNGWTYGPHTFQLMVALTVSKILTALVSCATAITLITLIPLLLKVKVREFMLKKKTWDLGREVDHIMRQKEAAMHVRMLTQEIRKSLDRHTILYTTLVELSKTLGLQNCAVWMPNVDKTEMNLTHELNGRNFNLTIPISDPDVVRIKGSDDVNILSSDSALAVGSRGVSGEAGPVAAIRMPMLRVCNFKGGTPELRQACYAILVLILPTGDNQEPRSWSNQELEIIKVVADQVAVALSHAAILEESQLMREKLEEQNRALQQAKRNALMASQARNAFQKVMSDGMRRPMHSILGLLSMIQDDNLKNEQKLIVDAMLRTSNVLSNLINDAMDNSTKDEGRFSLEIRSFGLHSMLKEAACLSKCMCVYKGFGFMVEVEKSLPDNVMGDERRVFQKTGSQGRSDKGWTTWRPSSSSGDVNIRFEIGINSSDSEVGSSISSGFGGRKYSSDRVGGRLSFSICKRVVQLMQGNIWLVPCNHGFPQSMTLLLRFQLRPSITIAISDPGEGSEHTDSNSMLRSLQVLLVDNDDVNRAVTQRLLQKLGCVVTSVASGFECLTVIGPAGSSIQVILLDLHMPDIDGFEVATRIRKFRSGNRPMIVALTASAEEDLWDRCMQVGINGVIRKPVLLHGIASELRRILMQGNIVL

>Glyma.12G241700.1

MESCNCIDPQVPADDLLMKYQYISDFFIALAYFSIPLELIYFVKKSAVFPYRWVLVQFGAFIVLCGATHLINLWTFRIHSRTVAVVMTTAKVLTAVVSCATALMLVHIIPDLLSVKTRELFLKNKAAELDREMGLIRTQEETGRHVRMLTHEIRSTLDRHTILKTTLVELGRTLALEECALWMPTRTGLELQLSYTLRQQNPVGYTVPIHLPVINQVFSSNRAVKISPNCPVARLRPYAGKYMPGAVVAVRVPLLHLSNFQIYDWPEVSTRSYALMVLMLPSDSARQWHVHELELVEVVADQVAVALSHAAILEESMRARDQLIEQNVALDLARREAETAIRARNDFLAVMNHEMRTPMHAVIALSSLLQETDLTAEQRLMVETILKSSNLLATLINDVLDLSRLEDGSLQLEAATFNLHSLFREVLNLIKPVASVKKLSLTSHVASDLPMYAIGDEKRLMQTILNVVGNAVKFSKEGCISITAFVAKPESFRDARIPDFLPVPSDNHFYLRVQVKDSGSGINPQDIPKLFTKFAQNQSLTRNPAGSGLGLAICRRFVNLMEGHIWVESEGIGKGCTVTFIVKLGIPDRSNEFKLPFVPKVPGNHGSTNFAGLKVLVMDDNGVSRTVTKGLLMHLGCDVTTASSSEECLRVVSLEHEVVFMDVCTGLDGYELAVRIHEKFTKHQDRPLIVALTGNTKKVTKENCMRVGMDGLILKPVSVDKMRGVLSELLERRVLFESV

>Glyma.17G181400.1

MERGLILLLFLLLVMVLSVYSNEVEYSHIHNVLVCQKVSDFFIAIAYFSIPLELLYFVSCSNVPFKLVFLQFIAFIVLCGLNHLLNAYTYYGRHSFQLFLSITIAKFLTALVSCATAISFPTLIPLLLKIKVRELFFWQNVLELG

>Glyma.18G069300.1

MMQGNIWISPNTLGLVQGMTLLLKFQIGTSLEKSIFAPKDCSSSQFRGLKVVLAEDDGVNRTVTKKLLEKLGCQVIAVSSGFECLSDISGAGNSFRIILLDLHMPEMDGFEVAKKIQKFHSHSWHLIIALIASAQEHLREKCLLVGMNGLIQKPIVLHQIANELRTILQRAGEKL

>Glyma.19G213300.1

MMESCDCIDTQYPPDELLVKYQYISDVLIALAYFSIPVELIYFVQKSAFFPYRWVLMQFGAFIVLCGATHFINLWTFSPHSKAVAVVMTIAKVSCAIVSCATALMLVHIIPDLLSVKTRELFLKNKAEELDREMGLILTQEETGRHVRMLTHEIRSTLDRHTILKTTLVELGRTLGLEECALWMPSRSGLNLQLSHTLTYHVQVGSTVQTNNPIVNEVFNSPRAMRIPPTCPLARIRPLVGRYVPPEVVAVRVPLLNLSNFQINDWPDMSAKSYAIMVLILPTDSVRKWRDHELELVDVVADQVAVALSHAAILEESMRARDQLLEQNVALDLARQEAEMAIHARNDFLAVMNHEMRTPMHAIIALSSLLLETELTPEQRVMIETVLKSSNVLATLINDVLDLSRLEDGSLELEKGKFNLHGVLGEIVELIKPIASVKKLPITLILSPDLPTHAIGDEKRLTQTLLNVVGNAVKFTKEGYVSIRVSVAKPESLQDWRPPEFYPASSDGHFYIRVQVKDSGCGIPPQEIPHLFTKFAQSRSGPARPSSGAGLGLAICKRFVNLMGGHIWIESEGPDKGSTATFIIKLEICGNPDPSDHQAANRSQAYSGSGGLARFKPFIKDEDDTGFSTRRNQRSF

>Glyma.19G249100.1

MERGLLLLLFLLLVMVLSVYANDVEYSQCNCDEEGLWSIHSVLVCQKVSDFFIAIAYFSIPLELLYFVSCSNVPFKLVFLQFIAFIVLCGLTHLLNAYTYYGPHSFQLFLSLTVAKFLTALVSCATAISFPTLIPLLLKIKVRELFLRQNVLELGQEVGMMKKQKEASWHVRMLTCEIRKSLDKHTILYTTLVELSKALDLHNCAVWMPDEDRREMHLTHELKPSSARSFHNSIAISDPDVLDIKKSQGVWILRPDSALGAASSGGGSGDSGAVAAIRLPILHVSNFKGGTPELVETSYGVLVLVLPNSNSRAWTSHEMEIVEVVADQVAVALSHASVLEESQLMSQKLAEQNRALQQAQKNAMMARKARSSFEKVMSHGMRRPMHSILGLLSMFQEDNIRPEQKIVIDSILKVSNALSRLINDVMEIAENDNGSFQLEMKPFHLHSMMREASCTAKCLCIYKGFGLEVDVDKSLPDLVIGDEARTFQVILHMIGYLLNIYDKGTLTFQVYLESDSGDKDDRSFGIWRSSIQNEYVHIKFNFQINGISFHSDESVSTRNYTGRNHCNNELKEGLSFSMCKTLVQMMQGNIWISTNSLGLAQGMTLLLKFQIGSSHGRFILAPKEFSNSQFRGLKVVLADDDDVNRTVTKKLLEKLGCQVTAVSSGFECLGAISASGNSFKIIMLDLHMPEMDGFEVARRIRKFQSHNFFLCSLLLLQVQKNTSRRDVYRWE

>Glyma.20G087000.1

MLKAVSFELLLICLLLLRTCVCATTDNGFPRCNCDDESSLWTIETILECQRIGDFLIAVAYFSIPIELLYFVSCSNFPFKWVLFQFIAFIVLCGMTHLLNGWTYGPHTFQLMVALTVFKILTALVSCATTITLLTLIPMLLKVKVRELMLKKKTWDLGREVGIIMKQKEAAMHVRMLTQEIRKSLDRHKILYTTLVELSKTLGLQNCAVWMPNVEKTEMNLTHELNGRNVNCSIPITNPDVVRIKGSDEVNIIDSDSILATASSGVYGGAGPVAAIRMPMLQVCNFKGGTPELRQTCYAILVLTLPSAEPRSWGAQELEIIKVVADQVAVALSHASILEESQLMREKLEEQNRALQMEKMNTMMASQARASFQKVTSNGMRRPMHSILGLLSMMQDDNLKSEQKLIVNSMLRTSTVLSNLINDAMDYSTRDDGRFPLEMKPFGLHAMVKEAACLAKCIFEIGINNGDPELESSVPSGQLAGTDRTSDKVEERLSFSICKRIIQLMQGNIWLVPNAQGFPQVMALFLRFQLWRSIAVSNSEPGENSETSNSNSFFRGLQVLLADNDDVNRAVTQKLLQKLGCVVTSVSSGFECLNVIGPAGSSFQVILLDLHMPELDGFEVATRIPKFRSRNWPVIVALTASTDDLWERCMQIGMNGVIRKPVLLHGIASELRRIILQGNSVM

>Glyma.20G202200.1

MLKAVASGLVITSILICVSANDNGFPRCNCDDEASLWTIESILECQRVGDFLIAVAYFSIPIELLYFISCSNVPFKWVLIQFIAFIVLCGLTHLLNGWTYGPHTFQLMVALTVSKILTALVSCATAITLITLIPLLLKVKVREFMLKKKTWDLGREVDHIMRQKEAAMHVRMLTQEIRKSLDRHTILYTTLVELSKTLGLQNCAVWMPNVDKTEMNLTHELNGRNFNLTIRITDPDVVRIKGSDGVNILSSDSALAVGSRGVSGEAGPVAAIRMPMLRVCNFKGGTPELRQACYAILVLILPSGDNQEPRSWSNQELEIIKVVADQVAVALSHAAILEESQLMREKLEEQNRALQQAKRNALMASQARNAFQKVMSDGMRRPMHSILGLLSMIQDDKLKSEQKLIVDAMLRTSNVLSNLINDAMDNSTKDEGRFPLEIRSFGLHSMLKEAACLSKCMCVYKGFGFMVEVEKCLPDNVMGDERRVFQVILHMVGNLLEHNHGGGILVYRVFAETGSQGRSDKGWTTWRPSSSSGDVNIRFEIGINSSDSEVGSSISSGFGGRKYSSDRVGGRLSFSICKRVVQLMQGNIWLVPCNHGFPQSMTLLLRFQLRPSISIAISDPGEGSERTDSNSMLRNLQVLLVENDDVNRAVTQRLLQKLGCVVTPVASGFECLTVIGPAGCSIQVILLDLHMPDLDGFEVATRIRKFRSGNQPMIVALTASAEEDLWERCMQVGINGVIRKPVLLHGIASELRRILMQGNNVL

>Glyma.20G221800.1

MEITPPPLLLLFLLLLFFLCYLVLCASATDVDFDNCNCDDGEGIWSIHSILVGQKVSDFFIAVAYFSIPIELLYFVSRSNVPFKLLFLQFIAFIVLCGMTHLLNAYSYHGPPSFQLLLSLTVAKFLTALVSCATALTLPPLIPLLLKIKVRELFLRQNVMELGQEVGMMKKQKEASWHVRMLTREIRKSLDKHNILYTTLVELSKALDLHNCAVWMPNEDRREMHLTHELKTNSAKNFQNSIPVNDPDVLEIRKTKGVKILRPDSAIGAASSGGSAELGAVAAIRMPLLHVSNFKGGTPQLVETCYAILVLVLPSSSTRVWTYHEMEIVEVVADQVAVALSHASVLEESQQMRQKLEERNRALQQAKKNAMMASQARKSFQEVMSHGMRRPMHSILGMLSLFQEDNLRSEQKIIGDTMLKVGHVLSSLINDVMEISENEKGGFRLEMKPFLLHSMMREAASIAKCLCVYEGFGFEIDVQKSLPETVMGDEARTFQVILHMIGYLLNMNDKGTLNFRVFLESDGGDRDDKNIGIWRSSSQNEYVHIKFDFQITESSQSDKAISTIHYSSRRQYYNNEPKEGLSFSMCKKLVQMMQGNIWISPNSLGLVQGMTLLLKFQIGPSLGKSIFAPKDYSSSQFRGLKVLLAEDDGVNRTVTKKLLEKLGCQVIAVSSGFECLSAISGAGNSFRIILLDLHMPEMEGFEVAKRIRKFHSRSWPLIIALIASAEEHVREKCLLAGMNGLIQKPIVLHQIANELRTVLQRAGEKL

>AT1G66340.1

MEVCNCIEPQWPADELLMKYQYISDFFIAIAYFSIPLELIYFVKKSAVFPYRWVLVQFGAFIVLCGATHLINLWTFTTHSRTVALVMTTAKVLTAVVSCATALMLVHIIPDLLSVKTRELFLKNKAAELDREMGLIRTQEETGRHVRMLTHEIRSTLDRHTILKTTLVELGRTLALEECALWMPTRTGLELQLSYTLRHQHPVEYTVPIQLPVINQVFGTSRAVKISPNSPVARLRPVSGKYMLGEVVAVRVPLLHLSNFQINDWPELSTKRYALMVLMLPSDSARQWHVHELELVEVVADQVAVALSHAAILEESMRARDLLMEQNVALDLARREAETAIRARNDFLAVMNHEMRTPMHAIIALSSLLQETELTPEQRLMVETILKSSNLLATLMNDVLDLSRLEDGSLQLELGTFNLHTLFREVLNLIKPIAVVKKLPITLNLAPDLPEFVVGDEKRLMQIILNIVGNAVKFSKQGSISVTALVTKSDTRAADFFVVPTGSHFYLRVKVKDSGAGINPQDIPKIFTKFAQTQSLATRSSGGSGLGLAISKRFVNLMEGNIWIESDGLGKGCTAIFDVKLGISERSNESKQSGIPKVPAIPRHSNFTGLKVLVMDENGVSRMVTKGLLVHLGCEVTTVSSNEECLRVVSHEHKVVFMDVCMPGVENYQIALRIHEKFTKQRHQRPLLVALSGNTDKSTKEKCMSFGLDGVLLKPVSLDNIRDVLSDLLEPRVLYEGM

>AT2G40940.1

MESCDCFETHVNQDDLLVKYQYISDALIALAYFSIPLELIYFVQKSAFFPYKWVLMQFGAFIILCGATHFINLWMFFMHSKAVAIVMTIAKVSCAVVSCATALMLVHIIPDLLSVKNRELFLKKKADELDREMGLILTQEETGRHVRMLTHGIRRTLDRHTILRTTLVELGKTLCLEECALWMPSQSGLYLQLSHTLSHKIQVGSSVPINLPIINELFNSAQAMHIPHSCPLAKIGPPVGRYSPPEVVSVRVPLLHLSNFQGSDWSDLSGKGYAIMVLILPTDGARKWRDHELELVENVADQVAVALSHAAILEESMHARDQLMEQNFALDKARQEAEMAVHARNDFLAVMNHEMRTPMHAIISLSSLLLETELSPEQRVMIETILKSSNLVATLISDVLDLSRLEDGSLLLENEPFSLQAIFEEVISLIKPIASVKKLSTNLILSADLPTYAIGDEKRLMQTILNIMGNAVKFTKEGYISIIASIMKPESLQELPSPEFFPVLSDSHFYLCVQVKDTGCGIHTQDIPLLFTKFVQPRTGTQRNHSGGGLGLALCKRFVGLMGGYMWIESEGLEKGCTASFIIRLGICNGPSSSSGSMALHLAAKSQTRPWNW

>AT3G23150.1

MVKEIASWLLILSMVVFVSPVLAINGGGYPRCNCEDEGNSFWSTENILETQRVSDFLIAVAYFSIPIELLYFVSCSNVPFKWVLFEFIAFIVLCGMTHLLHGWTYSAHPFRLMMAFTVFKMLTALVSCATAITLITLIPLLLKVKVREFMLKKKAHELGREVGLILIKKETGFHVRMLTQEIRKSLDRHTILYTTLVELSKTLGLQNCAVWMPNDGGTEMDLTHELRGRGGYGGCSVSMEDLDVVRIRESDEVNVLSVDSSIARASGGGGDVSEIGAVAAIRMPMLRVSDFNGELSYAILVCVLPGGTPRDWTYQEIEIVKVVADQVTVALDHAAVLEESQLMREKLAEQNRALQMAKRDALRASQARNAFQKTMSEGMRRPMHSILGLLSMIQDEKLSDEQKMIVDTMVKTGNVMSNLVGDSMDVPDGRFGTEMKPFSLHRTIHEAACMARCLCLCNGIRFLVDAEKSLPDNVVGDERRVFQVILHIVGSLVKPRKRQEGSSLMFKVLKERGSLDRSDHRWAAWRSPASSADGDVYIRFEMNVENDDSSSQSFASVSSRDQEVGDVRFSGGYGLGQDLSFGVCKKVVQLIHGNISVVPGSDGSPETMSLLLRFRRRPSISVHGSSESPAPDHHAHPHSNSLLRGLQVLLVDTNDSNRAVTRKLLEKLGCDVTAVSSGFDCLTAIAPGSSSPSTSFQVVVLDLQMAEMDGYEVAMRIRSRSWPLIVATTVSLDEEMWDKCAQIGINGVVRKPVVLRAMESELRRVLLQADQLL

>AT1G04310.1

MLKTLLVQWLVFFFFFLIGSVVTAAEDDGSLSLCNCDDEDSLFSYETILNSQKVGDFLIAIAYFSIPIELVYFVSRTNVPSPYNWVVCEFIAFIVLCGMTHLLAGFTYGPHWPWVMTAVTVFKMLTGIVSFLTALSLVTLLPLLLKAKVREFMLSKKTRELDREVGIIMKQTETSLHVRMLTTKIRTSLDRHTILYTTLVELSKTLGLKNCAVWIPNEIKTEMNLTHELRPRIDDENENEHFGGYAGFSIPISESDVVRIKRSEEVNMLSPGSVLASVTSRGKSGPTVGIRVPMLRVCNFKGGTPEAIHMCYAILVCVLPLRQPQAWTYQELEIVKVVADQVAVAISHAVILEESQLMREKLAEQNRALQVARENALRANQAKAAFEQMMSDAMRCPVRSILGLLPLILQDGKLPENQTVIVDAMRRTSELLVQLVNNAGDINNGTIRAAETHYFSLHSVVKESACVARCLCMANGFGFSAEVYRALPDYVVGDDRKVFQAILHMLGVLMNRKIKGNVTFWVFPESGNSDVSERKDIQEAVWRHCYSKEYMEVRFGFEVTAEGEESSSSSSGSNLEEEEENPSLNACQNIVKYMQGNIRVVEDGLGLVKSVSVVFRFQLRRSMMSRGGGYSGETFRTSTPPSTSH

>AT3G04580.1

MLRSLGLGLLLFALLALVSGDNDYVSCNCDDEGFLSVHTILECQRVSDLLIAIAYFSIPLELLYFISFSNVPFKWVLVQFIAFIVLCGMTHLLNAWTYYGPHSFQLMLWLTIFKFLTALVSCATAITLLTLIPLLLKWKVRELYLKQNVLELNEEVGLMKRQKEMSVQVRMLTREIRKSLDKHMILRTTLVELSKILDLQNSAVWMPNENRTEMHLTHELRANPMRSFRVIPINDPDVVQVRETKVVTILRKNSVLAVESSGCGGSEEFGPVAAIRMPMLHGLNFKGGTPEFVDTPYAIMVLVLPSANSRVWTDKEIEIAEVVADQVAVAISHASVLEESQLMREKLGIQNRALLRAKQNAMMASQARNTCQKVMSHGMRRPMHTILGLLSMFQSESMSLDQKIIVDALMKTSTVLSALINDVIDISPKDNGKSALEVKRFQLHSLIREAACVAKCLSVYKGYGFEMDVQTRLPNLVVGDEKRTFQLVMYMLGYILDMTDGGKTVTFRVICEGTGTSQDKSKRETGMWKSHMSDDSLGVKFEVEINEIQNPPLDGSAMAMRHIPNRRYHSNGIKEGLSLGMCRKLAQMMQGNIWISPKSHGQTQSMQLVLRFQTRPSIRRSILAGNAPELQHPNSNSILRGLRITLADDDDVNRTVTKRLLEKLGCEVTAVSSGFECLNALSNVEMSYRVVILDLQMPEMDGFEVAMKIRKFCGHHWPLIIALTASTEDHVRERCLQMGMNGMIQKPVLLHVMASELRRALQTASE

**Sequence related to *GmERS1-1***

>*GmERS1-1* promoter

TTGCTTTTTTATTCTTTTGTTTTTCTCCAATTTCACTTTATTTTTTTATTTCTTATTTCAACTTAACCAAGGATATCAAAAAGTCTAGTTCAATTGATTGAATAGAGTATATAAGTATTATAAATTTATTGGTACTGCATTTAATTTATATTAAAAAAAAAAACTTAACCAAGGAGTAAAACCAGTAGAAATTATAAGGACGGATGCATTATTATCATTACTGTCTTTTTTATTTATTTTGTTTAATGGGAGATTCCTTTTGGGACTTACAAGAAAAAGAGCAAGATGAAATACAAGAATAAGAAAGCGAAAAAAAGAGTAGAATGCTATGAATATGAATGATCAAATATGGATTATTCATGTACTATTATTTAACTCAGTTTTTTTACAGGATTATTCAACTTAGTTAGCTGTAACCACTATGAATTTATTTAGATTTTACTTAAAATAATTCAAAAGTATTTTTATTCTTAAAATGACAGTTTTTATTTATTTTTTATTTTGTGCATTAAAATTTGTAATTACATATTAGAATTCAAGAAATATTTTTTTTTCAGATTCGGAAAGAATACTTTCCAAACATTACTTCATACATAATGTACTTATTTGTAATTTCTTTTAAGAGTTTAATATTAATAATGTAAAATTTTATATTATCATTTAATTATAAATCATCATTTAAATTATTTTAAAATAATTATTTTAAAATTTTAATAAATGTATCATACGTAATATGTTATGACTGAACGATTATGTTAAATTTTCTATATTATTAATGTATATTTTTTCTTAAGTCTTTTTCAAGATATTTTTTATGACATATTCAAGATATATACTTTTTATTTGAAACACAAGTATCTCTTTCATCATAATTTTATTCAAGATGAATATAATTATTATAAATAACAAAAGTTTATATTCAAATAATTAACATAATTTGAATATGAAATCAGTGAATAAACACCAATTAATTCTTAATCGAAACATGTGGTTTGATAAACAAACAAGGCAATTGGAAATTTGGACCGGATGCACGGGATTTGACACCCATTAGGAAAAGTTAACACACATCGATATAGTGGTCCAAATTCAAAGAAACTTCACAATGCCACGGCCAAGAAAGTTGAATTACGAACACCAAAGCAGAATTAAGGACTTCAAATATATTTTTTTGAAAGGGTGAATCGAATCTGAAATTTATTTTACACGTAGCCAGAAAAAGTGAAGCTGAGTAGGCCCAGGCCACCGTCCAATGAATTCCCATTTCCTAACATAAAAAACCACAGCCACATTAGCTTCTAGGCAAAAATGAGAAAATATAAAATAAAAAAAGGTCCCAGCCAATCCGAAAATGATGTGGCACGTGTGTAAAATGAGAGAAGCATGGAAAAAAGGGAACGCCCGAGTCAGAGTAGGCGTTTGCCATGCTTGTCCGTGAGGGAGAGAGCTTAAGGATTAATGAATGCTTTTTTAACTCACTCTCACACTCTTTCTTCCTTTCTTCTTCTTCTTCTTCTTCCTGTCTTCACCTAACCAAATGGGCGCCTCCACCACCTGATCCACCTCTGTTTCTTCTTCCTTTCGTAATAATCACCTCTCTCTCTCTCTCTCTCTCTCTCAATCATTGGATCCATTCATCACGATTTCTCTATGCTGTTTCATAAACACTCTTAGCTTTGTTTTTATTCGCTGTATTTACTTCCGCAGATTTATATTGACCGTTATCCTATGTAGATCAAATACTCCGTTATTTTCATTTCCGCTAGATTTTATCAGCATTCGAGCGATTGTTGGATACGCGTAATGATTTTTTTTTTTTTGGATTGGTCTCTTAACTCGTGTTTTTAGGGTTTATTTATCCAAGTTAATGGTGGTTGACTTGTTGCTATTGTTCACTTGTTGAAGTTTGAATACTATTTTGGGAAATGATGGAATACAAAATGCATAGAGAATAATTTTTTTGATACCTGTTATTACAGATCCGTTCATACGGGGGTTAG

>Glyma.03G216700|Gm03:45379664..45384968 reverse|upstream=2000|downstream=0

GTTTTTTTTTTTTTTCTATTCAATGCTCATAAAAATTTATGGTGTGTGGAAAAAATTATGGAGCATGGATATACACAATAAAACTAACCATTGATTTAATATTGATAAATCAACAACGCATCGAGAATTTAACTCGGTTAGACGAATATGATAAATGAATTTTATAAACTTTTTTAGTACTGTGTTCAATTTTTACGAGTAAAAAAAAAAAATCAACAATAATAAATATAAGTAATAATAAATTATATTTCTTAACCAAGTGACTTACTATTCAAATTTTAACCTTAAATATGAAATAAGAAAAAAATATTTATTACCTATAGACTCACGATTTTTTATGAACATTAATCATTGTTATTGACAAAGACTATTTCATATTTATATCATAATAAAAAAATATTTAAAAATATGTTACTTAGATTTATGATCCTAACTTATAAAATAGGTGACTCAATTATTTTATAAATATTAGTTAGGACAAAATTTGCAAAATTAAGTAAGTTTAGAGTCAAAATGAGATCTTTTAAGGGCACAATTCACAAAAATTTGAAAATTAAGAGACAAGAAGTCCGTTGAAATTTTTTAATGAACTAATTTAATCGAGGATTAAATTAATTGATATTTATAATGTCAAAAGTTATTTTGTCATTTATTTATTTTAGGGATCAATTTGTCCTTTCATTAGAATTTCAATGACAAAATTGATTGATAATATAAGACTATTTTATTCTAAAAAAAGATTTGGAAGCAATTTATCTGTCTATTATAATTTCATGAACTAAATTTGTATTTCATTTTTCTTTTTTTCTTATTTTGCTTTTTTATTCTTTTGTTTTTCTCCAATTTCACTTTATTTTTTTATTTCTTATTTCAACTTAACCAAGGATATCAAAAAGTCTAGTTCAATTGATTGAATAGAGTATATAAGTATTATAAATTTATTGGTACTGCATTTAATTTATATTAAAAAAAAAAACTTAACCAAGGAGTAAAACCAGTAGAAATTATAAGGACGGATGCATTATTATCATTACTGTCTTTTTTATTTATTTTGTTTAATGGGAGATTCCTTTTGGGACTTACAAGAAAAAGAGCAAGATGAAATACAAGAATAAGAAAGCGAAAAAAAGAGTAGAATGCTATGAATATGAATGATCAAATATGGATTATTCATGTACTATTATTTAACTCAGTTTTTTTACAGGATTATTCAACTTAGTTAGCTGTAACCACTATGAATTTATTTAGATTTTACTTAAAATAATTCAAAAGTATTTTTATTCTTAAAATGACAGTTTTTATTTATTTTTTATTTTGTGCATTAAAATTTGTAATTACATATTAGAATTCAAGAAATATTTTTTTTTCAGATTCGGAAAGAATACTTTCCAAACATTACTTCATACATAATGTACTTATTTGTAATTTCTTTTAAGAGTTTAATATTAATAATGTAAAATTTTATATTATCATTTAATTATAAATCATCATTTAAATTATTTTAAAATAATTATTTTAAAATTTTAATAAATGTATCATACGTAATATGTTATGACTGAACGATTATGTTAAATTTTCTATATTATTAATGTATATTTTTTCTTAAGTCTTTTTCAAGATATTTTTTATGACATATTCAAGATATATACTTTTTATTTGAAACACAAGTATCTCTTTCATCATAATTTTATTCAAGATGAATATAATTATTATAAATAACAAAAGTTTATATTCAAATAATTAACATAATTTGAATATGAAATCAGTGAATAAACACCAATTAATTCTTAATCGAAACATGTGGTTTGATAAACAAACAAGGCAATTGGAAATTTGGACCGGATGCACGGGATTTGACACCCATTAGGAAAAGTTAACACACATCGATATAGTGGTCCAAATTCAAAGAAACTTCACAATGCCACGGCCAAGAAAGTTGAATTACGAACACCAAAGCAGAATTAAGGACTTCAAATATATTTTTTTGAAAGGGTGAATCGAATCTGAAATTTATTTTACACGTAGCCAGAAAAAGTGAAGCTGAGTAGGCCCAGGCCACCGTCCAATGAATTCCCATTTCCTAACATAAAAAACCACAGCCACATTAGCTTCTAGGCAAAAATGAGAAAATATAAAATAAAAAAAGGTCCCAGCCAATCCGAAAATGATGTGGCACGTGTGTAAAATGAGAGAAGCATGGAAAAAAGGGAACGCCCGAGTCAGAGTAGGCGTTTGCCATGCTTGTCCGTGAGGGAGAGAGCTTAAGGATTAATGAATGCTTTTTTAACTCACTCTCACACTCTTTCTTCCTTTCTTCTTCTTCTTCTTCTTCCTGTCTTCACCTAACCAAATGGGCGCCTCCACCACCTGATCCACCTCTGTTTCTTCTTCCTTTCGTAATAATCACCTCTCTCTCTCTCTCTCTCTCTCTCAATCATTGGATCCATTCATCACGATTTCTCTATGCTGTTTCATAAACACTCTTAGCTTTGTTTTTATTCGCTGTATTTACTTCCGCAGATTTATATTGACCGTTATCCTATGTAGATCAAATACTCCGTTATTTTCATTTCCGCTAGATTTTATCAGCATTCGAGCGATTGTTGGATACGCGTAATGATTTTTTTTTTTTTGGATTGGTCTCTTAACTCGTGTTTTTAGGGTTTATTTATCCAAGTTAATGGTGGTTGACTTGTTGCTATTGTTCACTTGTTGAAGTTTGAATACTATTTTGGGAAATGATGGAATACAAAATGCATAGAGAATAATTTTTTTGATACCTGTTATTACAGATCCGTTCATACGGGGGTTAGATGATGGAGTCCTGTGATTGTATAGATACGCAGTATCCGCCGGATGAACTTCTTGTGAAGTATCAGTATATCTCGGATGTGCTAATTGCTCTTGCGTATTTTTCGATTCCGGTGGAGCTCATCTATTTTGTTCAGAAGTCTGCTTTCTTTCCGTATAGATGGGTGCTTATGCAGTTTGGTGCCTTTATTGTTCTCTGTGGAGCAACTCATTTCATAAACCTGTGGACATTCTCTCCGCACTCTAAGTCTGTTGCTGTTGTCATGACGATTGCCAAAGTTTCGTGTGCTATTGTGTCATGTGCGACGGCACTGATGCTTGTACACATTATTCCCGATCTGTTGAGTGTCAAGAGGCGCGAATCGATCCTCAAGAACAAGGCTGAAGAGCTTGACAGAGAGATGGGACTTATTCTTACTCAGGAAGAGACGGGAAGACATGTTAGAATGTTGACTCATGAAATTAGGAGCACACTTGACAGGCATACAATTTTAAAGACTACTCTTGTGGAGCTGGGGAGGACTTTGGGCTTGGAGGAGTGTGCATTATGGATGCCTTCGAGAAGTGGTCTGAATCTGCAACTTTCCCATACTTTAACCTATCATGTGCAAGTTGGGTCTACGGTGCAAACAAACAATCCTATTGTCAATGAAGTTTTCAACAGCCCTCAAGCTATGCGGATACCACCCACCTGTCCATTGGCCAGGATCAGACCTCTTGTGGGAAGATATGTGCCCCCTGAAGTTGTTGCTGTTCGGGTGCCGCTTTTGAATCTGTCAAATTTTCAAATTAACGATTGGCCTGATATTTCAGCAAAGAGCTATGCAATCATGGTTCTCATCCTCCCTACTGACAGTGTTAGAAAATGGCGAGACCATGAGTTGGAACTTGTTGATGTTGTTGCAGATCAGGTATTGGATTCATTTTGAGATTTCCTTTATCATACTAGTTGTCATTTTTCTTGTGCCATTCTTGCCTTTGATTGTATGTTCCCCATACCTTATTTTCATTATTGTGTTTTCCTTTACTCCTTATTAATTTGCTTAGATAGTCGTACTCATTGTCATTGGTAGTTCTGTTCTTACTGTCATATGGCACTTGTTTTGGAGAATGGAATATTAAGAGAATTGAGAATTGGGAAAGGTTATCAACCATTGGCCATTCGACCTTGCCTTAAACAATAACGATTCAGGATTTATTGGTTTTTCTCACCCACTGAGGAATAAAGAAGGTAAGTAAATGCTATTTGGTGAACTTGATCCCAAAAGCTAGCTAAGGCAGAGGCAAGATTAAAGGATGTAATTACAGTAATTAGAATATCTGTTATCACTGTATTTGTTGCTTTTCACTGTGTAATTAAAATGATTAAAATATCTGTGATCTTTGTATTAGTTGCTTTTCTCTATATAACCCAACCTCTACCATCTAGTCAATACCTAGATTTCATCATGCCTCTGATTTTCTTTTCCCTTAACGTGATATTAGGAATGGGATAGAGTTTGATACCATATAAAAATTGAGAATGGACTGGGCCTTACTCAACCAAAAGCTAGCTCAAGGGGGATCTTGGCTAGCTGATGTGGCCATGTGGGATCTTAACAAACCCCATTTGGGGATTGGATAAAGGTGATAACATCTAATAGCATCTTGACAGAAGAGATTGAGCTAATCTGATGTAGATCTATAATTTCAATCAGTAATTCAGTATGTGTTTGGATCTGCCTCATTCTCTACTTTAACTACCTGAAACAGTTCATTCATTTGTGTACTTTCTAGGAATATAATTTTCTTACTGGATTATATGACCATGTTAACCTGATGTTTGAAATTAGGCTTTATGGAAGTTGTCATACCAAAACTTTGCTGACACTTTCTGGTGTCAGTGGGATTTGTTTCTTTGTACACACCTCAAATTGGGTGACATGGCTGTGTTATTGTTAAAACATTTCAAAATACTTATATTTGGTGTCTATTCCTCTACTAGTTGAATTCATTTGTAGTTTCCAAACAGGTAGCAGTTGCCCTTTCACATGCTGCTATTTTGGAGGAGTCTATGCGTGCCCGTGATCAACTCATGGAGCAGAATGTTGCTTTAGATTTAGCCCGGCGAGAGGCAGAGATGGCAATTCATGCTCGCAATGATTTTCTTGCTGTCATGAATCATGAAATGAGGACACCAATGCATGCAATTATAGCATTGTCATCACTTCTCTTGGAGACTGAACTGACGCCAGAGCAAAGGGTTATGATAGAGACAGTGTTGAAGAGTAGTAATGTTTTGGCAACACTCATTAATGATGTTCTAGATCTTTCTCGACTTGAAGATGGTAGCCTCGAATTAGAAATGGGAAAATTCAACCTCCATGGAGTTTTGGGAGAGGTAATTGCTATAGGATTTTCTGCTTCATTATTCTGAAAAGGCAGTAACATAATTCTTTAGTCTTTACTATGAACTGCATTTATGCATTGTGTTCATGTTTTCTGTCTTGAATGCAGATTGTTGAACTGATAAAACCAATAGCATCCGTGAAAAAGTTACCTATCACCTTAATTCTGTCTCCTGATCTGCCTACTCATGCCATTGGTGATGAAAAGCGGCTTACACAAACTCTTTTGAATGTCGTGGGTAATGCTGTCAAATTCACTAAGGAGGGCTATGTTTCTGTAAGAGTATCAGTTGCAAAACCAGAATCTTCACAGGATTGGCGACCTCCAGAGTTTTATCCAGCATCTAGTGATGGCCATTTCTACATTCGAGTCCAGGTTATGACCGAGCCATTCAATATCTGAGCCTTAATTCTTTTAATTGTTAACTTATAAGATTTAGAAAAATACCATGTGTTTTATATCCAGGTAGTGATGTGATTGAACTCTTCAGTTATCTATAAATAATTGAGCCATCTGAAGCATCCATATGGCCATACCTAGATGCCTAGGAATTAAGCCAAAGACATTTGCCTTTTATTATTGTTGTCAGTATAACAACTGTTAGAAGTTAATCTAGATATTGCTTTTTCTTTATTATTATTATTGACCCAAGCATATCCCTTATGCAGGTTAAGGACTCTGGATGTGGGATTCTCCCACAAGATATTCCGCATCTCTTTACCAAGTTTGCCCAGTCTCGGAGTGGACCAGCTCGACCTAGCAGTGGTGCAGGTCTTGGGCTTGCCATTTGTAAAAGGTAGGGGGAAAGCATCTAATTCTGCTTTGAGTTTGCATTAGTTTCTCAATTCTCACTATTTACATTAATTTTTTGTTGATTACCTAGTGAGACACCGTTATTTATTGTATTTATACTAAATGGATGTCAATACTATAGCTTAAAGAATAGTTTCGTCTATTAAGCCTTTGACTCCTCCATAAAATTGATTCAGATTTGTAAACCTCATGGGAGGCCACATATGGATTGAGAGTGAGGGCCTTGACAAAGGAAGCACAGCTACATTTATAGTCAAACTTGGGATTTGTGGCAATCCAGATCCATCCGATCATCAAGCTGCAAACAGAAGCCAAGCATATAGTGGAAGTGGTGGCCTCGCTAGATTTAAACCCTTCATCACAGATGAAAACGACACTGGTTTTTCTACTAGACGCAATCAAAGAAGTTTCTAATATCCAAGACACCAAAATTGCATACCTCTCAATCACAGTCGACAAGATGGAGTAAACCTCCTACTTTCATTTTGCATGTGCTAAATTGTATAATTTGCAGTTTGCGGCTGTAGAATAGGATTACTATGTCAGGCATATTTCTAACAAACAGGTTTTATGCGAACTCCTGTTGTATTAACTAGAATACAATACAACAAAACAAAGTTGAGAGACGATCTGTTACTTAATTTGACTTGTAGTATATATTGTCAGTGGCTTTTGCCGTTTAGCGTTCATAAAAATGACAGTATTGGCATGAATCAGTTGTCATCCCTCCCCATGTTCCGTGATTCGTTAGATTTGGTGTCTGTCGTTTGTGCTAAACTTAAACTGTTTTGTGAAAGAAACATAGGATAAGTGGACGGATCAAGAAGCTTTGGTACAAAATTTGTGGGTTGGGAGCTTAATATTGGAACGGAATATATGCTTGTATAGCAAATATAGGGTGGGAAATTCACACAATAAACTCAACATAAAATATTACATGACATGAAACTAATATGATGTATATATTTATAAAATAATGTTACTTTAGTAATTTTATTTTTCATTCGCGTAGACTGCCTTTCCAAATTGTTATCATCATTTTATTTATGTTTAAGCCATGGTTGGTCAGTTGGGTTGCTTAAGCCATAATTTTTTTACATTTGAATGTTTATTTTATTTTTTAGTTTGGATATTAACGTCCAAATACCAAACCGAACACAAAGCCTATTTTGGACCCTATAAAACAAGTGCATGTAAACCTTAATCCATTTTCAAGCTTAACTTTGCGAGAAAATAATTCCTAAGAGTAGCATTTGTTTGTTTTTAGCCCTCTCGTTTCATCTGAACTAATT

>G.max Wm82.a6.v1|Glyma.03G216700.1 CDS

ATGATGGAGTCCTGTGATTGTATAGATACGCAGTATCCGCCGGATGAACTTCTTGTGAAGTATCAGTATATCTCGGATGTGCTAATTGCTCTTGCGTATTTTTCGATTCCGGTGGAGCTCATCTATTTTGTTCAGAAGTCTGCTTTCTTTCCGTATAGATGGGTGCTTATGCAGTTTGGTGCCTTTATTGTTCTCTGTGGAGCAACTCATTTCATAAACCTGTGGACATTCTCTCCGCACTCTAAGTCTGTTGCTGTTGTCATGACGATTGCCAAAGTTTCGTGTGCTATTGTGTCATGTGCGACGGCACTGATGCTTGTACACATTATTCCCGATCTGTTGAGTGTCAAGAGGCGCGAATCGATCCTCAAGAACAAGGCTGAAGAGCTTGACAGAGAGATGGGACTTATTCTTACTCAGGAAGAGACGGGAAGACATGTTAGAATGTTGACTCATGAAATTAGGAGCACACTTGACAGGCATACAATTTTAAAGACTACTCTTGTGGAGCTGGGGAGGACTTTGGGCTTGGAGGAGTGTGCATTATGGATGCCTTCGAGAAGTGGTCTGAATCTGCAACTTTCCCATACTTTAACCTATCATGTGCAAGTTGGGTCTACGGTGCAAACAAACAATCCTATTGTCAATGAAGTTTTCAACAGCCCTCAAGCTATGCGGATACCACCCACCTGTCCATTGGCCAGGATCAGACCTCTTGTGGGAAGATATGTGCCCCCTGAAGTTGTTGCTGTTCGGGTGCCGCTTTTGAATCTGTCAAATTTTCAAATTAACGATTGGCCTGATATTTCAGCAAAGAGCTATGCAATCATGGTTCTCATCCTCCCTACTGACAGTGTTAGAAAATGGCGAGACCATGAGTTGGAACTTGTTGATGTTGTTGCAGATCAGGTAGCAGTTGCCCTTTCACATGCTGCTATTTTGGAGGAGTCTATGCGTGCCCGTGATCAACTCATGGAGCAGAATGTTGCTTTAGATTTAGCCCGGCGAGAGGCAGAGATGGCAATTCATGCTCGCAATGATTTTCTTGCTGTCATGAATCATGAAATGAGGACACCAATGCATGCAATTATAGCATTGTCATCACTTCTCTTGGAGACTGAACTGACGCCAGAGCAAAGGGTTATGATAGAGACAGTGTTGAAGAGTAGTAATGTTTTGGCAACACTCATTAATGATGTTCTAGATCTTTCTCGACTTGAAGATGGTAGCCTCGAATTAGAAATGGGAAAATTCAACCTCCATGGAGTTTTGGGAGAGATTGTTGAACTGATAAAACCAATAGCATCCGTGAAAAAGTTACCTATCACCTTAATTCTGTCTCCTGATCTGCCTACTCATGCCATTGGTGATGAAAAGCGGCTTACACAAACTCTTTTGAATGTCGTGGGTAATGCTGTCAAATTCACTAAGGAGGGCTATGTTTCTGTAAGAGTATCAGTTGCAAAACCAGAATCTTCACAGGATTGGCGACCTCCAGAGTTTTATCCAGCATCTAGTGATGGCCATTTCTACATTCGAGTCCAGGTTAAGGACTCTGGATGTGGGATTCTCCCACAAGATATTCCGCATCTCTTTACCAAGTTTGCCCAGTCTCGGAGTGGACCAGCTCGACCTAGCAGTGGTGCAGGTCTTGGGCTTGCCATTTGTAAAAGATTTGTAAACCTCATGGGAGGCCACATATGGATTGAGAGTGAGGGCCTTGACAAAGGAAGCACAGCTACATTTATAGTCAAACTTGGGATTTGTGGCAATCCAGATCCATCCGATCATCAAGCTGCAAACAGAAGCCAAGCATATAGTGGAAGTGGTGGCCTCGCTAGATTTAAACCCTTCATCACAGATGAAAACGACACTGGTTTTTCTACTAGACGCAATCAAAGAAGTTTCTAA

>G.max Wm82.a6.v1|Glyma.03G216700.1.p

MMESCDCIDTQYPPDELLVKYQYISDVLIALAYFSIPVELIYFVQKSAFFPYRWVLMQFGAFIVLCGATHFINLWTFSPHSKSVAVVMTIAKVSCAIVSCATALMLVHIIPDLLSVKRRESILKNKAEELDREMGLILTQEETGRHVRMLTHEIRSTLDRHTILKTTLVELGRTLGLEECALWMPSRSGLNLQLSHTLTYHVQVGSTVQTNNPIVNEVFNSPQAMRIPPTCPLARIRPLVGRYVPPEVVAVRVPLLNLSNFQINDWPDISAKSYAIMVLILPTDSVRKWRDHELELVDVVADQVAVALSHAAILEESMRARDQLMEQNVALDLARREAEMAIHARNDFLAVMNHEMRTPMHAIIALSSLLLETELTPEQRVMIETVLKSSNVLATLINDVLDLSRLEDGSLELEMGKFNLHGVLGEIVELIKPIASVKKLPITLILSPDLPTHAIGDEKRLTQTLLNVVGNAVKFTKEGYVSVRVSVAKPESSQDWRPPEFYPASSDGHFYIRVQVKDSGCGILPQDIPHLFTKFAQSRSGPARPSSGAGLGLAICKRFVNLMGGHIWIESEGLDKGSTATFIVKLGICGNPDPSDHQAANRSQAYSGSGGLARFKPFITDENDTGFSTRRNQRSF

**>GmACR1**

ATGTCAGACATGGATCCGTCCAAAAAAGTGGCCGATAGATATCTGAAGCGTGAAGTCCTTGGTGAAGGTACCTATGGAGTTGTGTACAAAGCTATTGATACCCATACAGGGCAGACGGTTGCAATTAAGAAAATTCGGCTTGGCAAACAGAAAGAAGGGGTAAATTTTACAGCTCTTAGAGAAATTAAGCTGCTAAAAGAGCTCAAAGATCCAAACATTGTTGAATTGATTGATGCATTCCCACATAAAGGGAATTTGCATCTTGTTTTTGAATTCATGGAGACGGACCTAGAAGCCGTCATACGAGACCGAAATATTTTTCTTTCACCATCTGATACAAAATCCTACCTGCAAATGACACTAAAAGGTCTTGCTTATTGCCACAAGAAATGGGTTTTGCATAGGGACATGAAGCCAAATAATTTATTGATAGGATCTAATGGACAGCTGAAACTTGCAGATTTTGGTTTAGCACGGATGTTTGGAAGCCCAGATAGAAGGTTCACTCATCAGGTTTTTGCTCGGTGGTATAGAGCTCCTGAGCTATTATTTGGTGCCAAGCAATATGGTCCTGGAGTTGATGTTTGGGCTGCAGGCTGTATATTTGCTGAACTTCTTCTTCGTCGACCTTTTTTGCAGGGTACAAGTGACATTGATCAATTAGGAAAGATTTTTTCAGCATTTGGCACTCCAACGGCTTCTCAGTGGCCTGATATGGTATACCTGCCTGATTATGTCGAATACCAATATGTTCCTGCCCCCCCTCTGCGTTCTTTATTTCCGATGGTAACTGATGATGCTTTAGATCTGTTGTCAAAGATGTTTACATATGATCCAAAAGCTAGAATTTCAGTGCAGCAGGCACTAGAGCATAGGTACTTTTCTTCTGCTCCTCTGCCTTCGGATCCAGACAAGCTCCCTAGACCTGCTCCCAAGAGGGAATCTAGGACTTCTGATTTCAATTTACAGGAGGGTCCTACTGTATTATCACCTCCAAGGAAGTCTAGGAGAGTGATGCCAGGACGTGATGGGTTTGAAGGAAGTTCTCTGCAAGCAGATAAGGTTGATGCCAGTTTTGGTGATTTCAACTGA
